# Chronic brain functional ultrasound imaging in freely moving rodents performing cognitive tasks

**DOI:** 10.1101/2022.01.29.478327

**Authors:** Ahmed El Hady, Daniel Takahashi, Ruolan Sun, Oluwateniola Akinwale, Tyler Boyd-Meredith, Yisi Zhang, Adam S Charles, Carlos D Brody

## Abstract

Functional ultrasound imaging (fUS) is an emerging imaging technique that indirectly measures neural activity via changes in blood volume. To date it has not been used to image chronically during cognitive tasks in freely moving animals. Performing those experiments faces a number of exceptional challenges: performing large durable craniotomies with chronic implants, designing behavioural experiments matching the hemodynamic timescale, stabilizing the ultrasound probe during freely moving behavior, accurately assessing motion artifacts and validating that the animal can perform cognitive tasks at high performance while tethered. In this study, we provide validated solutions for those technical challenges. In addition, we present standardized step-by-step reproducible protocols, procedures and data processing pipelines that open up the opportunity to perform fUS in freely moving rodents performing complex cognitive tasks. Moreover, we present proof-of-concept analysis of brain dynamics during a decision making task. We obtain stable recordings from which we can robustly decode task variables from fUS data over multiple months. Moreover, we find that brain wide imaging through hemodynamic response is nonlinearly related to cognitive variables, such as task difficulty, as compared to sensory responses previously explored.

## 1.1 Introduction

A central aim of systems neuroscience is to identify how brain dynamics give rise to behavior. Cognitive behaviors, such as evidence accumulation-based decision making and working memory, involve concerted neural activity distributed across multiple brain areas (Pinto et al., 2019; Steinmetz et al., 2019). While there has been much effort in identifying and assessing the contributions of localized brain regions to specific tasks, fully understanding the neural circuitry underlying these behaviors requires identifying all areas involved and the dynamical interactions between them. Moreover, the brain-wide activity patterns may not follow the well-studied anatomical boundaries (Kaplan and Zimmer, 2020). Therefore, methodologies must be developed that can image large areas of the brain during cognitive behaviors to simultaneously uncover large-scale neuronal interactions and guide further, targeted recordings. Moreover, the computational models we use to interpret large-scale neural activity must be able to identify interacting brain regions regardless of anatomical constraints.

The development of such methods is further complicated by the desire of many neuroscientists to study naturalistic behaviors (Dennis et al., 2021; Krakauer et al., 2017; Egnor and Branson, 2016). Naturalistic behaviors unfold over multiple temporal and spatial scales, and are best studied in freely moving animals. Thus the developed methods must be stable and capable of recording brain activity chronically under freely moving conditions. Before transitioning to naturalistic behaviors, one first needs to standardize appropriate methods in more constrained tasks. The well-known two forced alternative choice (2-AFC) decision-making tasks, e.g., evidence accumulation, can serve as a behavioral test paradigm to develop such methods (Brunton, Botvinick, and Brody, 2013; Shadlen and Newsome, 2001; Brody and Hanks, 2016).

Historically, electric potential recordings have been the most accessible method to record the large-scale brain activity (Penfield and Jasper, 1954; Dimitriadis, Fransen, and Maris, 2014; Xie et al., 2017). Despite its portability and increasing sampling rates (Kaiju et al., 2017; Chiang et al., 2020), electrocorticogram (ECoG) has the remains constrained in its spatial resolution and penetrating depth. Similarly, cortex-wide optical imaging is also depth-constrained.

As an alternative, hemodynamic related imaging has been the dominant modality for wholebrain imaging due to its ability to penetrate deep into the brain. In particular, Functional Magnetic Resonance Imaging (fMRI) provides minimally invasive recordings of blood oxygenation (Ma et al., 2016; Hillman, 2014). The high level of penetration makes hemodynamic imaging ideal for capturing whole-brain networks. The longer physiological time-scale of hemodynamic activity, however, complicates subsequent analyses, especially during many systems neuroscience tasks that are designed to probe neuronal-scale activity and thus evolve on the subsecond time-scale. Moreover, the high magnetic field requirement for fMRI prohibits its use in freely moving animals.

One technology that is showing promise in recording the large brain areas (up to 15 mm depth by 16 mm lateral) needed to understand global brain dynamics is the recently developed functional ultrasound imaging (fUS). fUS is a Doppler-based technique that measures differentials in cerebral blood volume, which has been shown to be highly correlated to neural activity (Nunez-Elizalde et al., 2021; Aubier et al., 2021; Boido et al., 2019). Moreover, fUS can image deep structures simultaneously with the cortex.

fUS stands to play a pivotal role in the future of neuroscience by helping build an integrative platform for probing neural mechanisms underlying cognitive behaviors. To date, it has been used to probe the brain-wide dynamics in a variety of behaviors such as optokinetic reflex (Macé et al., 2018), auditory cortex mapping (Bimbard et al., 2018), hippocampal involvement in movement (Sieu et al., 2015; Bergel et al., 2020), innate behaviors (Sans-Dublanc et al., 2021), saccades guided task (Dizeux et al., 2019), visual cortex mapping (Blaize et al., 2020) and memory-guided behavior (Norman et al., 2021). In all previous studies, the animals were either head fixed or freely moving with the experimenter holding the wires. Ultimately, one would like to have a totally freely moving configuration where the animal can move around unencumbered by the tethering, allowing the animal to behave as naturally as possible and perform learned cognitive tasks.

Recent work has demonstrated the use of fUS in freely moving rats (Urban et al., 2015; Sieu et al., 2015). Nevertheless, in this freely moving configuration, fUS has been not been used in cognitive tasks, was not demonstrated in chronic recording over many months, and was not assessed for potential motion artifacts over extended timescales. Here we extend fUS in freely moving rats to months-long chronic imaging of cognitive behaviors, in particular demonstrating our methodology in chronic large-scale imaging of rats during an evidence accumulation task.

Using fUS robustly in freely moving rats thus maintains a number of challenges. Specifically, we require 1) stable recording over long periods of time 2) a pipeline that can correct motion artifacts, denoise the data, and extract meaningful components of brain-wide hemodynamic activity, and 3) matching the time-scale of behavior to the time-scale of hemodynamics. Moreover, we would like all the aforementioned to hold over a long chronic recordings, e.g., to enable imaging across learning. We present here a set of solutions that combine advances in surgical methods to enhance stability in the probe insertion, a computational toolkit for fUS pre-processing and basic data analysis, and behavioral training extensions to temporally elongate animal behavior. We demonstrate our methodology on multiple recordings spanning two months in the same freely moving rat performing an evidence accumulation task. We demonstrate robust, stable recordings as well as the extraction of task-related signals via our processing pipeline.

## 1.2 Results

### 1.2.1 fUS imaging

The imaging technique we use is the ultrafast ultrasound imaging developed in (Doppler, 1842; Macé et al., 2011; Montaldo et al., 2010; Urban et al., 2015) and used extensively in many applications (Tiran et al., 2017; Sieu et al., 2015; Demene et al., 2017; Takahashi et al., 2021). This method, based on power doppler (Montaldo et al., 2009; Rubin et al., 1995; Osmanski, Montaldo, and Tanter, 2015), relies on measured returns of a transmitted plane wave to determine the density of blood per voxel. The location of each voxel is computed by the time-to-return at an angle determined via beamforming. As the signal returns are noisy and have low dynamic range for blood moving parallel to the plane wave, multiple returns at different angles of plane wave transmission are used to generate each fUS video frame (Fig 1C).

**Figure 1:**
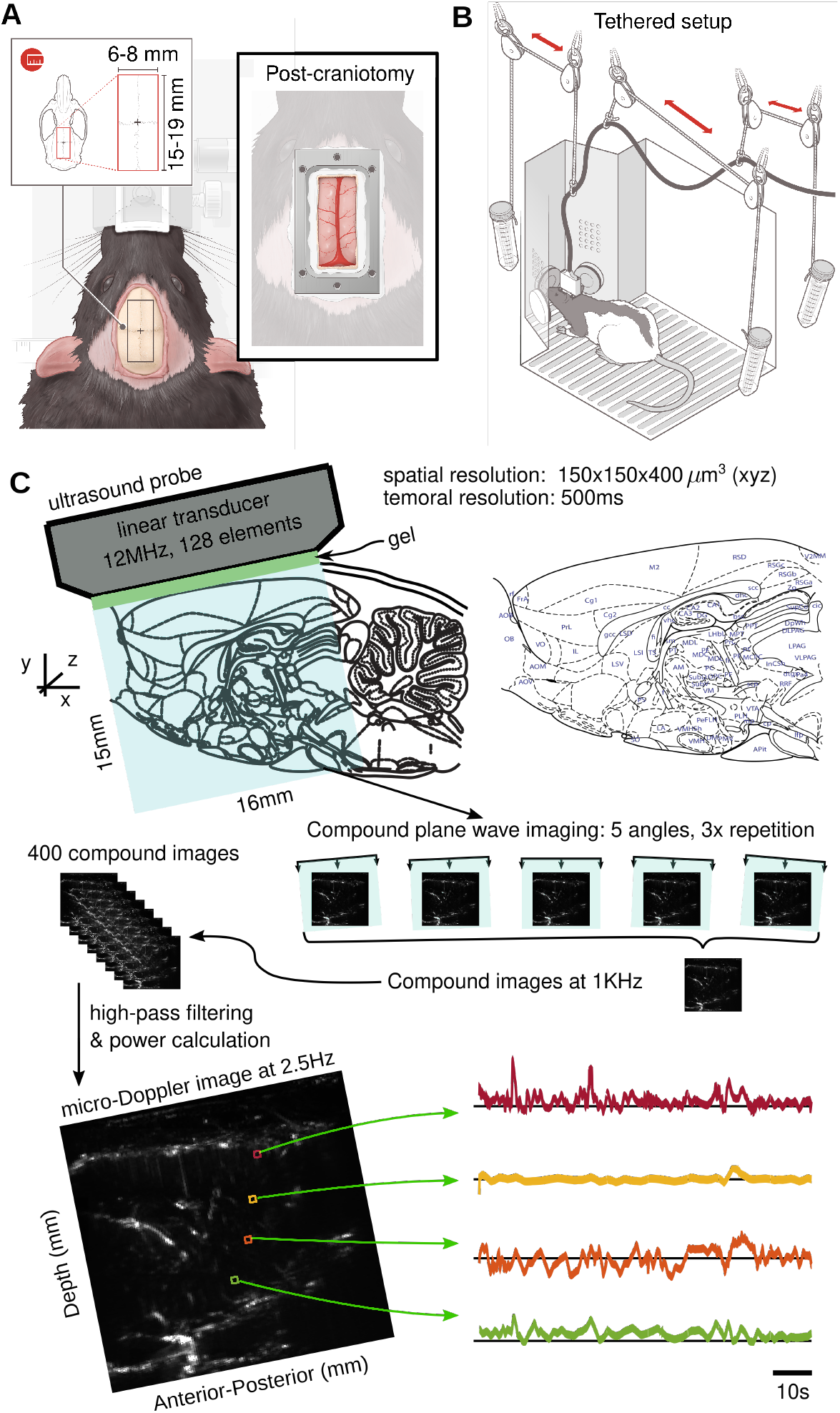
Functional ultrasound imaging in freely moving rats. A: Left; Rat skull dorsal surface. The red rectangle represents the area in which the craniotomy is done. Right; Surface view of the headplate positioned on the rat skull dorsal surface after the craniotomy (see Methods and appendix for surgical details). B: Post-craniotomy rat in the operant training chamber tethered with the fUS probe inside the probe holder. The weight of the probe (15 g) is low enough to enable comfortable behavior and movement for the rats. The pulley tethering system allows the animal to move naturally, without any manual intervention. C: The fUS probe we use images a 15 mm ×16 mm slice of the brain, including cortical and sub-cortical areas. The probe pulses at 12 MHz, and pulses are distributed across different probe angles to capture angle-agnostic activity, and repetitions to improve signal fidelity. The resulting compound image is produced at 1 KHz, every 400 images of which are filtered and integrated to produce a single *µ*Doppler image at 2.5 Hz. The bottom panel depicts an example image (Left) an fUS activity traces over time for four example pixels (Right).

As fUS imaging cannot be performed with an intact skull, due to the attenuation of the ultrasonic waves, we perform a craniotomy to access the brain (Fig. 1A). The probe is mounted using sturdy glue [AHMED ADD DETAILS] enabling the rat to freely move inside a behavioural operant chamber during imaging (Fig 1B). The extent of the large window provides access to 15 mmx16 mm images, e.g., from the +1 mm medio-lateral saggital plane (Fig 1C).

### 1.2.2 Behavior on the correct timescale

To understand brain dynamics underlying cognitive behaviors with fUS, it is crucial that the behavior matches the timescale of the hemodynamic signal. As one example, consider one important behavior of wide interest: evidence accumulation based decision making (Fig. 2A). Conventionally, systems neuroscientists train animals to accumulate evidence over relatively short timescales (100 ms-1.5 s). Both non-human primates and rodents can be trained to perform this task with high accuracy (Brunton, Botvinick, and Brody, 2013; Shadlen and Newsome, 2001). The hemodynamic signal measured by fUS, however, evolves over much longer timescales than either electrophysiology or calcium signals (Nunez-Elizalde et al., 2021) (Fig. 2B). Thus for fUS, as with fMRI, it is critical to be mindful of the behavior design. We therefore developed a modified version of the evidence accumulation task wherein the animal is trained to accumulate evidence over a more extended 3-5 seconds (Fig. 2A).

**Figure 2:**
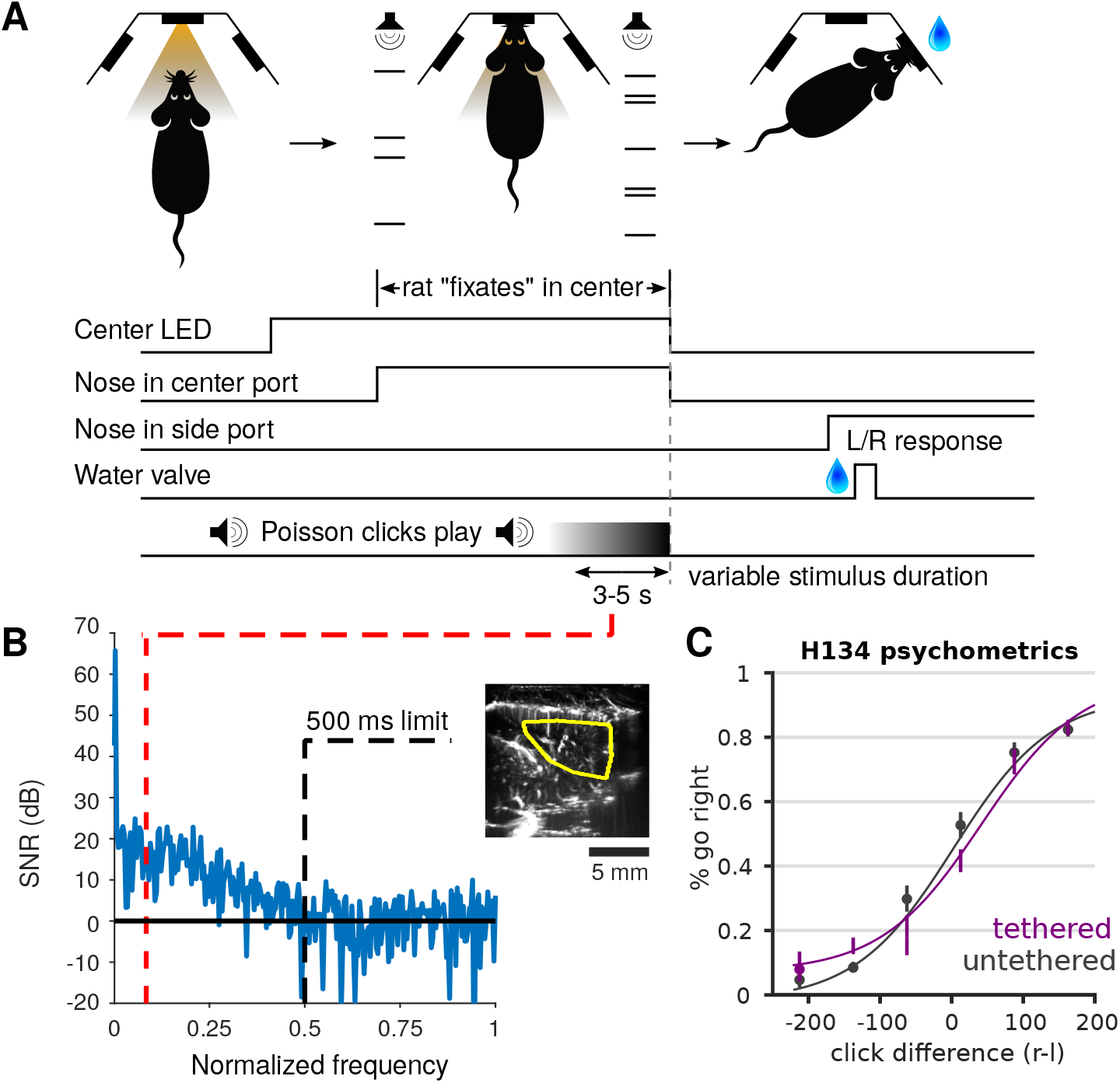
Decision making behavior on a time-scale appropriate for fUS recording. A: Example behavior of the previously used Poisson clicks task for rats (Brunton, Botvinick, and Brody, 2013). At each trial an LED lights to indicate the trial start. The rat then uses the center nose poke to initiate the trial, wherein an auditory Poisson click train is played on each of the left and right sides, simultaneously. In the version of the task used here, we extended the duration of the click trains, to play for 3-5 s until the end of click trains and the turning off of the center poke LED. The aforementioned signals to the rat that it should poke the left or right nose poke to indicate a selection, and is rewarded if it selected the side on which the greater total number of clicks were played. B: Frequency response of the total fUS signal averaged over an example area from a fUS imaged behavioral session. Previously used Poisson click trials tend to last approximately 500 ms (Brunton, Botvinick, and Brody, 2013), indicating that the rise/fall time of detectable physiological signatures is constrained to frequencies above 0.5 Hz (where 500 ms is 1/4 wavelength). While this sufficient for electrophysiology and calcium imaging time-scales, hemodynamic signals are much slower. The 3-5 s window in this experiment ensures that much slower physiological responses can be isolated (i.e., the red line of approximately 0.08 Hz). C: Psychometric curve of a tethered rat performing the task is comparable to a untethered rat

Specifically we modify an auditory evidence accumulation task previously developed in the context of electrophysiological and calcium imaging recoring; the Poisson Clicks task Scott et al., 2017; Hanks et al., 2015. In the Poisson clicks task, trials can be of variable duration, during which a stimulus is played on the right and left side of the animal. At the end of the trial, the animal gets rewarded with water if it orients towards the side that the largest number of auditory clicks originated from. We extend the trial duration from the original 100 ms-1 s duration to a 3-5s trial duration. We find that rats are able to perform the task with great accuracy and show behavioral characteristics similar to those observed in shorter timescale trials. The rats are also able to perform the task similarly whether they are tethered or not (Fig. 2C).

We validate that the trial duration is sufficient to capture the information content of the fUS signal by observing that the additional trial time lowers the minimum frequency content that can evolve during a single trial. The added observable low-frequency content includes much of the signal energy, e.g., when averaged over a large portion of the mid-brain (Fig. 2B), as well as averaged over specific brain areas (Sup. Fig. 1).

An important methodological advent in our study is the development and use of a tethering system for the fUS probe. Previous reports involving fUS in freely moving rodents required the experimenter to hold the wire by hand. Manual manipulation of the wire is both a distraction to the experimentor and prohibits long-term recording over many self initiated trials. The tether allows the animal to move freely and perform the behavior at the same accuracy and efficiency as without (Fig. 2C).

### 1.2.3 Functional ultrasound data curation

Our developed surgical procedures, behavioral training, and tethering system enable us to record long segments of hemodynamic activity that then must be processed to identify active brain areas. We thus develop a code-base including the main steps: 1) denoising 2) motion correction, and 3) decomposition of fUS data into core components.

For the first of these steps, the long timescales allow for efficient wavelet denoising (Donoho, 1995). Our code runs denoising the temporal dimension of the data, rather than image-based denoising of each frame for three main reasons. First, we aim to minimize mixing of signals. Second, time-domain denoising has an efficient computational complexity of *MNT* log(*T*), where *MN* is the number of pixels in the image, and can be parallelized over pixels. Finally, the time-domain signal statistics of hemodynamics should more closely match the polynomial assumptions of wavelet denoising (Daubechies, 1992).

The second processing task we include is motion artifact evaluation and correction. Paramount to scientific analysis of data is validating the integrity of the data. As imaging methods advance into new operating regimes, so too do the potential signal acquisition errors. Specifically, freely moving behavior can exacerbate motion-based errors. It is thus necessary to 1) isolate such errors early in the pipeline and 2) either correct such errors or propagate the uncertainty through to the ensuing scientific analyses. For fUS data, the motion-based artifacts come in two main forms: burst errors (Brunner et al., 2021) and lateral motion offsets.

We detect burst errors by computing the frame-wise norms (sum-of-squares of the pixels in each frame) and identifying frames that have abnormally large intensities (Fig. 3A). While multiple outlier detection methods could be applied, we find that setting a single threshold on the norm intensities is sufficient. In particular we note that the distribution of frames has a fairly sharp falloff, and we thus the dropoff in the histogram of frame norms to determine the appropriate decision threshold (Fig 3B, see Methods Section 2.6). Burst errors can also be corrected, as they tend to occur as single, isolated frames. We use polynomial interpolation (Brunner et al., 2021) between frames to fill in the missing frames to produce smooth video segments (Fig. 3A).

**Figure 3:**
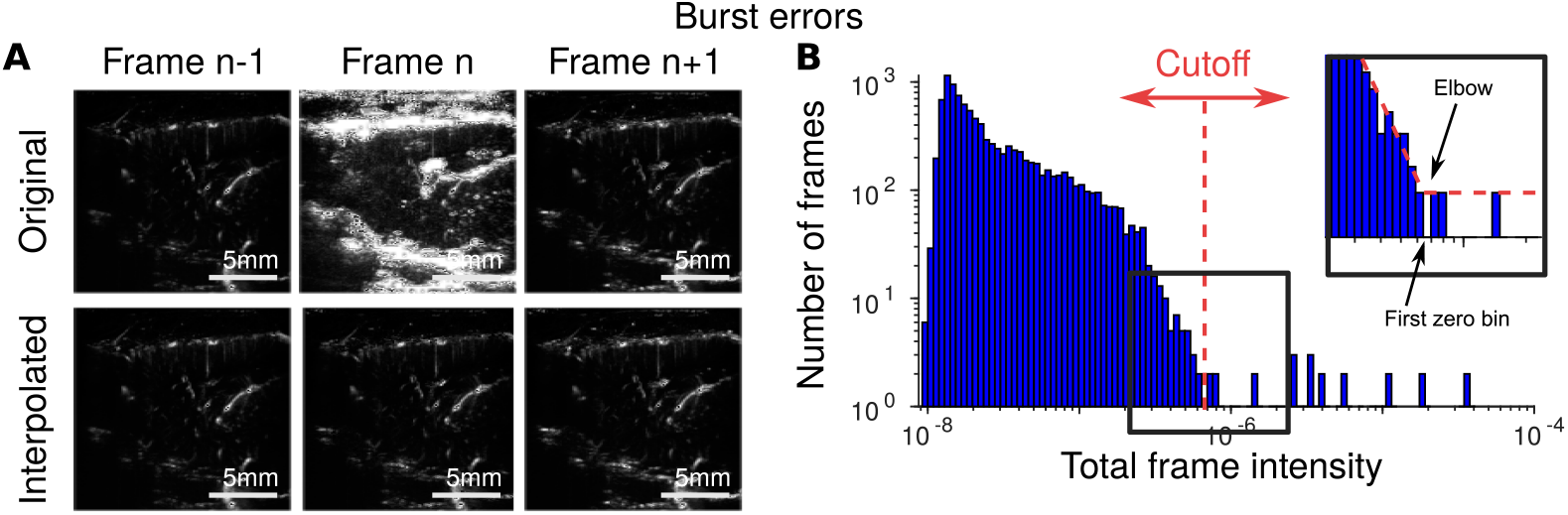
Identifying and correcting burst errors. A: Burst errors (top, middle frame), or sudden increases in frame-wide image intensity (A, Top, middle), are caused by sudden motion of the animal. Burst errors can be corrected using spline-based interpolation (A, bottom). B: Histograms of frame intensity can be used to easily identify burst frames from using a data-adaptive cutoff (see Methods Section 2.6).

Offset motion correction is more subtle than burst-error correction. As we did not observe rotational motion, only rigid motion of the probe with respect to the brain, we used the rigidmotion model for motion correction (Heeger and Jepson, 1992; Pnevmatikakis and Giovannucci, 2017). Under the rigid motion assumption, each frame is offset by a (potentially fractional) number of pixels in each direction. Rigid motion can most easily be assessed via the cross-correlation function of each frame with a reference image. The peak of the cross-correlation determines the shift that induces the maximal correlation with the reference image, however for noisy data with discrete offsets, identifying the sub-pixel peak of the autocorrelation can be challenging. We enable subpixel motion estimation by fitting the offset and scale of a Laplace function to the autocorrelation computed with integer shifts, and we use the inferred non-integer offset as the motion estimate (see Methods Section 2.7, Fig. 4A).

**Figure 4:**
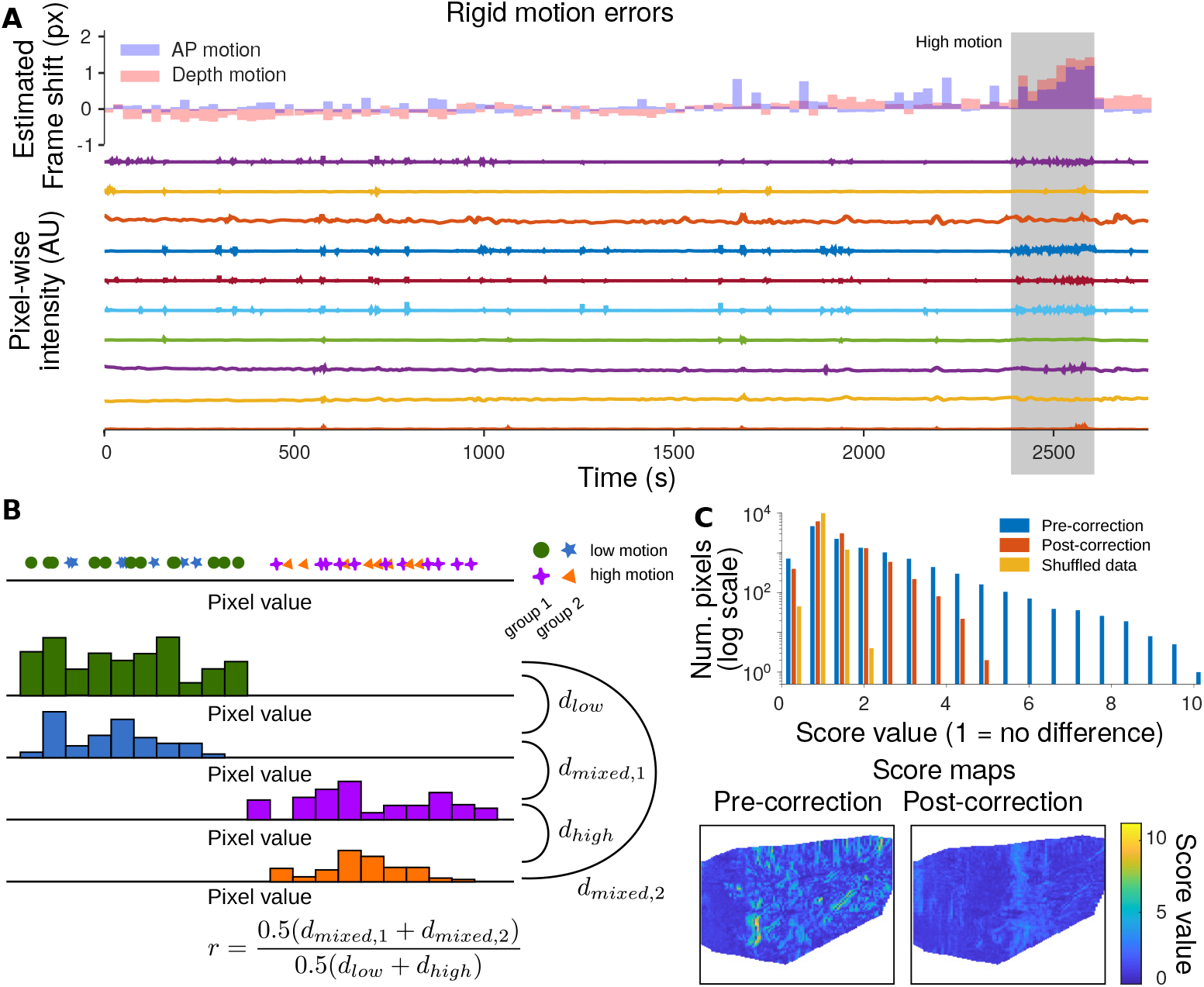
Identifying and reducing rigid motion. A: Rigid motion estimation corrects frameby-frame image offsets via frame-wise autocorrelations (top panel). Frames with high estimated motion correlate with changes in the blood-flow signal, as we quantify with a new metric: the MEPS score. B: MEPS Quantifies the effect of rigid motion error on pixel statistics enabling video stability assessment and evaluating of motion correction. The MEPS computes for each pixel the distribution of intensities for frames tagged as low-motion and frames tagged as high-motion (see methods). Each motion category is then further subdivided into multiple groups, to enable withinversus acrossgroup comparisons. The MEPS score quantifies the ratio of within-group versus across-group differences in these distributions. A score of ‘1’ means that the high-motion frames are indistinguishable from the low-motion frames in aggregate. C: (top) The MEPS value for a session with relatively high motion. The distribution of scores across pixels, both before motion correction (blue) and after motion correction (red). As a comparison, a shuffle test that randomly distributes data to the highand lowmotion tags is shown (yellow). The MEPS score is greatly reduced post rigid motion correction. (bottom) The spatial distribution of the MEPS score preand postmotion correction shows how clear anatomical features that create motion artifacts are greatly reduced.

Motion correction is often difficult to assess quantitatively. While movies may appear well aligned qualitatively, there may be small residual motion that can affect down-stream analysis. To assess motion’s impact on analysis, we assume that during repeated trials in a single session that the statistics of single pixels are similar in distribution. In essence, the values at pixel *i* over time should not have vastly different histograms when conditioned on different sets of frames. We quantify this effect by measuring the change in pixel value histograms for each pixel between high and low motion frames (see Methods Section 2.7). To account for natural variability across the entire session, the high-motion frames and low-motion frames are in turn split into randomly selected subsets. Blocks of frames are selected together to retain the temporal correlation of the signal. The histograms are then compared using an earth-mover’s distance (Haker et al., 2004), with the inter-motion distances (high vs. low) given as a fraction of the average intra-motion distance (low vs. low and high vs. high). For this metric, values closer to one indicate more similar histograms and therefore a lesser effect of motion on data statistics. We denote this metric the Motion Effected Pixel Statistics (MEPS) score.

Our metric enables us to both test the stability of recordings, as well as the effect of postprocessing motion correction. For three of the four recordings spanning two months we found that that the distribution MEPS scores over all pixels was close to a shuffle-test comparison indicating that the recordings are stable (Sup.Fig. 3). Moreover comparing preand postrigid motion correction for these sessions resulted in comparable histograms, indicating that motion correction does not introduce spurious motion artifacts. For the remaining session with high MEPS values, motion correction significantly reduced the metric, indicating much more stable videos post motion correction (Fig. 4C).

MEPS further allowed us to test motion correction more broadly on two additional recording sets and two additional rats (Sup.Fig. 4,5,6,7). To obtain a more precise metric we computed the standard deviations of the MEPS value histograms before (*σ*_*pre*_) and after (*σ*_*pre*_) motion correction, and compared those values to the standard deviation of the shuffle test MEPS score (*σ*_*shuffle*_). We found that on average, in 17 sessions across three rats, that the relative pre-motion correction standard deviation *σ*_*pre*_*/σ*_*shuffle*_ was 565% higher than after motion correction *σ*_*post*_*/σ*_*shuffle*_ (N=13).

### 1.2.4 Decoding of behavioral variables

Using fUS to characterize global brain activity requires a data processing pipeline that can relate the raw data to behavioral variables. Denoising and motion correction methods serve as the first segments of the pipeline. The final stages require identifying correspondences between the video data to task-related behavioral quantities. Such correspondences can be achieved either by computing on the native pixel-space videos, or, as often implemented for fMRI and more recently widefield data, over summary representations via decompositions of the video into individual components (sometimes referred to as regions-of-interest; ROIs).

With the registered data, we first provide pixel-domain tools to map pixels associated with different aspects of the evidence accumulation task. The first is a comparison between conditions (e.g., activity within and outside of a trial; Fig. 5A-C) and the second is a correlation map that identifies pixels tuned to specific events (e.g., reward times Fig. 5D,E).

**Figure 5:**
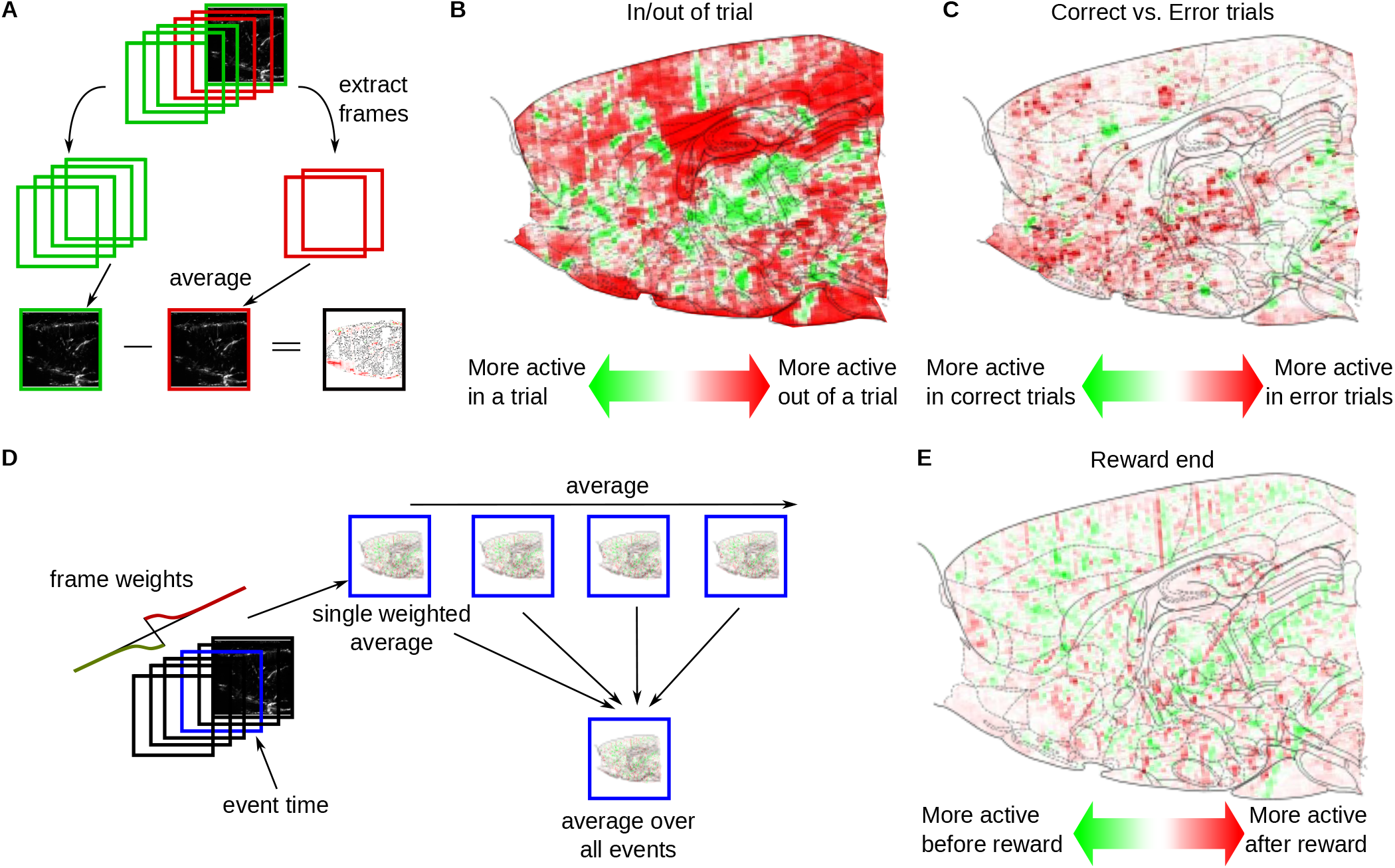
Event-triggered maps of global brain hemodynamics. A: To determine differences between global hemodynamic activity during different epochs, frames during those epochs (e.g., during a trial and between trials) were averaged separately. The difference between the two maps was then taken to produce the difference map comparing two conditions. B: The difference between the average data frame during a trial and between trials demonstrates areas with different hemodynamic activity (computed as in panel A). C: Similarly, the difference between the average data frame within a correct trial versus an incorrect trial demonstrates even more differences, especially in the cortical areas M2 and RSD. D: To determine event-triggered changes in activity, fUS activity was correlated with a synthetic trace representing a derivative of activity time-locked to the event. Here the event was the end of correct trials. A Gaussian shape with a width of 2 seconds (4 frames) was selected. The resulting estimate of the time derivative is shown in the top panel, with red indicating locations with increases in brain activity time-locked to end of correct trials, and green indicating decreases. E: Example event-triggered changes locked to the end (last frame) of correct trials, as computed via event-triggered correlation maps in panel D.

We implemented comparisons between conditions as a difference-of-averages (Fig. 5A); The average of all frames in condition two is subtracted from the average of all frames in condition one. The difference removes baseline activity levels constant over both conditions. For example, maps comparing activity within and outside of a trial demonstrate significantly decreased activity in V2 visual cortex (V2MM) (Fig. 5B). Similarly, error trials seem to induce more activity in area Motor Cortex (M2) and Retrosplenial Cortex (RSD) than correct trials (Fig. 5C).

Correlation maps depend on instantaneous events, rather than spans of time. We create correlation maps that average the frames surrounding each event to discover which pixels are most active in the vicinity of those events. To prioritize frames closest to the event, we weigh the frames in the average with an exponential-squared fall-off from the nearest event (Fig. 5D). Furthermore, to differentiate pixels active before and after the event, we weigh frames right before an event with negative values and frames right after with positive values (Fig. 5E). The correlation maps can further be used to compute the responses across brain areas by integrating the computed values over voxels corresponding to registered brain areas (see Methods Section 2.10, Supp.Fig. 10,11,12,13). While informative, these maps ignore the high degree of temporal correlation between pixels, and can be sensitive to single-pixel fluctuations. Thus we expand the data processing pipeline with a signal extraction stage to permit efficient and accurate decoding (Fig. 6A). We leverage recent advances in time-trace extraction from neural imaging data which seeks to identify the independent temporal components underlying the full fUS video (Fig. 6B). This method, Graph-Filtered Time-trace (GraFT) analysis, iterates between updating the time-traces that best represent each pixels timetrace through a linear combination provided by the spatial weights, and then updating the spatial weights based on the new, updated time-traces (Charles et al., 2022). Importantly, GraFT uses a data-driven graph to abstract away spatial relationships within the fUS video to learn large-scale and non-localized activity profiles. The result is a decomposition *Y ≈Z*^*T*^ Φ where the pixel-by-time matrix *Y* is described by the product of a component-by-time matrix Φ that has each row represent one independent time-trace in the data with the pixels-by-component matrix *Z*^*T*^ of spatial weights describing which pixels are described by which independent time-traces in Φ. We selected GraFT due to its prior success in analyzing widefield calcium imaging data, which similarly benefits from a lack of *a-priori* assumptions on anatomical structures Charles et al., 2022.

**Figure 6:**
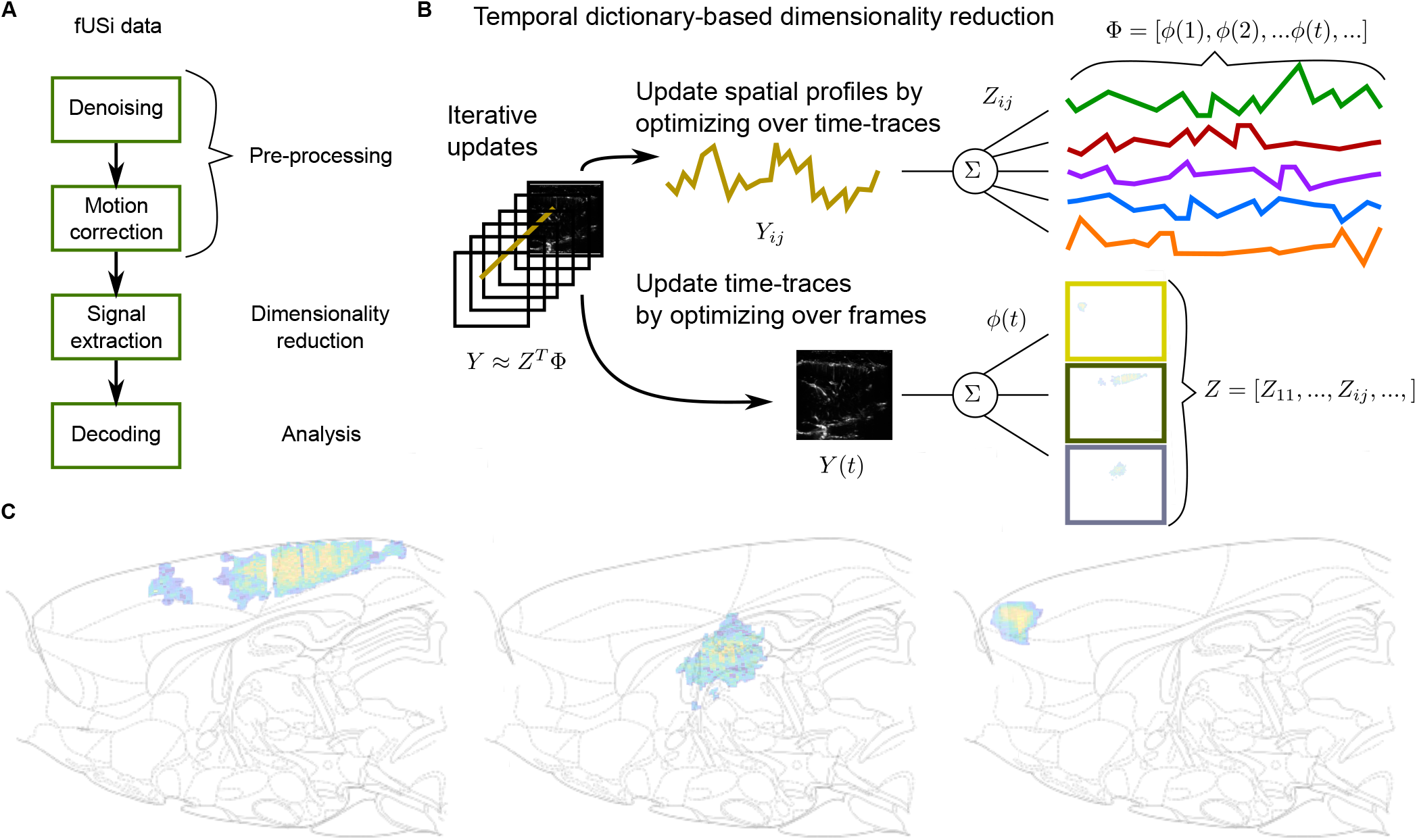
Decomposing fUS data using GraFT. A: The general pipeline for fUS decoding is to first denoise and motion correct the data via the previously mentioned pre-processing steps. Signal extraction then identifies the basic temporal signals from the raw data. Finally decoding, in our case using support vector machines, is applied. B: Signal extraction is accomplished via GraFT (Charles et al., 2022) GraFT is a data-driven decomposition that seeks to describe each pixel’s time-trace in the movie as a sparse linear combination of a dictionary of basic temporal components in the data. GraFT uncovers these basic components via an unsupervised dictionary learning procedure that iteratively updates the time-traces and the spatial profiles (i.e., how much each time-trace is present in each pixel). Key to GraFT is that the pixel decompositions are correlated via a learned graph based on temporal correlations (Charles et al., 2022). C: GraFT uncovers spatial areas exhibiting independent activity. Three example areas (of the 45 learned areas) are shown, and the full complement provide a low-dimensional description of the data.

The result of GraFT is a reduced description of the fUS video that identifies independently co-active regions (e.g., Fig. 6C, Sup.Fig. 15,16, Supp.Fig. 19). For example we identify RSD being highly co-active within itself and with parts of M2. Importantly, GraFT does not assume spatial cohesion, indicating that the connectedness of the identified co-active areas is truly a product of the data statistics and not an assumption built into the algorithm. We observe this spatial cohesion in 6 recording sessions across two rats (Supp.Fig. 15,16, 20, 21, 22, 23)

**Figure 7:**
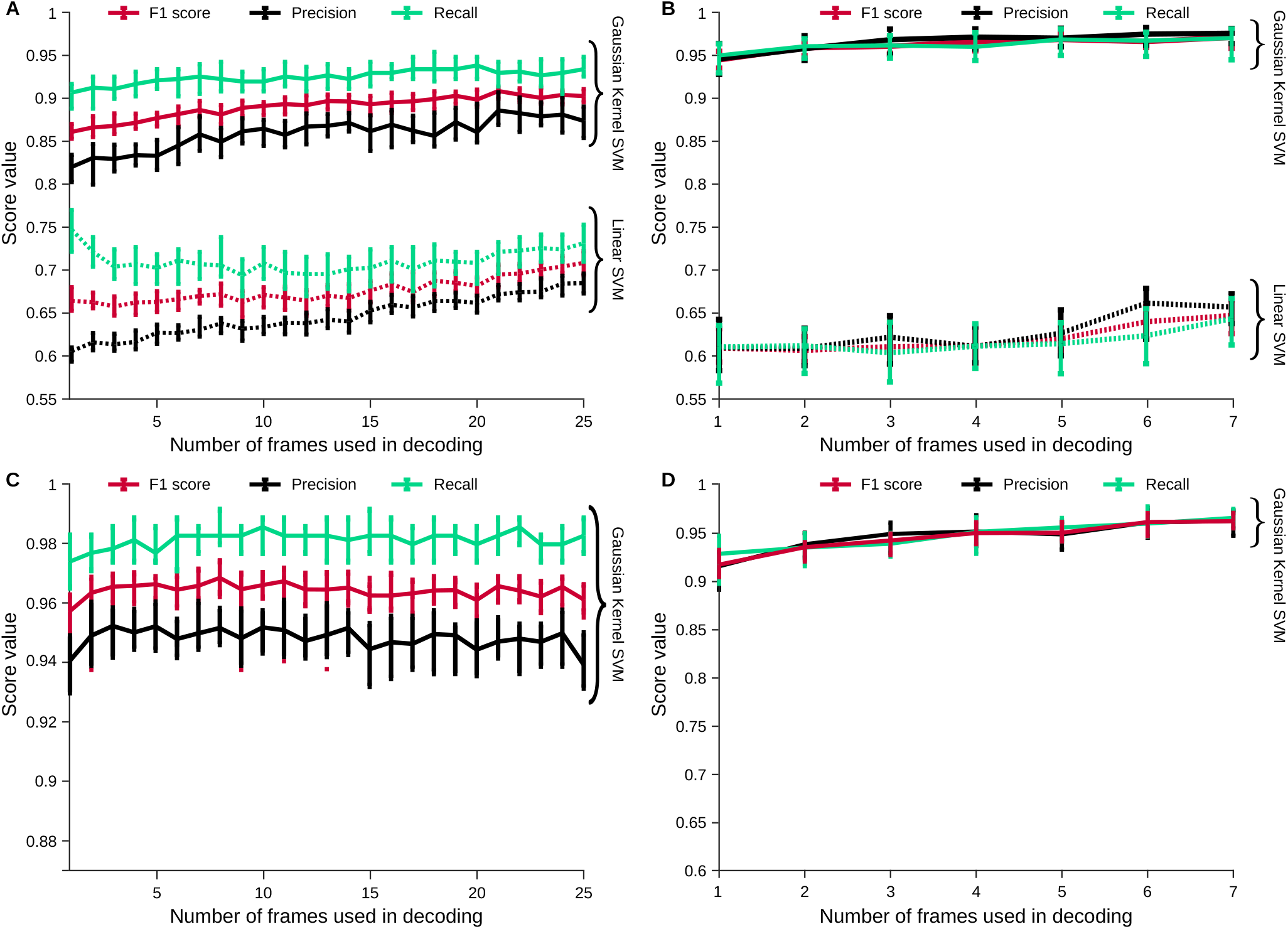
Decoding cognitive variables from decomposed hemodynamic activity. A: Decoding inand outof trial imaging frames can be computed in the learned low-dimensional representation from Figure. 6. We find that classifying frames is much more accurate with a Gaussian Kernel SVM than with a Linear SVM, indicating complex nonlinear activity in fUS brain-wide dynamics. Moreover, the number of consecutive frames used in decoding increases accuracy slightly, indicating non-trivial temporal dynamics. B: Similarly, when decoding hard vs. easy trials (as determined by the difficulty parameter *γ*), we similarly see a large boost in performance using a nonlinear decoder. In fact the linear SVM performs worse in decoding hard vs. easy trials than it does in determining in vs. out of trial frames, while the nonlinear decoder performs better. C: Same Gaussian kernel SVM decoding as (A) for a second data set two months later shows comparable decoding of inand outof trial activity. D: Same Gaussian kernel SVM decoding as (B) for a second data set two months later shows comparable decoding of task difficulty.

Another benefit of the GraFT decomposition is that the reduced dimension enables us to efficiently decode behavioral variables by using the 30-60 time-traces instead of the *>*20K pixels as regressors.

As an initial test of regression over the GraFT components, we decoded whether a rat was within or out of a trial. We found that the GraFT decomposition, despite being a linear decomposition of the data, did not yield high decoding levels with a linear support-vector machine (SVM) classifier (Fig. 7A). When we applied a nonlinear SVM—using a Gaussian radial basis function (RBF) kernel (see methods)—to the same data, we instead obtained very high decoding performance (Fig. 7A), indicating that cognitive variables may be represeted in a nonlinear fashion in fUS data. We obtained similar results on more intricate cognitive variables, such as decoding whether the trial was difficult or easy, from only the fUS brain images (Fig. 7B). In general, we found similar decoding results across a number of behavioral cognitive variables (Supp. Fig. 24).

### 1.2.5 Stability over multiple sessions

A major aim of our study is to be able to record from the same rat performing a cognitive task over many months. We therefore asses the stability of our recordings over short and long timescale by comparing fUS data collected in the same rat during sessions spaced days and months apart. We first found that all data could be registered via rigid motion. The estimated movement of the probe between sessions was ≤ ∓0.5 mm in AP, and ≤ ∓ mm in depth (Fig. 8A). The registration was further validated by comparing the average image (i.e., an ‘anatomical image’) between sessions. The correlation between the anatomical image between sessions was high both between days and across months (Fig. 8B). These shifts allow us to further compare behaviorally dependent measures, such as the changes in event-triggered brain activity, over many months (Fig. 8B, Supp. Fig. 26, 27,10,11,12,13).

**Figure 8:**
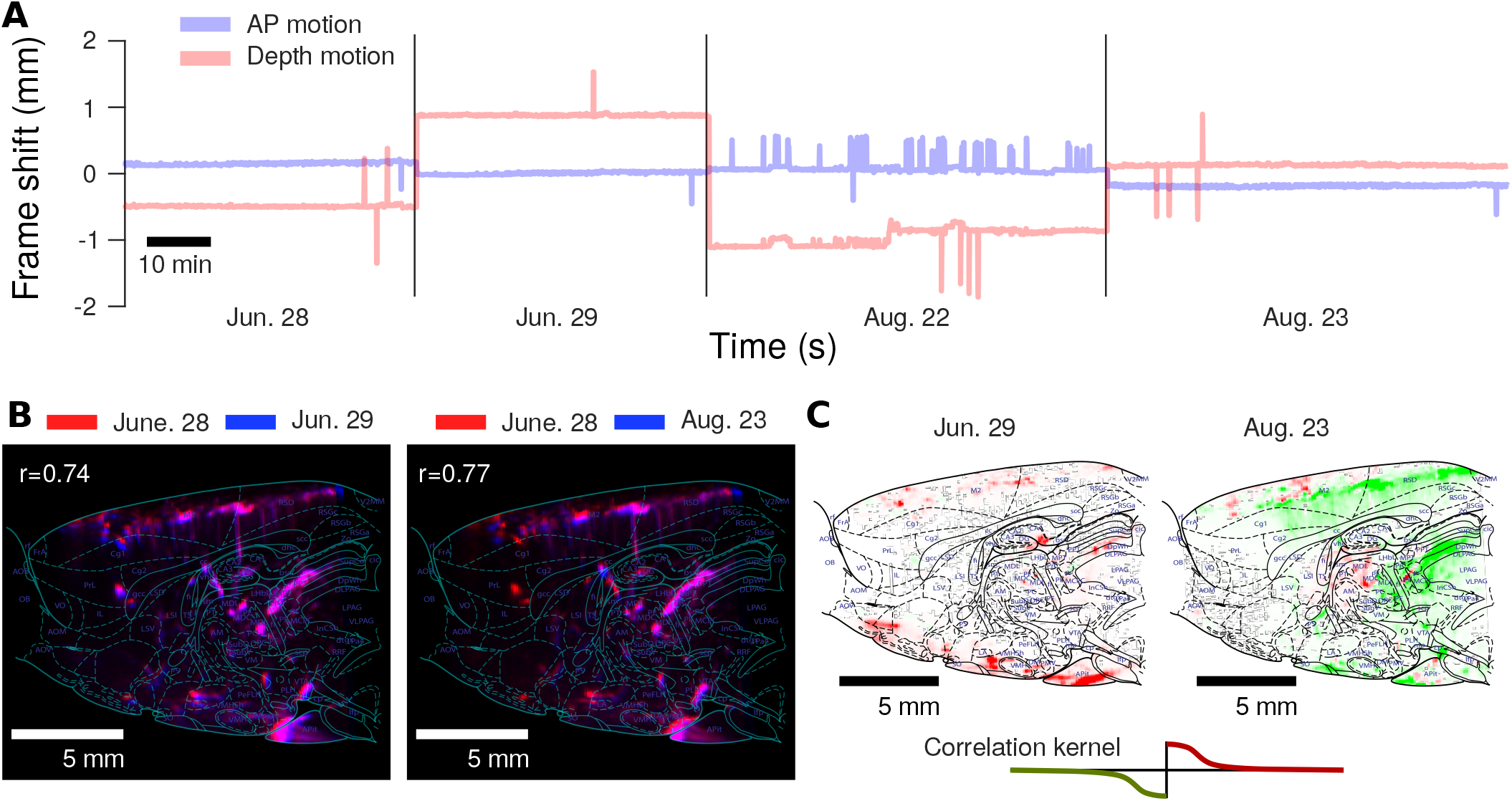
Stable fUS recording over two months. A: Rigid motion estimation infers at most only a ±2 mm shift in either depth or the anterior-posterior direction over two months of chronic implantation and recording. Interestingly, AP motion is much more stable, with typical offsets between recording days at *<* 0.5 mm. B: Image stability can be assessed over multiple months by comparing the anatomical images computed via averaging over time. We note that the anatomy is stable over long time-frames, with very high correlations both between the anatomical images on subsequent days, as well as two months later. C: As the anatomical plane is stable over long time-frames, changes in hemodynamic activity during the decision making task can be detected over the course of chronic recording. The maps of differential brain activation at the start of a trial can be compared between sessions months apart to study

## 1.3 Discussion

In this paper, we have presented detailed experimental and algorithmic techniques that enable practical chronic implementation of functional ultrasound (fUS) imaging of the brain in freely moving rodents. The criteria that we have set for the success is to obtain chronic stable fUS data while the freely moving animals perform cognitive behaviors. Our solution was realized by: 1) devising a surgical protocol that allowed us to open a large craniotomy and stabilize the functional ultrasound probe over many months, 2) developing and validating a rodent behavior that matches the timescale of the hemodynamics, 3) developing motion artifact detection and correction algorithms that further stabilize the fUS images, and 4) developing analytical methods to extract brain activity aligned with behavioral events.

Prior work demonstrating fUS in freely moving rodents (Urban et al., 2015; Sieu et al., 2015; Bergel et al., 2018; Bergel et al., 2020; Rabut et al., 2020) was restricted to sensory stimulation or spontaneous movement and did not extend to chronic recordings during cognitive behaviors. Moreover, previous studies required that the experimenter actively hold the tether, which limits the experimenter’s ability to monitor and respond to the experiment’s progress. In our study, we have developed a tethering system that allows animals to move freely without intervention over extended periods of time. This has allowed us to extend the applicability of fUS to longer sessions, which may be needed for self-initiated, or complex cognitive behaviors, e.g., parametric working memory and spatial navigation tasks (Fassihi et al., 2014; Aronov and Tank, 2014). As a prime example, there is increased interest in studying freely moving naturalistic behaviors, e.g., foraging, wherein movement is a crucial component of its execution. Optimizing the imaging apparatus to enable the most natural animal movement will be increasingly crucial to the animals behave most naturally.

One primary concern with data from freely moving animals is the potential for severe motion artifacts. Our study took a two-pronged approach to solving this problem. First we introduced new surgical techniques to adhere and stabilize the probe. We further provided motion correction tools that remove burst errors and align frames via rigid motion correction. For three of the four recordings spanning two months we found minimal motion impact even without any software motion correction. For the remaining recording, motion correction significantly reduced the metric values, indicating a reduction in potential motion artifacts.

Motion correction is often assessed via correlations of the images to a template image. Such correlations often rely sharp features for accurate alignment. In fUS, there are fewer internal features as opposed to e.g., micron-scale calcium imaging. Our metric instead relies on temporal statistics to assess potential shifts in activity histograms on a per-pixel basis. In general we envision this metric, and other activity-oriented metrics, complementing spatially-based assessment.

In our pipeline, we implemented polynomial interpolation to fill in missing burst-error frames followed by rigid motion correction. Interpolation was included to enable any subsequent analysis, which requires regularly sampled data. Advanced analysis methods, however, could instead treat burst-error frames as “missing data”, removing the need to interpolate (Tlamelo et al., 2021; Pimentel-Alarcón et al., 2016). Similarly, while rigid motion correction significantly reducing the overall motion, future work should explore more flexible deformation based correction, e.g., diffeomorphic mappings (Ceritoglu et al., 2013) used in *ex vivo* brain alignment, or patch-based affine mappings (Hattori and Komiyama, 2021).

A well known effect of both motion and erroneous motion correction (especially with global, rigid motion) is “spurious correlations” (Fellner et al., 2016; Power et al., 2012) (Sup. Fig. 28). Spurious correlations can make completely unrelated pixels appear correlated: confounding many functional connectivity analyses, e.g., in fMRI (Power et al., 2012). These correlations (derived in Methods Section 2.13) depend on the difference-of-means as compared to the per-pixel sample variance. The total number of frames that are affected by motion play a critical factor and demonstrate the importance of stable imaging.

Furthermore, the use of component-based analyses have been shown to reduce the effects of spurious correlations (Perlbarg et al., 2007). Our use of GraFT here assists in two ways. First, the graph constructed in GraFT is a k-connected graph. Thus weaker correlations are removed. Second, the components learned must have the same time-trace. Thus signals contaminated with motion related high/low alternating patterns would at worst be learned as a separate component, if at all.

The benefits of GraFT against spurious motion has a number of limitations. For one, if the difference of means between mixed pixels is large compared to the variance, a step-function like sequence might be learned. A second danger is if a large component with very well defined boundaries undergoes motion. In this case, pixels at the edges on either side of the component would have a shared time course that is on (or off) during shifted (or not shifted) frames. This artifact would be picked up by the dictionary learning in GraFT as a unique component with the on/off signature. None of the components we learned displayed either of these statistical patterns, indicating that these effects were well controlled by our stable imaging and data pipeline.

To ensure complete clarity and reproducibility of our procedures, we have provided detailed protocols that include step-by-step schematic description of the surgical procedures, open source code for motion detection, correction and data analysis, along with a detailed description of the codes and an up-to-date repository for updated analysis methods. It complements already existing detailed protocols in the literature for fUS imaging in head fixed mice (Brunner et al., 2021).

While the potential for imaging plane offsets (D-V) exists, we do not notice significant deviation as based on longitudinal consistency computed via session-wise average-image correlations Fig. 4C, Supp. Fig. 8B). Additional assessment of D-V stability can be obtained by emerging volumetric fUS recordings (Brunner et al., 2020). These recordings will enable full 3D alignment of the brain between sessions.

Our analysis tools represent a complete pipeline for fUS, up to and including decoding analyses. For brain parcellation, rather than relying on averaging brain areas from an anatomical atlas, we instead chose to use an activity-based decomposition, GraFT. GraFT, despite not making anatomical assumptions, was able to find both activity isolated within individual brain areas and activity crossing anatomical boundaries. For decoding we used support-vector machines. We found that a linear SVM decoder could not accurately decode behavioral variables from brain data, while a nonlinear kernel-SVM decoder achieved high decoding accuracy. Moreover, using multiple consecutive frames improved decoding performance. These results raise important questions as to how brain dynamics encode task variables, e.g., implying nonlinear, time-dependent encoding in global brain dynamics. Further work should continue analyzing the encoding principles underlying brain activity on a global scale during cognitive behaviors.

fUS is thus a complementary imaging technique to traditional targeted imaging: fUS can give us unprecedented insights into global brain dynamics (mesoscale mechanisms) while other methods, e.g., electrophysiology, can give us detailed insights into the inner workings of one or more brain areas (microcscopic mechanisms).

Systems neuroscience is fast approaching the point where local microscopic activity and global brain dynamics must be understood together in complex tasks. fUS offers a vital complementary technique to more targeted imaging modalities, e.g., such as electrophysiology and calcium imaging, that can be implemented in freely moving animals. In particular, fUS give us the opportunity to record and screen many brain areas simultaneously, enabling experimentors to find new brain areas involved in a given behavior. Identifying new task-related regions is especially critical in complex behaviors wherein we lack a full understanding of the brain-wide networks involved. The capability to scan the brain and identify new regions will only improve as fUS technology improves. Volumetric fUS (Brunner et al., 2020; Rabut et al., 2019), for example will enable near full-brain imaging, and new signal processing for ultrasound (Bar-Zion et al., 2021) have the potential to vastly improve the speed and signal quality of fUS data reconstruction from the ultrasound returns.

The combination of global activity imaged with fUS and local recordings targeted to new brain areas discovered via global analyses offers a critical path forward. Specifically, fUS can give us unprecedented insights into global brain dynamics (mesoscale mechanisms) while other methods, e.g., electrophysiology, can give us detailed insights into the inner workings of one or more brain areas (microcscopic mechanisms). fUS can also be combined with optogenetics (Sans-Dublanc et al., 2021) so one can study the global effects of local perturbations (Lee et al., 2010). Given the aforementioned, fUS in freely moving animals will provide a critical new technology in refining our understanding of concerted brain dynamics during naturalistic behavior.

## Supporting information

Supplemental surgery guide and code documentation

## Acknowledgements

The authors would like to thank Alan Urban, Gabriel Montaldo, Emilie Macé for advice concerning functional ultrasound imaging, Diogo Guerra for the surgical procedures illustrations, Emily Jane Dennis for help with the registration and the rat brain atlas, Princeton University Physics Department Workshop, Charles Kopec, David Tank, Brody lab animal training technicians. AEH acknowledges support by NIH grant 1R21MH121889-01.

## Author Contributions

AEH and DT: designed and performed the surgeries, designed and built freely moving behavior arena, and performed animal training and data collection. TBM performed the animal behavior analysis. YZ performed the registration of the fUSi data to the rat brain atlas. RS and ASC coded the data processing pipeline, including motion correction, denoising, and demixing. AEH, DT, RS and ASC all performed data analysis. RS and ASC performed all correlation and decoding analyses. ASC and CDB supervised the project. All authors contributed to the writing of the paper.

## Data availability

Data used in this study is available at https://osf.io/8ecrf/.

## Code availability

The data processing pipeline is coded in MATLAB and available at https://github.com/adamshch/fUSstream.

## 2 Methods

### 2.1 Rat Behavioural Training

#### 2.1.1 Training overview

Animal use procedures were approved by the Princeton University Institutional Animal Care and Use Committee and carried out in accordance with National Institute of Health standards. All animal subjects were male Long-Evans rats. Rats were pair-housed and kept on a reverse 12-hour light dark cycle; training occurred in the rats’ dark cycle. Rats were placed on a water restriction schedule to motivate them to work for water reward.

fUS can be performed on behaviors with timescales that matches the timescale of the hemodynamic changes measured which is on the scale of 3 s Nunez-Elizalde et al., 2021. Therefore, it is important to train animals on behaviors matching this long timescale or stretch the temporal dimension by elongating the trials of already established behaviors. There are many behaviors that can be studied for example parametric working memory which already employs long delay periods between two stimuli thus suitable fUS. For this paper, we have extended the evidence accumulation task in (Brunton, Botvinick, and Brody, 2013) to longer trials up to 5 s. In the following, we will delineate the training pipeline that we have established to train rats, in a high throughput manner, on this task that we named “Ultralong evidence accumulation”.

The training proceeds as follows: Approximately 75% of naive rats progress successfully through this automated training pipeline and perform the full task after 4-6 months of daily training. Rats were trained in daily sessions of approximately 90 minutes and perform 100-300 trials per session. We trained rats on extended version of the Poisson Clicks task (Brunton, Botvinick, and Brody, 2013). Rats went through several stages of an automated training protocol. The behavior is done in an operant behavioral chamber with three nose pokes. It is crucial to mention that the final configuration used in this behavior is the right or left port deliver the water rewards and the central nose poke has an LED that turns on and off depending on whether the animal is poking in it or not.

#### 2.1.2 Training Stage 1

In training stage 1, naive rats were first shaped on a classical conditioning paradigm, where they associate water delivery from the left or right nose port with sound played out of the left or right speakers, respectively. After rats reliably poked in the side port, they were switched to a basic instrumental paradigm; a localized sound predicted the location of the reward, and rats had to poke in the appropriate side port within a shortening window of time in order to receive the water. Trials had inter-trial intervals of 1-2 minutes. For the first couple of days, water was delivered only on one side per session (i.e., day 1: all Left; day 2: all Right; etc.). On subsequent days, Left and Right trials were interleaved randomly. This initial stage in the training lasted for a minimum of one week, but with some rats it can be as long as two weeks.

#### 2.1.3 Training Stage 2

In training Stage 2, we aimed at enforcing the center “fixation period” called “grow nose in center”. At first, the rat needed to break the center port Infrared (IR) beam for 200 ms before withdrawing and making a side response. Over time, this minimum required time grew slowly, so that the rat was required to hold its nose in the center port for longer periods of time before allowed to withdraw. The minimum required fixation time grew by 0.1 ms at every successfully complete center poke fixation trial. The clicks were played through the whole fixation period. The offset of the clicks coincided with the offset of the center light, both of which cued the rat that it was allowed to make its side response. Rats are then trained to center fixate for 3-5 s. Premature exit from the center port was indicated with a loud sound, and the rat was required to re-initiate the center fixation. Rats learned center fixation after 3-8 weeks, achieving violation (premature exit from center port) rates of no more than 25% (violation meant the rat’s nose breaking the fixation for a period longer than 500 ms, the violation time interval was varied among rats with the majority of rats had a violation period around 500 ms and some went down to 100 ms) . Unless indicated otherwise, these violation trials were ignored in all subsequent behavioral analyses.

#### 2.1.4 Training Stage 3

In training stage 3, we have Poisson clicks trains played from both right and left ports. Each “click” was a sum of pure tones (at 2, 4, 6, 8, and 16 kHz) convolved with a cosine envelope 3 ms in width. The sum of the expected click rates on left and right (rL and rR, respectively) was set at a constant *rL* + *rR* = 40 (for rats) clicks per second. The ratio 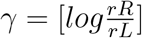 was varied to generate Poisson click trains on each trial, where greater |*γ*| corresponded to easier trials, and smaller |*γ*| corresponded to harder trials. In this stage we use a *γ* = 4 which means the easiest trials.

#### 2.1.5 Training Stage 4 to 6

In training stage 4, we add a second type of difficulties meaning we introduce trials more difficult than the trials from stage 3. During the stage, the stimulus being played is a mix of *γ* = 3 and *γ* = 4 interleaved trials.

In training stage 5, we add more difficult trials so we have then *γ* of 1-2-3-4 intermixed.

In training stage 6, we train the rats on the same task that as in stage 5 but beginning to shorten the trial from 5 s to 3 s slowly (with an increment of 100 ms at each trial). The reason to do is that we want the rats to be exposed to trials with variable duration. At the end of this stage the rat is performing the task at variable trials duration (between 3-5 s) and *γ* (1-2-3-4).

#### 2.1.6 Final Training Stage

In the final stage, each trial began with an LED turning on in the center nose port indicating to the rats to poke there to initiate a trial. Rats were required to keep their nose in the center port (nose fixation) until the light turned off as a “go” signal. During center fixation, auditory cues were played on both sides with a total rate of 40 Hz. The duration of the stimulus period ranged from 3-5 s. After the go signal, rats were rewarded for entering the side port corresponding to the side that have the highest number of clicks. A correct choice was rewarded with 24 *µ*l of water; while an incorrect choice resulted in a punishment noise (spectral noise of 1 kHz for a 0.7 s duration). The rats were put on a controlled water schedule where they receive at least 3% of their weight every day. Rats trained each day in a training session on average 90 minutes in duration. Training sessions were included for analysis if the overall accuracy rate exceeded 70%, the center-fixation violation rate was below 25%, and the rat performed more than 50 trials. In order to prevent the rats from developing biases towards particular side ports an anti-biasing algorithm detected biases and probabilistically generated trials with the correct answer on the non-favored side. The rats that satisfied those final training criteria were the ones that were used for the fUS experiments.

#### 2.1.7 Psychometric

The performance of each rat (pooled across multiple sessions) was fit with a four-parameter logistic function:

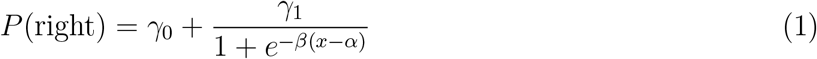

where *x* is the click difference on each trial (number of right clicks minus number of left clicks), and *P* (right) is the fraction of trials when the animal chose right. The parameter *α* is the *x* value (click difference) at the inflection point of the sigmoid, quantifying the animal’s bias; *β* quantifies the sensitivity of the animal’s choice to the stimulus; *γ*_0_ is the minimum *P* (right); and *γ*_0_+*γ*_1_ is the maximum *P* (right). The lapse rate is (1-*γ*_1_)/2. The number of trials completed excludes trials when the animal prematurely broke fixation.

### 2.2 Surgical Procedures

Animal use procedures were approved by the Princeton University Institutional Animal Care and Use Committee and carried out in accordance with National Institute of Health standards. All animal subjects for surgery were male Long-Evans rats that have successfully gone through the behavioral training pipeline.

#### 2.2.1 Craniotomy and headplate implantation

The surgical procedure is optimized in order to perform a craniotomy that goes up to 8 mm ML x 19 mm AP cranial window on the rat skull and be able to keep it for many months in order to be able to perform chronic recordings. A detailed appendix at the end of the manuscript with the step by step surgical procedure can be found along with illustrations that walk through every surgical step.

In brief, the animal is brought to the surgery room, then placed in the induction chamber with 4% isoflurane. Administer dexamethasone 1 mg/kg IM to reduce brain swelling and buprenorphine 0.01–0.05 mg/kg IP for pain management. Put the animal back to the induction chamber (with 4% isoflurane) until the animal achieves the appropriate depth of anesthesia. The rat hair,covering his head, is shaved using a trimmer. It is important to remove all the hair on the head skin that will be the site of incision afterwards. Put the animal head in the ear bar. Clean copiously the shaved head with alternating betadine and alcohol. From now on, the surgical procedure becomes a sterile procedure. Inject 0.15 ml of a mixture of lidocaine and norepinephrine (2% lidocaine with norepinephrine 20 *µ*g/mL mixture) under the head skin where the incision is planned. The skull overlying the brain region of interest is exposed by making an incision (∼10-20 mm, rostral/caudal orientation) along the top of the head, through skin and muscle using a surgical scalpel feather blade. Tissue will be reflected back until the skull is exposed and held in place with tissue clips. The surface skull is thoroughly cleaned and scrubbed using a double ended Volkman bone curette and sterile saline. The cleaning and scrubbing continue until the surface of the skull is totally clean and devoid of signs of bleeding, blood vessels or clotting. Bregma is marked with a sterile pen then a rectangle is drawn using a sterile waterproof ruler. The center of mass of the rectangle is Bregma and it is 19 mm AP x 8 mm ML. Using the surgical scalpel blades, diagonal lines are engraved on the surface of the skull. Metabond (C&B Metabond Quick Adhesive Cement System) is then put all over the surface of the skull (except the area onto which the rectangle is drawn) It is highly advisable to put low toxicity silicone adhesive KWIK-SIL in the region marked by the pen using gas sterilized KWIK-SIL mixing tips to avoid that any metabond covers this area. Using a custom designed positioning device, the sterile headplate is positioned over the drawn rectangle on the surface of the skull. The headplate is pushed down pressing on the skin and Metabond is put copiously t ensure the adhesion of the headplate to the skull surface. The headplate is then slowly unscrewed from the positioning system and the positioning system is then slowly raised leaving the headplate firmly adhered to the skull. The craniotomy can then be performed. Slowly use the piezosurgery drilling (using a micro-saw insert) to remove the skull demarcated by the rectangle drawn by the sterile pen beforehand . The piece of skull cut is then removed very slowly with continuous flushing with sterile saline. The sterile headcover is then screwed to the headplate then a layer of KWIK-SIL is added on the sides of the headplate. The rat is left to wake up in the cage with a heat pad under the cage. Afterwards, the animal takes 10-14 days to recover during which they are observed for any signs of pain or inflammation.

#### 2.2.2 Chronic probe holder insertion

After the recovery period from the surgery, the following steps are performed to insert the mock probe and streamline the fUS imaging so that we do not need to anaesthetize the rat every time we are performing an imaging session. After the recovery from the surgery, the rat is ready for experiments. In order to perform the fUS imaging, there should be a direct access to the open cranial window (that we opened via craniotomy during the surgery). First the rat is anaesthetized. Throughout the whole coming procedure, sterile procedures are to be strictly followed in order to prevent infection of the exposed brain surface. The rat is then put in ear bars centered. The headcover is unscrewed and removed. The brain surface is flushed with sterile saline, then a piece of Polymethelyene Pentene film (PMP), cut to a size that fits the craniotomy, is glued onto the surface of the brain by carefully applying small drops of VETBOND around the sides. It is crucial to make sure that the PMP is tightly glued to the surface of the brain. This step is very crucial to ensure the total minimization of the brain motion during fUS in a freely moving setting. Afterwards the probeholder is screwed onto the headplate. Note that in this case the headcover is not screwed back. Ideally when you look through the probe holder, one can see the brain surface onto which the PMP is glued. We add a mock probe (a 3d printed analogue of the fUS probe) for further protection while the rat is in his cage.

We are able to achieve many months sterility and persistence of the craniotomy due to our multi-layered approach to preserve its integrity: first by employing fully sterile surgical procedures, by sealing it with a sterile transparent polymer and by having a mock probe that gives an extra layer of protection from debris and contaminants. The ability to obtain a reproducible imaging plane across session over many months was obtained due to accurate alignment of the headplate to the skull using our custom designed positioning system, the alignment of the probe holder to the headplate on which it is tightly screwed and the size of the inner surface of the probe holder that prevents vibrations of the fUS probe. In some cases, the inner surface of the probe can be made tighter by putting a clip around it after the fUS probe is inserted inside the probe holder.

### 2.3 Functional Ultrasound Imaging procedure

When the rat is not doing the task or being trained in its behavioral box, there is a “mock” 3D printed probe that is basically inserted in the probe holder in order for the animal to have the same weight over their head all the time (as explained in the previous section “Chronic probe holder insertion”). The aforementioned configuration gives us the advantage of not having to anesthetize the animal before every behavioral session which preserve the animal welfare and do not alter the animal’s behavioural performance.

Before performing fUS imaging in a trained rat, the rat needs to be acclimated to the new behavioural box in which the imaging will occur and to the weight on its head. In order to do this, incremental weighs are put over the mock probe (every session it is increased 2 grams up to 10 grams) while the animal is training in the new behavioural box where the imaging is going to take place. As soon as the rat reach a performance comparable to its behavioural performance before the surgery, we are ready to perform fUS imaging.

At the time of the experiment, the mock probe is removed. Then sterile de-bubbled ultrasonic gel (Sterile Aquasonic 100 Ultrasound Gel) is put on the actual probe. The real ultrasound probe is then inserted into the probe holder and fixed in place tightly. Before the probe is inserted, the functional ultrasound system is turned on. In order to synchronize the functional ultrasound imaging sequence and the behavior that the animal is performing, we employed a custom designed system composed of an arduino that receives a TTL signal from the ultrasound imaging machine and the behavioural control box and send it to an Open-Ephys recording box. We then use the Open-Ephys software to synchronize those signals. We have validated the synchronization to make sure that both the imaging sequence and the behavioral events are aligned.

We used the same imaging procedure and sequence we used in (Takahashi et al., 2021). The hemodynamic change measured by fUS strongly correlates with the cerebral blood volume (CBV) change of the arterioles and capillaries. It compares more closely to CBV-fMRI signal than BOLD-fMRI (Macé et al., 2018). CBV signals show a shorter onset time and time-to-peak than BOLD signals (Macé et al., 2011). fUS signals correlate linearly with neural activity for various physiological regimes (Nunez-Elizalde et al., 2021). We used a custom ultrasound linear probe with a minimal footprint (20 mm by 8 mm) and light enough (15 g) for the animal to carry. The probe comprises 128 elements of 125 *µ*m pitch working at a central frequency of 12 MHz, allowing a wide area coverage up to 20 mm depth and 16 mm width. The probe was connected to an ultrasound scanner (Vantage 128, Verasonics) controlled by an HPC workstation equipped with 4 GPUs (AUTC, fUSI-2, Estonia). The functional ultrasound image formation sequence was adapted from (Urban et al., 2015). The main parameters of the sequence to obtain a single functional ultrasound imaging image were: 200 compound images acquired at 500 Hz, each compound image obtained with 9 plane waves (−6^*°*^ to 6^*°*^ at 1.5^*°*^ steps). With these parameters, the fUS had a temporal resolution of 2 Hz and a spatial resolution (point spread function) of 125 *µ*m width, 130 *µ*m depth, and 200 *µ*m to 800 *µ*m thickness depending on the depth (200 *µ*m at 12 mm depth). We acquired fUS signal from the rat brain at the +1 mm lateral sagittal plane in this study.

### 2.4 Brain clearing and registration

#### 2.4.1 Perfusion

After the endpoint of each rat, we perfused the animal with ∼100-150 ml (or until clear) 1xPBS mixed with 10 U/ml heparin (cold). It was then perfused with ∼200 ml 4% PFA in 1xPBS. Then the descending artery was clamped, and the body was tilted by 30^*°*^ towards the head. The animal was next perfused with 50 ml fluorescence-tagged albumin hydrogel (recipe below). The animal’s head was immediately buried in ice for 30 min. The brain was extracted and further submerged in 4% PFA at 4^*°*^ C overnight. The brain was then processed using iDISCO clearing protocol (Renier et al., 2014).

Hydrogel recipe: Make 50 ml 2% (w/v) gelatin (G6144 Sigma-Aldrich) in PBS, warm up to 60^*°*^ C and filter once. Add 5 mg AF647-albumin (A34785 Invitrogen), mix well. Store in 40^*°*^ C water-bath till use.

#### 2.4.2 Light-sheet microscopy

The brain was cut along the midline into two hemispheres (Sup. Fig. 8). A brain hemisphere was immobilized with super glue in a custom cradle with the medial side facing up. This yielded the best quality towards the midline. The cradle was placed in the chamber filled with DBE of a lightsheet microscope (LaVision Biotech). We used the 488 nm channel for autofluorescence imaging and the 647 nm for vasculature imaging. The sample was scanned with a 1.3x objective, which yielded a 5 um x 5 um pixel size and with a z-step of 5 um. Two light sheets were adopted, and dynamical horizontal focusing was applied. We set a low NA and low dynamical focusing for the autofluorescence channel and a high NA high dynamical focusing for the vasculature channel. Each sample was covered by two tiles with overlap. Preprocessed images combining the dynamical foci were saved.

#### 2.4.3 Registration

We registered fUS images to a standard brain atlas via the alignment of brain vasculature. The registration procedure took two steps: one registered the autofluorescence channel of light-sheet images (taken in the same coordinate reference system as the brain vasculature) to a standard brain atlas (Step 1), and one registered a reference fUS image to a plane of vasculature light-sheet image from the same subject (Step 2). We detail the steps as follows.

Step 1: light-sheet to the rat brain atlas The light-sheet image stacks were first down-sampled to 25 *µ*m isotropic (or a voxel size matching the atlas). The autofluorescence channel of each brain hemisphere was then registered to the corresponding hemisphere of the brain atlas2 (e.g., a highresolution MRI template) by sequentially applying rigid, affine, and B-spline transformation using Elastix3 (parameter files in folder ‘registrationParameters LightSheetToAtlas’). The transformations estimated in this step were applied to the vasculature channel. The two hemispheres were then assembled, thereby registering the brain vasculature to the atlas.

Step 2: fUS to light-sheet To localize the fUS position in the brain, we created a 2D vascular atlas by taking the max-intensity projection over 300 *µ*m thickness along the midline of the registered vasculature volume.

We chose a reference session for each subject, calculated the mean intensity at the logarithmic scale, and converted it to an unsigned 8-bit image. The non-brain region above the sagittal sinus was masked out to avoid false signals. This fUS image served as a reference for other sessions to align with and for aligning to the subject-specific vascular atlas.

The fUS image of the same subject was aligned with this vasculature reference using a landmark-based 2D registration procedure (ImageJ plugin bUnwarpJ4). The matching locations were manually identified by the shape of the big vessels present in both images. To avoid overfitting, we imposed small weights on the divergence and curl of the warping. The generated transformation file can be used to warp any fUS results to the atlas.

### 2.5 Timing of trials

Assume that a trial runs for time *T* and that the fUS signal is collected at sampling rate *f*_*s*_. For a particular frequency in the signal to have a fraction 1*/L* of the wavelength within a trial. Thus the slowest frequency observable by this criterion *f*_*min*_ must satisfy

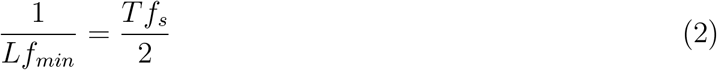

where *Tf*_*s*_*/*2 is the number of effective time-points in the trial given the Nyquist frequency. For *L* = 4 and *f*_*s*_*/*2 = 1 Hz, this reduces to *f*_*min*_ = 1*/*(4*T*), which, for the trial lengths of *T* = 0.5, 3, and 5 s correspond to *f*_*min*_ = 0.5, 0.083, and 0.05 s.

### 2.6 Burst error analysis

Burst errors are defined as sudden, large increases in movie intensity over the entire field-of-view. Such events can be caused, for example, by sudden movements of the animal. Consequently, these events are simple to detect by analyzing the total intensity of the frames, as measured by the *l*_2_ norm 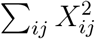 for each frame *X*. We dynamically select a cut-off for determining a burst frame by estimating the inflection point in the histogram of frame-norms loosely based off of Otsu’s method (Otsu, 1979).

### 2.7 Motion Correction

To ascertain the accuracy of the motion correction algorithm, we estimated the residual motion shift using a sub-pixel motion estimation algorithm based on fitting a Laplace function to the autocorrelation *C*_*ij*_ = ⟨*X*_*ref*_, *X ∗ δ*(*x − τ*_*i*_, *y − τ*_*j*_) ⟩, where *X*_*ref*_ is the z-scored reference image *X* is the z-scored image with the offset we wish to compute. The Laplace function is parameterized as

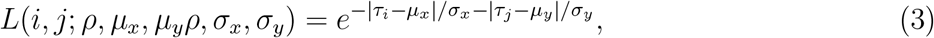

where and {*ρ, µ*_*x*_, *µ*_*y*_, *σ*_*x*_, *σ*_*y*_} is the parameter set for the 2D Laplace function, including the scale, x-shift, y-shift, x-spread and y-spread respectively. We optimize over this parameter space more robustly by operating in the log-data domain, i.e.,

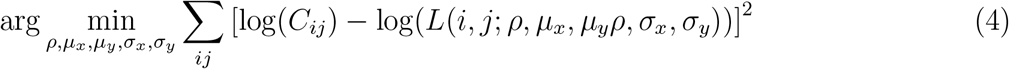

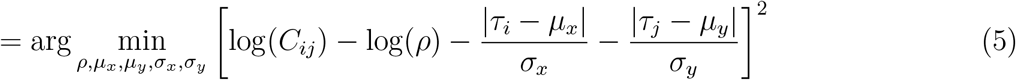

which makes the optimization more numerically stable, gradients are easier to compute and solving for log(*ρ*) directly removes one of the positivity constraints from the optimization. We further reduce computational time by restricting the correlation range to only shifts of |*τ*_*i*_|, |*τ*_*j*_| *<* 25 pixels and by initializing the optimization to the max value of the cross-correlation function {*i, j*} = arg max(*C*_*ij*_).

This computation relies on a global reference. If few frames are shifted, than the median image of the movie serves as a reasonable estimate. In some cases this is not possible, and instead other references, such as the median image of a batch of frames at start or end of the video. We also provide the option to estimate the motion residual over blocks of frames, sacrificing temporal resolution of the estimate for a less sensitivity to activity. In the case of fUS, the temporal resolution sacrifice is minimal due to the initial round of motion correction from the built-in processing steps.

Optimization for motion correction was performed using constrained interior-point optimization via MATLAB’s built-in fmincon.m function, constraining the Laplace parameters to be positive (*λ*_*x*_, *λ*_*y*_ *>* 0), and the offset values to be within the pre-defined limits (− *µ*_max_ ≤*µ*_*x*_, *µ*_*y*_ *≤µ*_max_).

### Motion correction assessment

Assessing motion correction for functional data can be nontrivial, as the individual pixels themselves may change over time along with the shifts in the full image. To assess motion, we leverage the fact that multiple trials repeat in the data, which implies that the per-pixel statistics should be relatively stationary. This observation means that if we divide the data into lowand highmotion frames, the distribution of pixel values (e.g., as a histogram) should be similar. We measure similarity in this case using the Earth Mover’s Distance (EMD) (Ling and Okada, 2007), a metric historically used to measure the similarity between probability distributions.

Regardless of the assumed stationarity, the data may still have local variability in different epochs. We thus would ideally want to compare the measured distribution deviation relative to a baseline. For this reason we further divide the low-motion frames and high-motion frames into multiple segments and measure an ANOVA-style ratio of the variability between segments in different classes to the variability within each class. Mathematically we divide the values of each pixel at location *i, j* into groups *G*_*low,k*_ and *G*_*high,k*_^*′*^ for *k* = 1, …, *K* and *k*^*′*^ = 1, …, *K*^*′*^, i.e., *K* blocks of low-motion frames and *K*^*′*^ frames of high-motion frames. The groups *G* aim to include contiguous blocks in order to capture local statistics. We also define the distances *d*_*EMD*_(*G*_1_, *G*_2_) denotes the EMD between the normalized histograms of groups *G*_1_ and *G*_2_. The metric we use to determine motion impact on the data is then

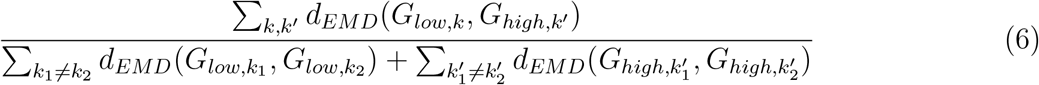

A high value for this ratio indicates that the high motion frames follow much different statistics and therefore motion will likely impact the results for any inference using that pixel. As a comparison, we shuffled the data blocks into random groupings and compute the shuffled score for each pixel. The resulting baseline distribution was concentrated in the range [0.5, 2]. Stable data had similar statistics, while data from recordings with larger jumps often resulted in values up to three times greater. We find that motion correction largely reduces these values, however for data with very large jumps the correction does not always result in a perfect baseline when compared to the shuffle test.

This metric provides one useful measurement of motion impact on a pixel-by-pixel bases. Thus it permits assessing local motion effects. Global motion effects can be measured using a second observation: significant motion adds nonlinear variability to the data, which broadens the eigenspectrum of the data. Specifically we can look at the data’s principal component singular values and compare the curves with and without motion correction.

### 2.8 Time-trace denoising

To denoise time-traces, we apply standard wavelet-based denoising methods available through MATLAB. Wavelet denoising depends on a number of parameters, including the wavelet class and level at which to denoise (i.e., the time-scale of the finest-resolution details kept). We find that for functional ultrasound data using the ‘sym4’ wavelets at level 6 resulted in adequate denoising.

### 2.9 Differential map computation

Differential maps compare activated areas under two separate conditions, e.g., in and out of trials or activity during correct or incorrect trials. Assume each frame *i* can belong to either condition Γ_1_, Γ_2_ or neither (Γ_1_ Γ_2_)^*C*^ The maps were computed in two stages. First the frames *Y*_*i*_ corresponding to conditions 1 *i* ∈Γ_1_ and 2 *i* ∈Γ_2_ were separated and averaged separately to obtain an average map for each condition. Second the difference between these two maps were computed. Together the difference map is

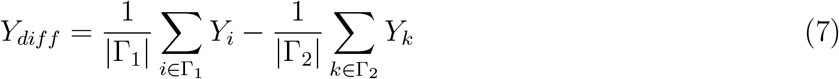

### 2.10 Correlation map computation

Correlation maps measure which image pixels tend to be more or less active during specific events, such as the start of a trial or the delivery of reward. To compute these maps, the weighted average of frames proceeding and succeeding each of the *L* events at frame *τ*_*l*_ is computed, using weights *w* ∈ R^2*K*^ . The average of the weighted average over all events then forms the correlation map.

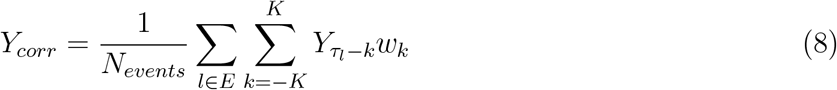

This computation is equivalent to computing the temporal cross-correlation of a kernel *w* ∈ R^2*K*^ with a spike train defined by the event times, and then computing the inner product of the resulting vector and each pixel. To differentiate activity directly before and directly after an event from constant activity, we use a kernel that is negative before the event and positive after. In particular we use a Gaussian kernel *w*_*k*_ = *exp*(−*k*^2^*/*4) multiplied by a shifted Heaviside function that is +1 for *k ≥*0 and −1 for *k <* 0.

We relate the computed correlation to known brain areas by integrating the before (negative) and after (positive) values in the correlation map over the extent of each brain area (Supp. Fig. 10, 11, 12, 13).

Mathematically we compute the sums

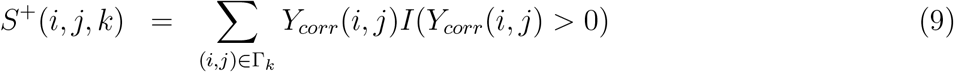

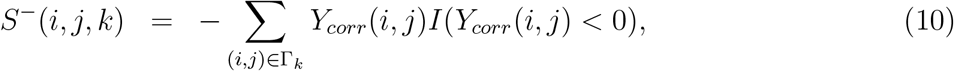

where Γ_*k*_ is the set of pixels in the *k*^*th*^ brain area (as per the registration procedure), and *I*(·) represents the indicator function that is 1 if the condition is true and 0 if false. The negative sign in the second computation is simply to reverse the encoding of “before” in *Y*_*corr*_ as negative values. Finally we normalize all the computed values by the maximum per session *S*_*max*_ = max(max_*ijk*_(*S*^+^), max_*ijk*_(*S*^*−*^)).

As different brain areas can have vastly different sizes (i.e., number of voxels covered), for visualization purposes we find it convenient to transform each value in *S*^+^ and *S*^*−*^ as *S*(*i, j, k*) ← *S*(*i, j, k*)*/*(0.01 + *S*(*i, j, k*)). The resulting two values provides a summary statistic for each brain area’s correlation to that event. Note that the differential nature of the correlation maps indicates that areas exhibiting activity both before and after the event must exhibit that activity in separate sets of voxels.

### 2.11 GraFT analysis

To isolate canonical time-traces from fUS data, Graph Filtered Time-trace GraFT (Charles et al., 2022) analysis was applied to the motion-corrected and denoised video matrix ***Y*** . ***Y*** is a *T ×P* matrix where *T* is the number of frames in the time-series and *P* is the number of pixels. The GraFT decomposition factorizes ***Y*** into two matrices ***Y≈* Φ*Z***: a *T ×M* matrix **Φ** consisting of *M* distinct time-traces and a *M ×P* matrix ***Z*** consisting of *M* spatial profiles indicating how strongly each time-traces is present in a pixel.

Practically, GraFT proceeds in three steps, where the first initializes the learning and the second two are iterated until convergence.

1. First a graph is constructed where each node represents one pixel and the connectivity (represented by the graph adjacency matrix ***W***) is based on the correlation of the pixel time-traces. Specifically we use diffusion maps (Coifman and Lafon, 2006) for this step.
2. Second we use a randomly initialized matrix **Φ** to identify the spatial maps ***Z*** using a graph-filtered reweighted *l*_1_ (GF-RWL1) procedure (Charles et al., 2022). GF-RWL1 represents a variational inference over ***Z*** where a second set of variables *γ* correlate the values of ***Z*** along the graph. Mathematically we iterate over the following two computations until convergence (practically 2-3 computations suffice):

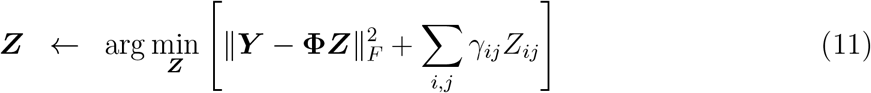

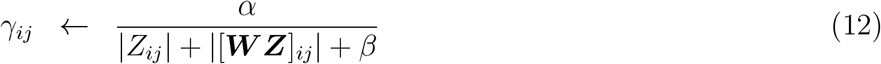

where *α* and *β* are model parameters and [***W Z***]_*ij*_ represents the {*i*.*j* }^*th*^ element of the graph values prjected once through the graph adjacency matrix ***W*** .
3. Third the time-traces are updated given the new spatial maps as

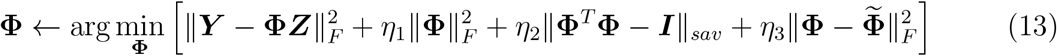

where *η*_1_, *η*_2_, and *η*_3_ are optimization trade-off parameters and Φis the previous estimate of Φ. This optimization is composed of four terms: the first enforcing data fidelity, i.e., that the decomposition should match the data, the second penalizing unused time-traces to automatically prune extra components, the third to constrain the time-traces away from being too correlated, and finally the fourth induces a slowness to the learning over iterations that stabilizes the full time-trace learning procedure.

The initial number of components is pre-set as *M* . For fUS data we set *M* = 30, 45, 60 and found minor improvements on decoding and no significant change in variance explained.

### 2.12 Decoding of behavioral variables

Decoding of behavioral events is performed after pre-processing and the video decomposition using GraFT. As an example we describe here the procedure for decoding inand outof trial frames, however the same procedure was used for all decoding, replacing frames in both categories with appropriately partitioned data. To create target labels *z*_*i*_, we label the in-trial frames as *z*_*i*_ = −1 and out-trial frames as *z*_*i*_ = −1. Since the variation of the signal always reflects more information than a static snapshot, we use a moving window of the last *N* − 1 frames to decode the frame class (e.g., in/out of trial). Additionally, in order to reduce the impact of noise and computational complexity, the decoding is performed on the reduced dimensionality provided by the GraFT timetraces (see previous methods section). We demonstrate this process with linear decoding as an example. Defining the target as a linear combination of the last *N* frames ***Y***_*i*_, *i* = 1, …, *N* we have

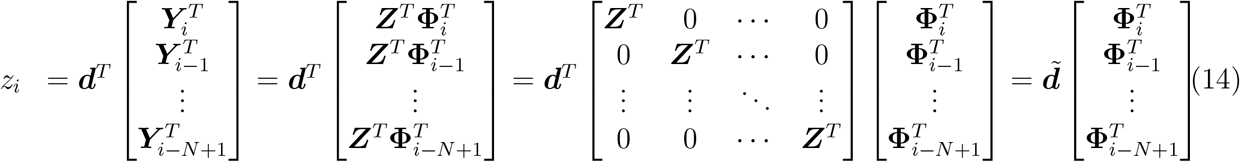

where we use the fact that each data image ***Y***_*i*_ can be defined as a linear combination of the spatial maps ***Z*** with weights **Φ**_*i*_ *∈* R^*M*^, as per the GraFT decomposition. The dimensionality of the initial decoder ***d*** can be reduced from *NP* to *NM* . More generally, the data vectors [**Φ**_*i*_, **Φ**_*i−*1_, …, **Φ**_*i−N*+1_]^*T*^ can be classified as per their labels *z*_*i*_ using other methods. In particular we applied both a Linear SVM and a Gaussian Kernel SVM in the decoding.

Two main parameters influence the decoding procedure are the number of past frames used in the decoding *N*, and the number of GraFT components *M* . We compared the decoding results of different *N* and *M* . *N* we swept from *N* = 1 (i.e., using only the current frame) to *N* = 20. In some cases, for example in decoding correct and incorrect trials, *N >* 10 exceeded the trial length, so a maximum of *N* = 5 was used. When decoding the trial difficulty, since the interval between two trials is always 7 frames in our data, a maximum of *N* = 7 was used to guarantee a good generalization ability of the decoder.

The second parameter *M* that influences decoder performance actually a parameter of the decomposition. The number of GraFT components *M* modifies the decoding performance based on the overall information and potential complexity of the reduced space. In our experiments, *M* was set to *M* ∈ {30, 45, 60}. We find that when *N* is small, larger *M* leads to a little higher precision, however this difference decrease as *N* becomes larger.

In general, we find that the non-linear SVM always performs better than the linear SVM. When applying a Gaussian Kernel SVM, there are two hyperparameters connected with its performance. One is the scale of the Gaussian kernel, and another is the box constraint, which controls the maximum penalty imposed on margin-violating observations. Using the function *fitcsvm* in MATLAB, and setting the input argument *‘OptimizeHyperparameters’* as *‘auto’*, we automatically find the optimal hyperparameters via five-fold cross-validation.

### 2.13 Spurious correlations due to motion

One factor to consider when quantifying the effect of motion is the possibility of spurious correlations across an image, as seen in fMRI (Power et al., 2012). To illustrate the problem, consider four pixels *p*_1_(*t*), *p*_2_(*t*), *p*_3_(*t*), *p*_4_(*t*), where *p*_1_(*t*) and *p*_1_(*t*) are neighboring and *p*_3_(*t*) and *p*_4_(*t*) are neighboring. Furthermore, assume for the moment that all pixel time-traces are independent, but with different mean values, i.e., E[*p*_*i*_(*t*)] = *µ*_*i*_. As all pixels are independent, *ρ*(*p*_*i*_, *p*_*j*_) = 0 for all pairs of pixels. We specify Γ ⊂ [1, …, *T*] as a set *K < T* times wherein shifts due to motion (or unstable motion correction) effectively mix neighboring pixels. In this case we can now specify

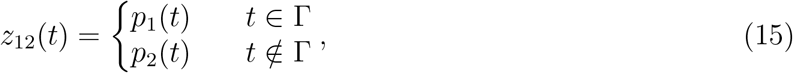

and similarly

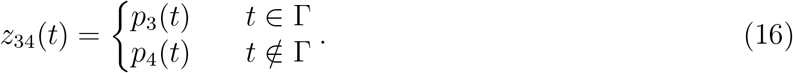

Note that the set Γ specifies the same times for both *z*_12_(*t*) and *z*_34_(*t*) as the motion is global. We can now compute the correlation between the motion-influenced pixels *z*_12_(*t*) and *z*_34_(*t*).

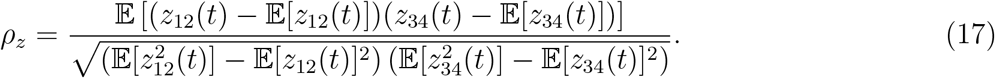

First note that the sample average

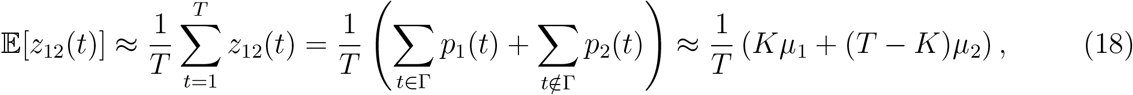

and similarly for *z*_34_(*t*). The sample co variance is then

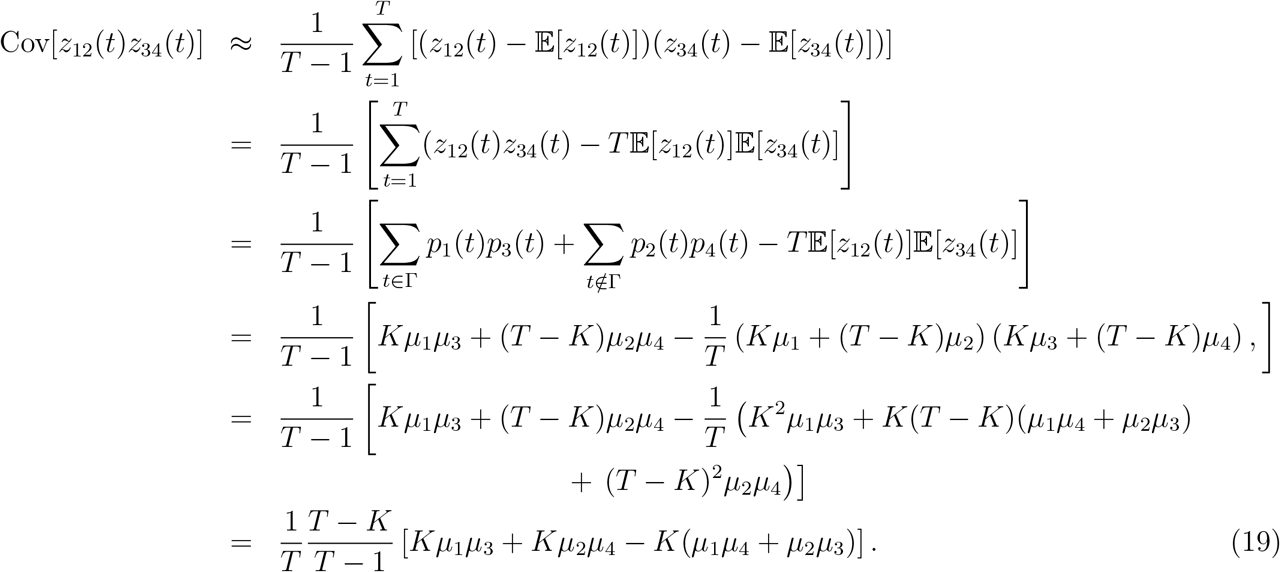

We can immediately see that in a trivial case of *µ*_2_ = *µ*_4_ = 0 and *µ*_1_ = *µ*_3_ = *b*, we have a non-zero covariance 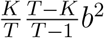 *b*^2^. Similarly for *µ*_1_ = *µ*_4_ = 0 and *µ*_2_ = *µ*_3_ = *b*, we have a non-zero covariance 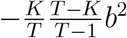. Depending on the sample size *T*, these values may be small—depending on the exact value of the non-zero pixel’s amplitude *b*—but significant.

To compute the full correlation, we also need the denominator of *ρ*_*z*_, which depends on the sample vairances of *z*_12_(*t*) and *z*_34_(*t*).

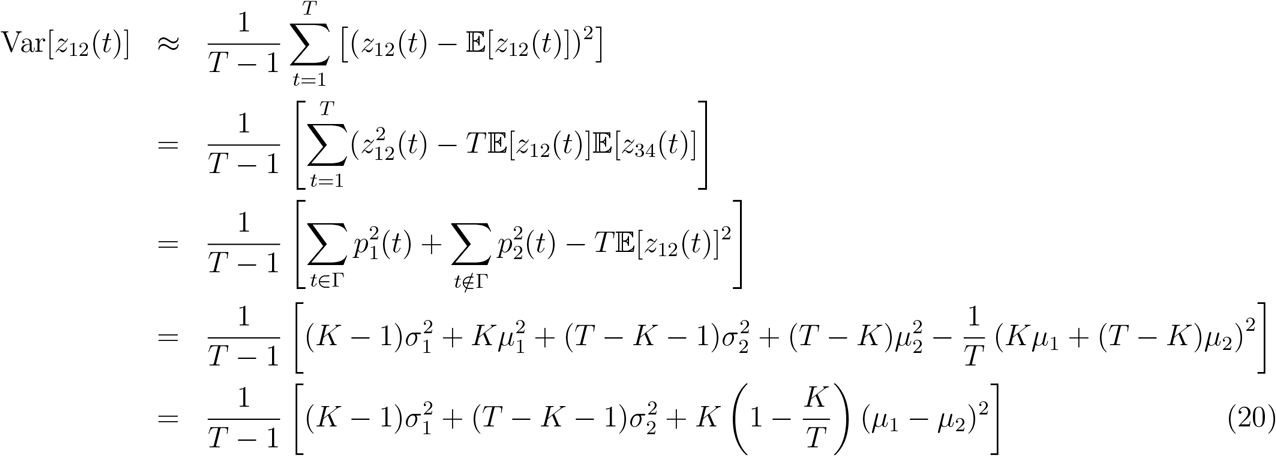

and similarly for *z*_34_(*t*). Note that the variance is essentially the weighted average of the variances of *p*_1_(*t*) and *p*_2_(*t*), with an additional factor depending on the difference-of-means.

The full correlation can now be written, however for previty we assume the following: *µ*_1_ = *µ*_3_, *µ*_2_ = *µ*_4_, and *σ*_1_ = *σ*_3_ = *σ*_2_ = *σ*_4_ = *σ*.

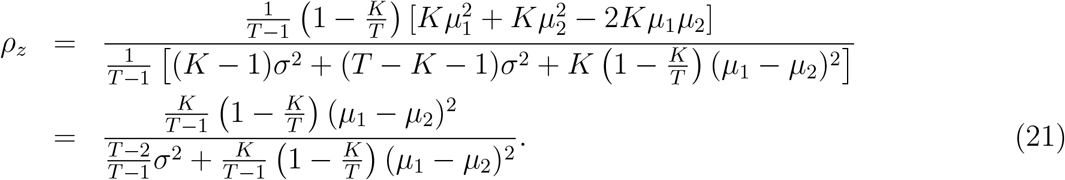

This expression emphasizes a critical point: as the variances of the independent signals shrink with respect to the difference of means, motion can induce a correlation that grows with respect to the fraction of motion-affected frames.

### 2.14 fUS code-base

Reproducible results are a cornerstone of scientific progress. Moreover, practical and available resources are critical to enhancing the capabilities of labs seeking to implement and use new technologies. We thus created an object-oriented code-base in MATLAB that contains all the methods discussed here and serves as a foundation for future advances for fUS methods. The code is written to be easily used, expandable, and capable of analyzing both single and multiple recordings, i.e., from chronic implantation as described in this work.

