## Supplementary Figures for "Chronic brain functional ultrasound imaging in freely moving rodents performing cognitive tasks"

### 3 Supplementary Figures

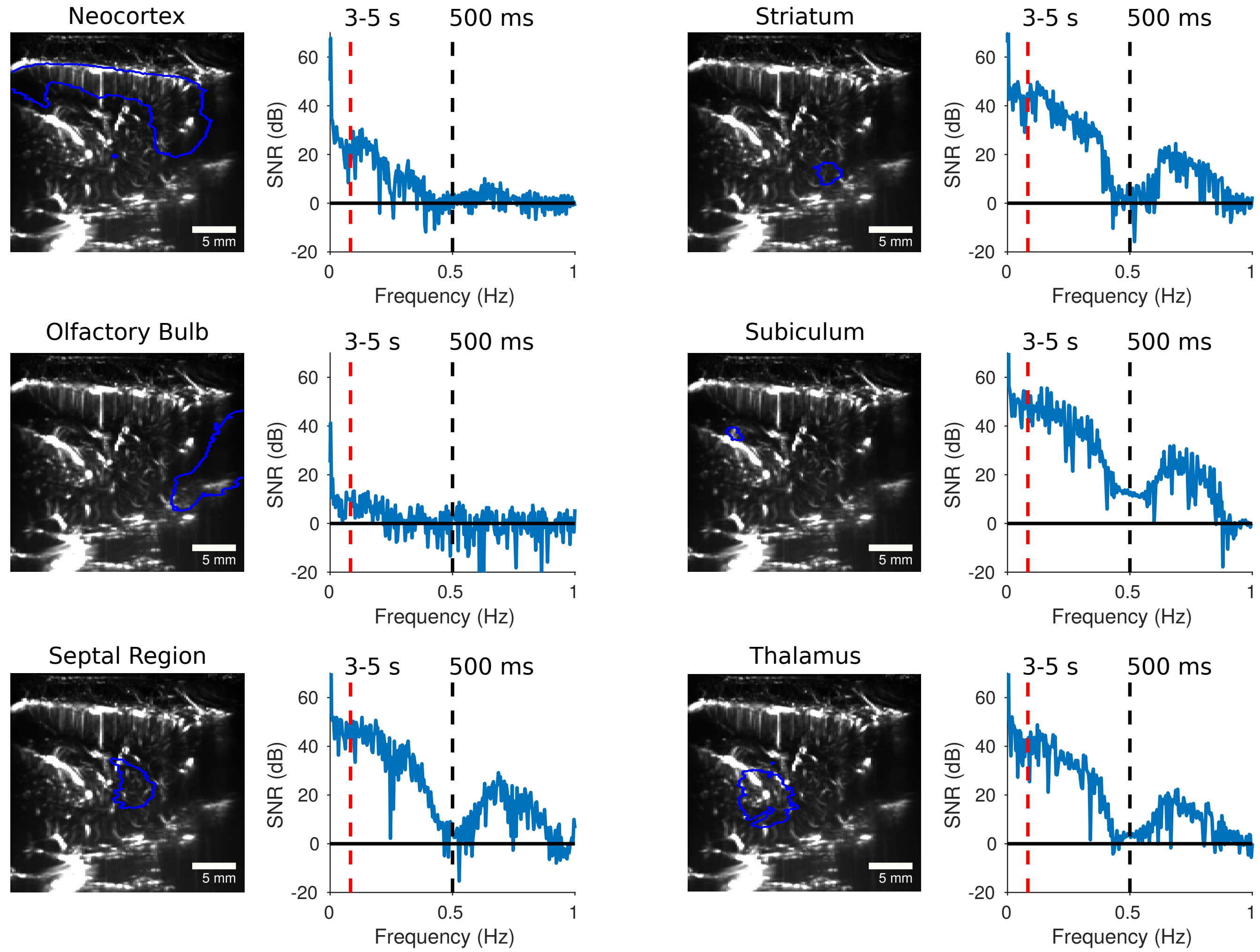


Supplementary Figure 1: Additional examples of fUS frequency content as compared to the observable signal given long (3-5 s) and short (500 ms) trials. Signals are shown for a variety of brain areas, as registered to a standard atlas.

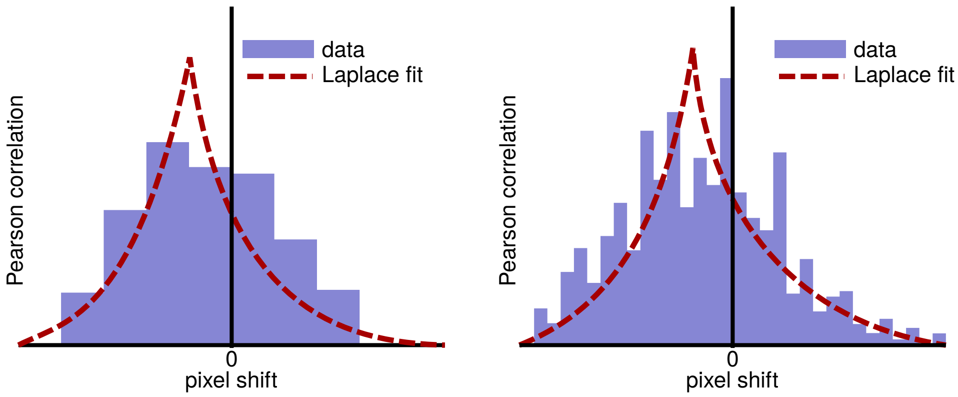


Supplementary Figure 2: Schematic figure of motion estimation using Laplacian fits to the cross-correlation function. Low spatial resolution (left) or high noise levels (right) can create uncertainty in sub-pixel motion estimation when looking for the maximum value in a spatial autocorrelation function (displayed as bar plots in each plot to capture the pixelated nature of the computed function). We fit a Laplace function (red dashed lines) to the autocorrelation function to capture the ideal peaky behavior of image correlations. The estimated offset of this function then becomes the sub-pixel motion estimate

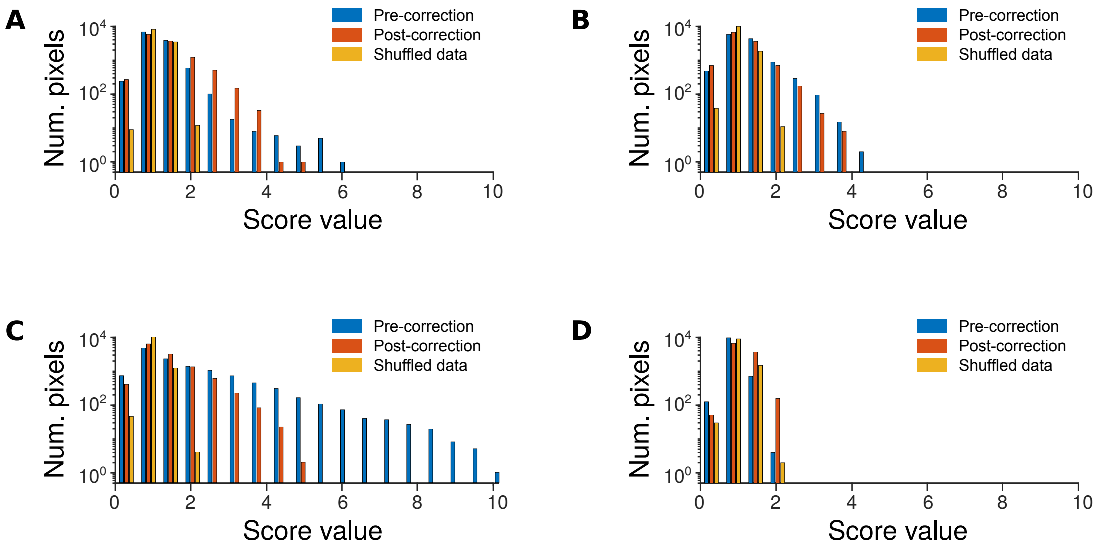


Supplementary Figure 3: Motion metrics for four different fUS recording session spanning two months. Each plot displays the metric values as a histogram over pixels for the raw data (blue), the motion corrected data (red) and the shuffle test (yellow). Well corrected data tended to stay below values of 3, while data with larger jumps could reach values at twice that level. The shuffled data tended to stay below a value of 2, indicating that there are additional correlations in the data that might affect this metric, however overall it enables an "at-a-glance" assessment of motion contamination: if the uncorrected distribution is much broader than the post-corrected distribution, motion correction is likely required.

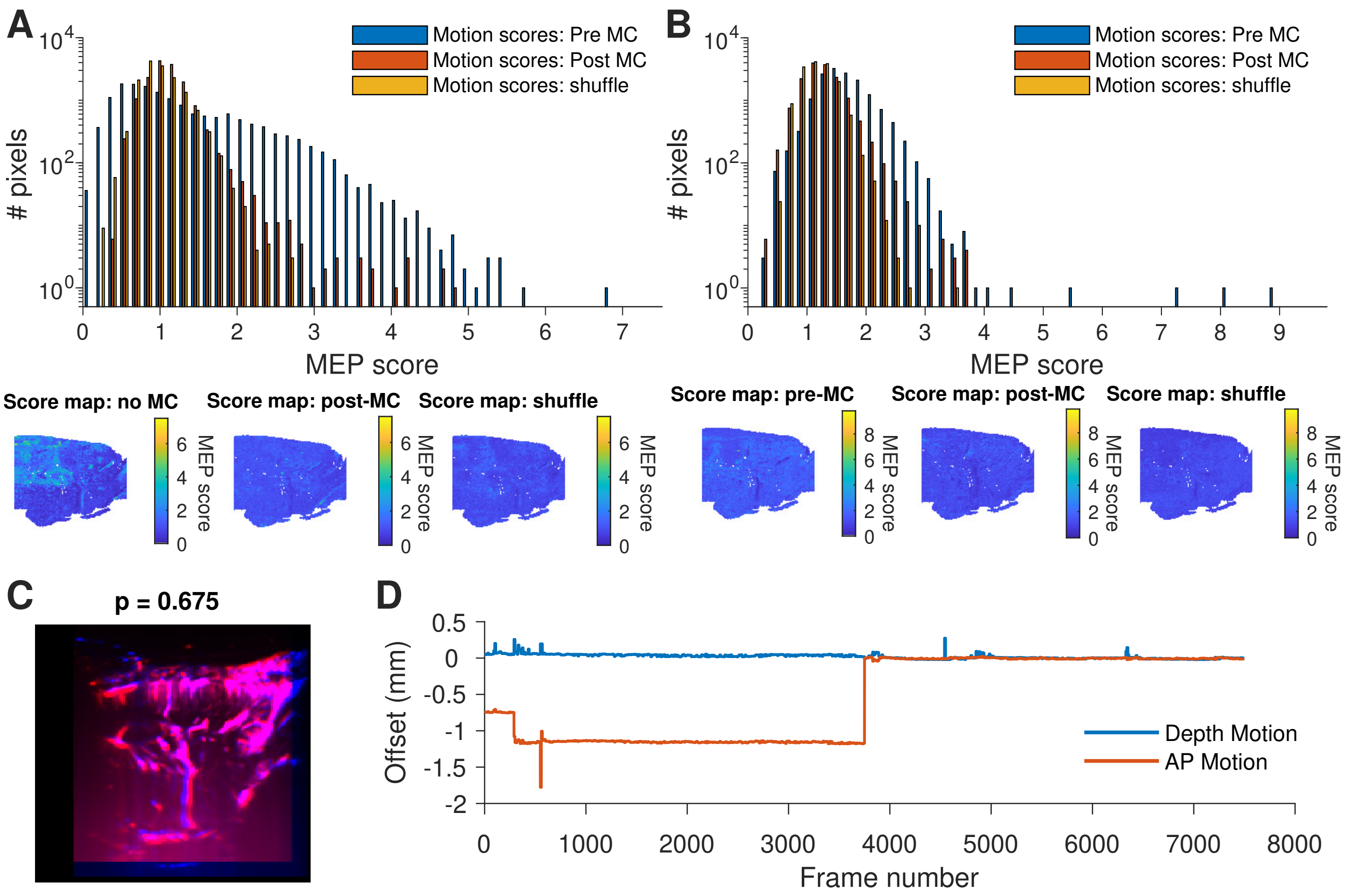


Supplementary Figure 4: Example motion correction in rat H124. A-B) MEPS score histograms from two different distributions, C) Example mean images from both datasets overlayed. D) Relative displacement across both datasets.

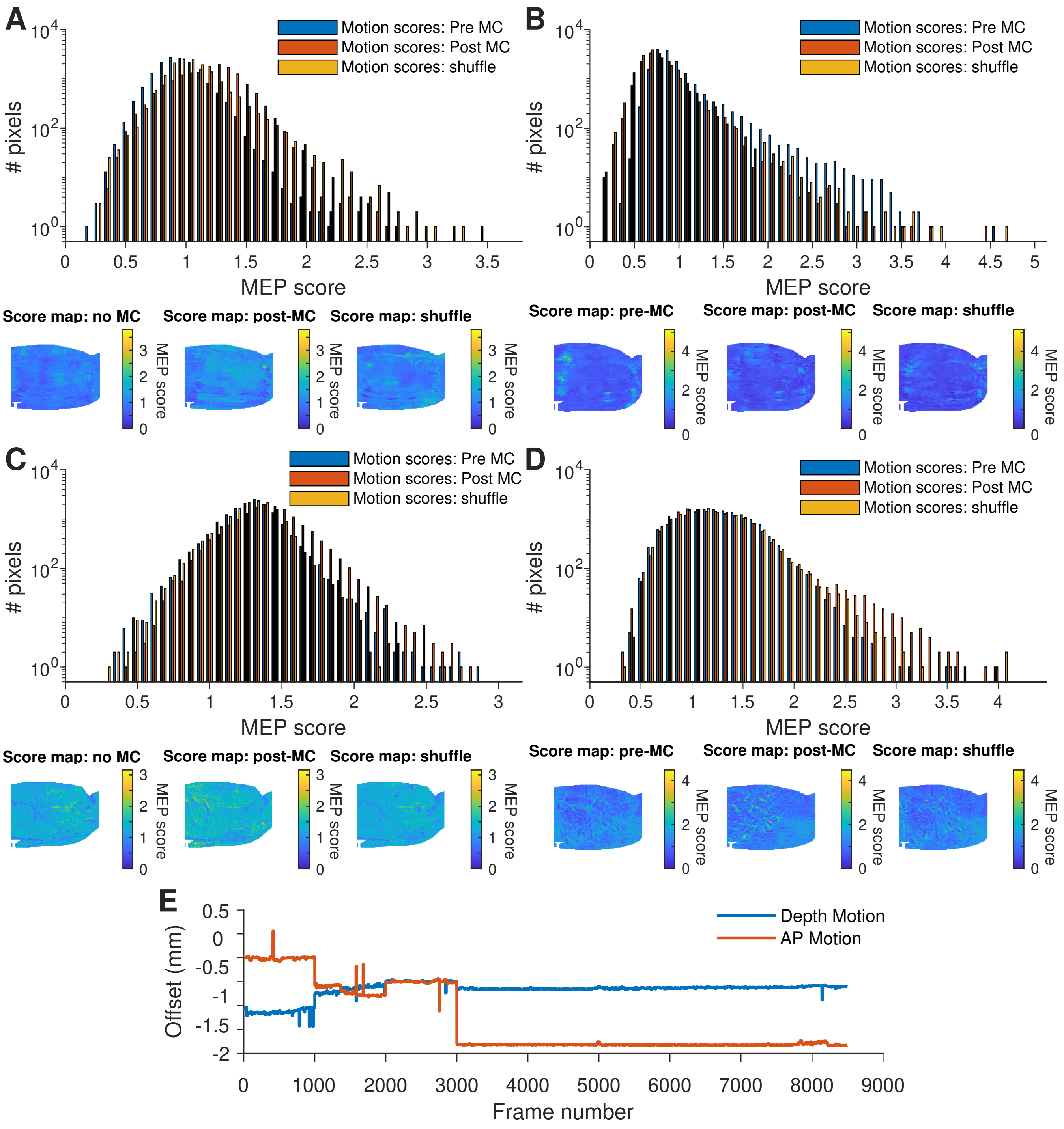


Supplementary Figure 5: Example motion correction in rat H136, set A. A-D) MEPS score histograms from four different distributions, E) Relative displacement across all datasets.

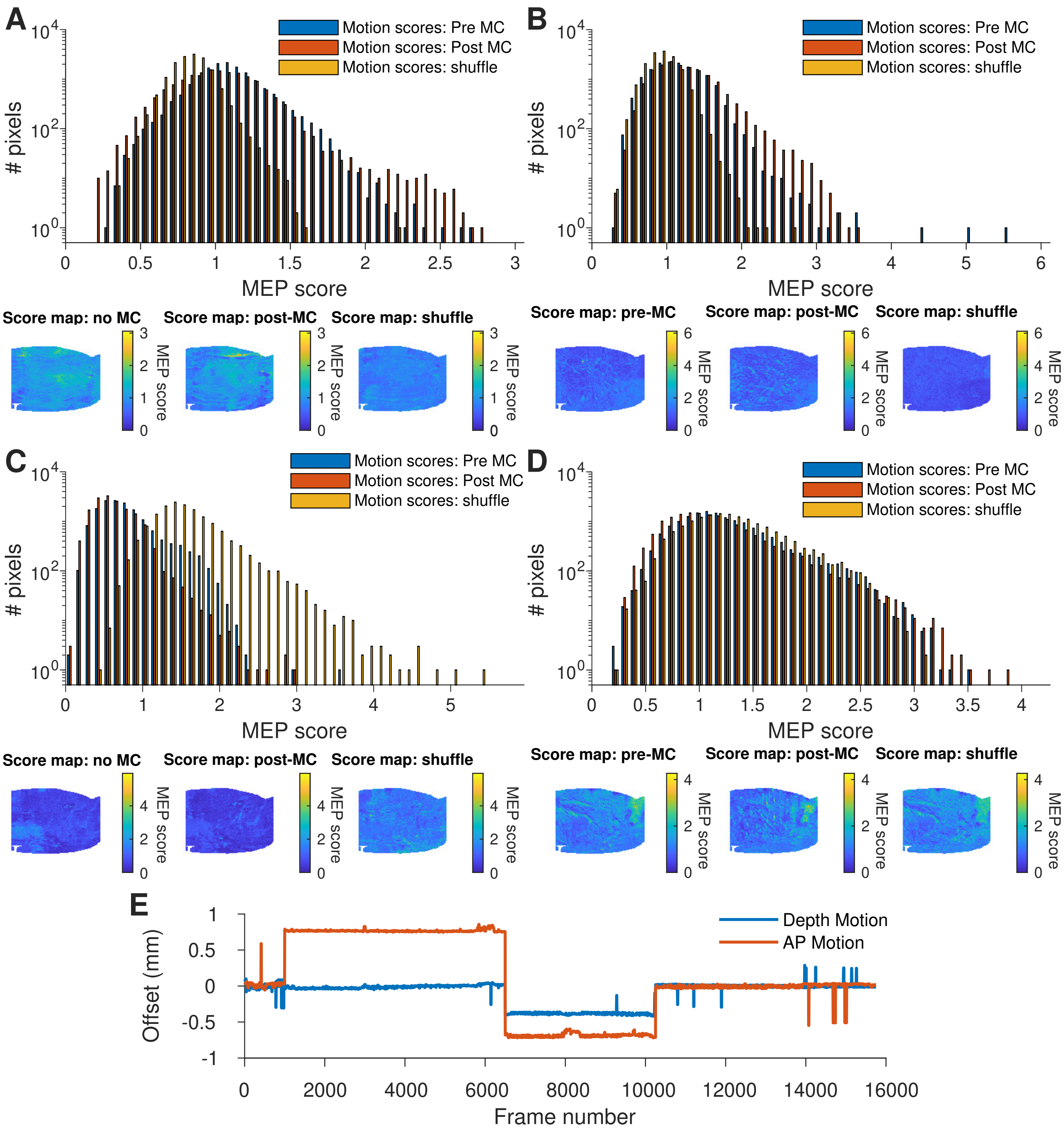


Supplementary Figure 6: Example motion correction in rat H136, set B. A-D) MEPS score histograms from four different distributions, E) Relative displacement across all datasets.

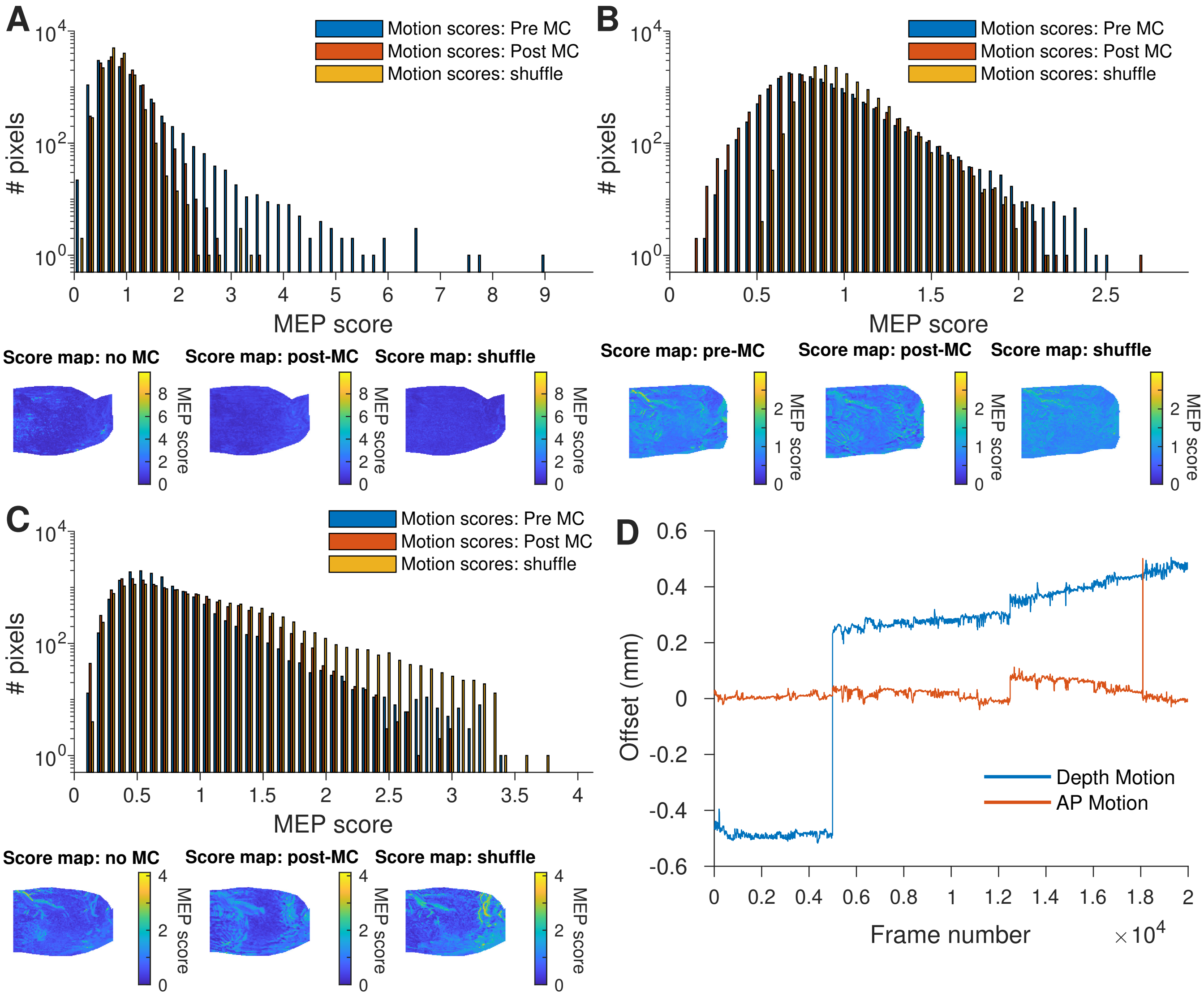


Supplementary Figure 7: Example motion correction in rat H156, set A. A-C) MEPS score histograms from four different distributions, D) Relative displacement across all datasets.

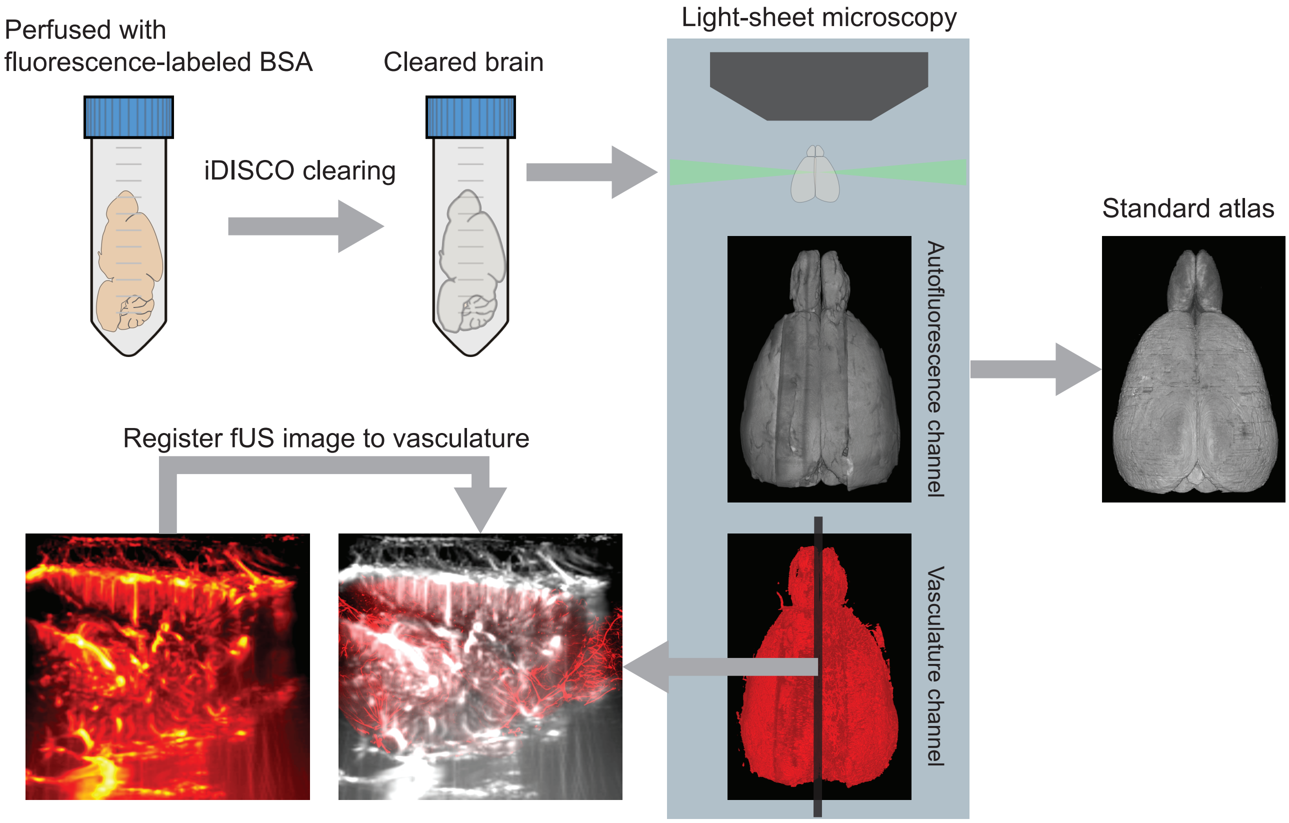


Supplementary Figure 8: fUS images registration: first step is rat brain perfusion followed by iDISCO clearing then light sheet imaging of the cleared brain vasculature. The fUS image is registered to the vasculature image then both fUS and vasculature images are registered to a common standardized rat brain atlas

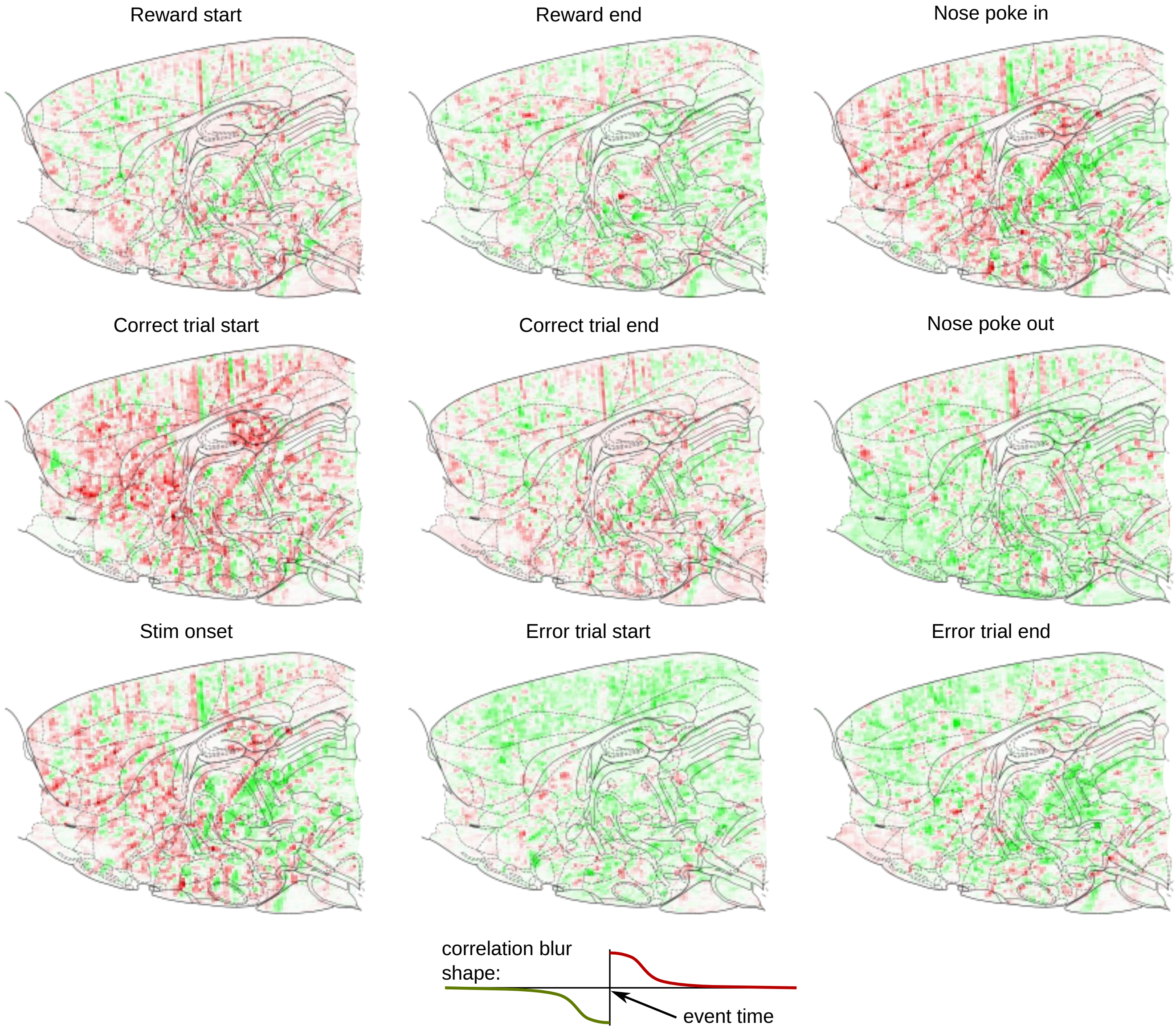


Supplementary Figure 9: Differential correlation images computed for different trial events on the normalized data (each pixel z-scored). Each image displays the correlation image computed by averaging frames with respect to the correlation shape below, where green indicates more activity right before the event, and red indicates more activity right after the event.

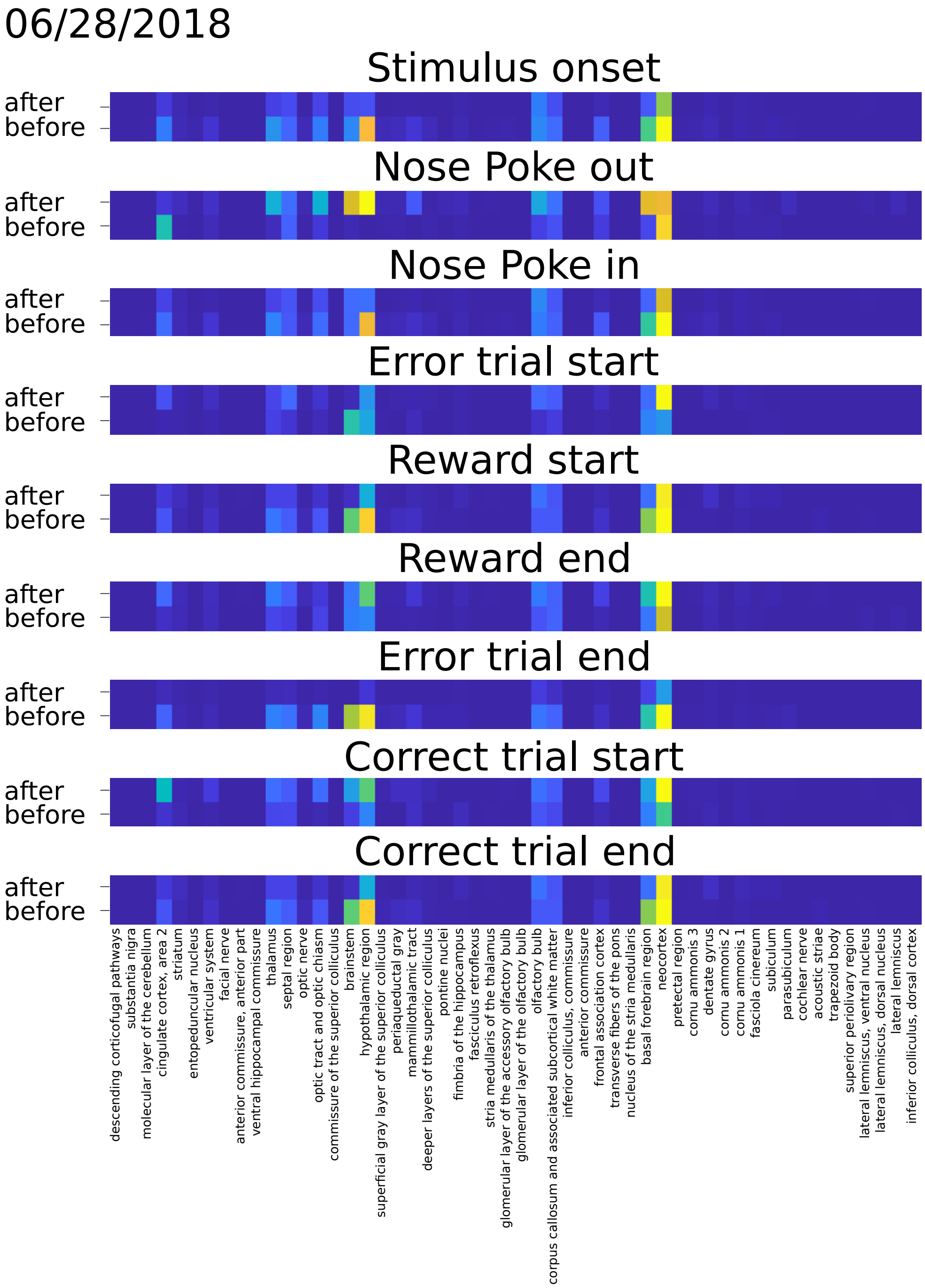


Supplementary Figure 10: Integrated activity in the correlation maps such as shown in Supplementary Figures 9, 26, 27. Activity before and after is integrated over the physical area corresponding to registered atlas components. Areas with activity both before and after indicate separate areas within the same brain area that are active before or after the event, as constant activity is removed by the correlation computation. Activity is shown on a normalized scale, shrunk by $x/(0.01 + x)$ to normalize for the differences in brain area sizes.

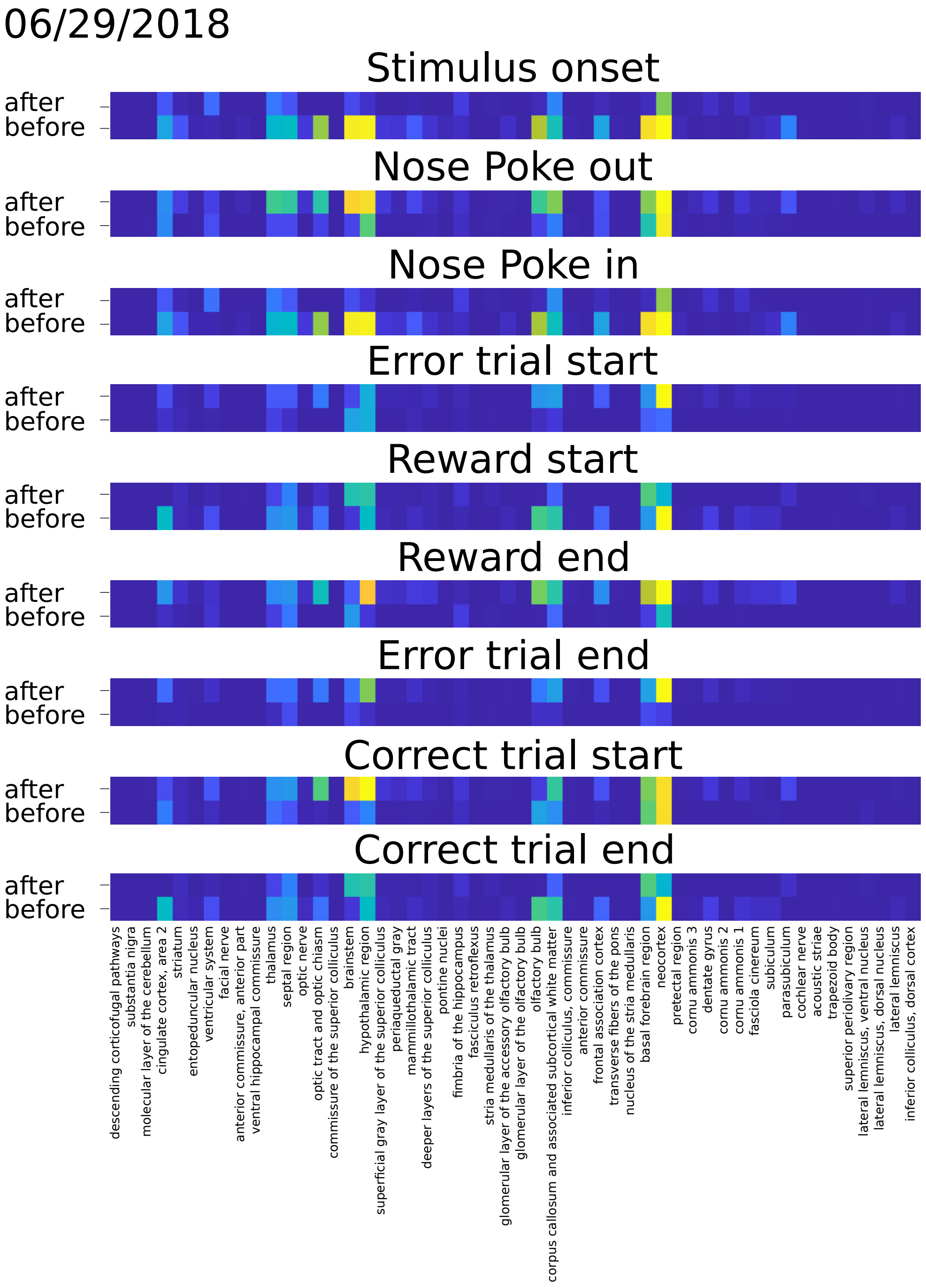


Supplementary Figure 11: Same as Supplementary Figure 10 for the second session.

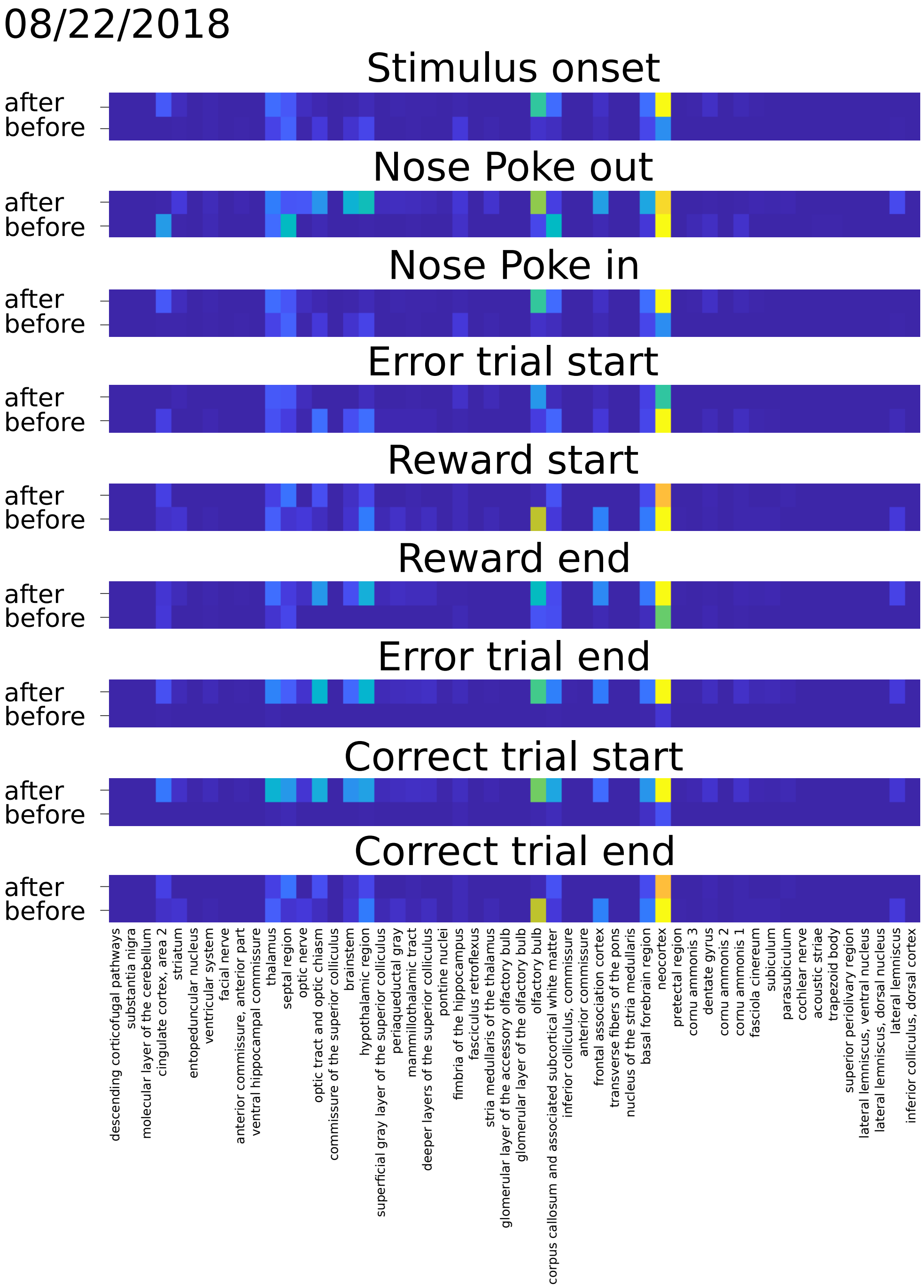


Supplementary Figure 12: Same as Supplementary Figure 10 for the third session.

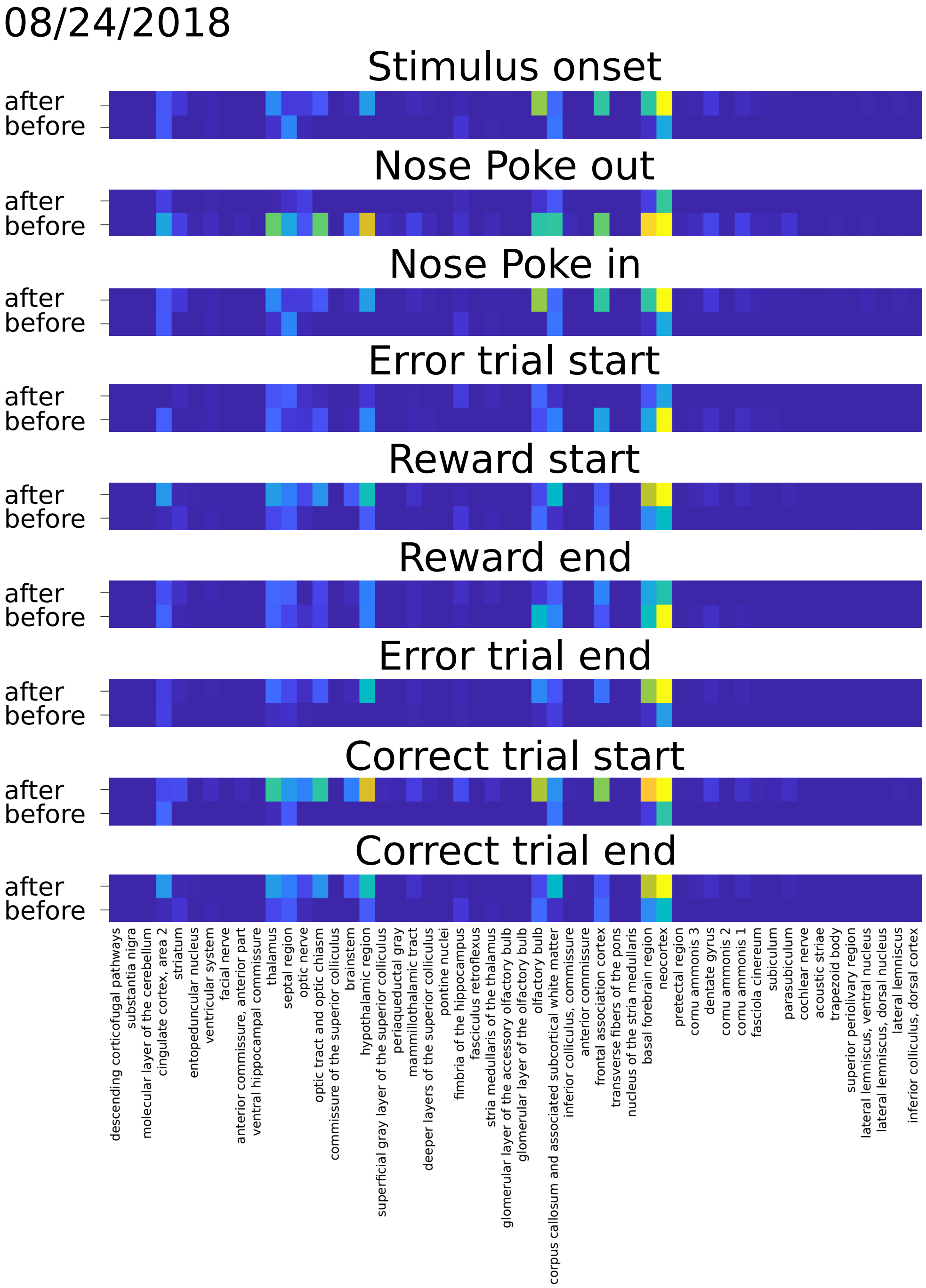


Supplementary Figure 13: Same as Supplementary Figure 10 for the fourth session.

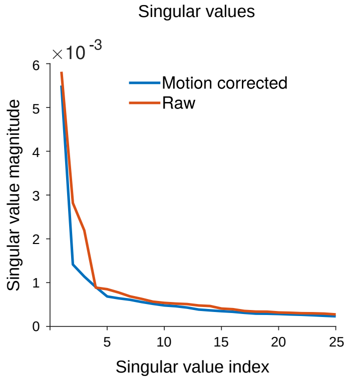


Supplementary Figure 14: The singular value spectrum of the data is reduced post-motion correction. This is an expected result, as translational motion is known to artificially inflate video dimensionality.

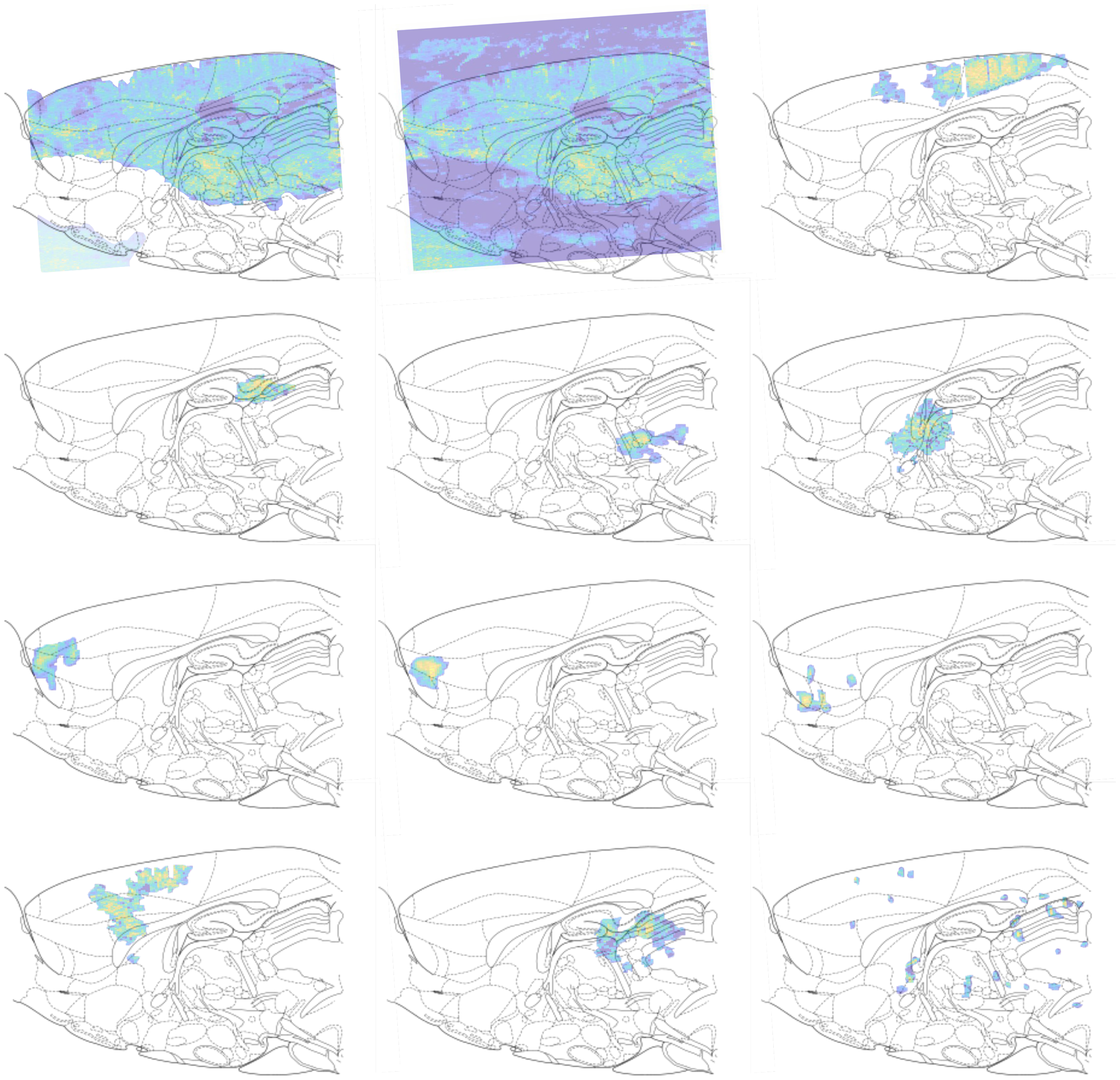

Supplementary Figure 15: Spatial profiles (independently active areas) extracted using completely unsupervised learning on the motion-corrected and denoised fUS data. We note that the first two profiles capture image-wide background levels of activity, while the remaining profiles begin to capture more localized activity pattern. GraFT, in particular, extracts both activity that seems to be well localized within single brain areas, as well as many activity patterns that appear to recruit across brain areas.

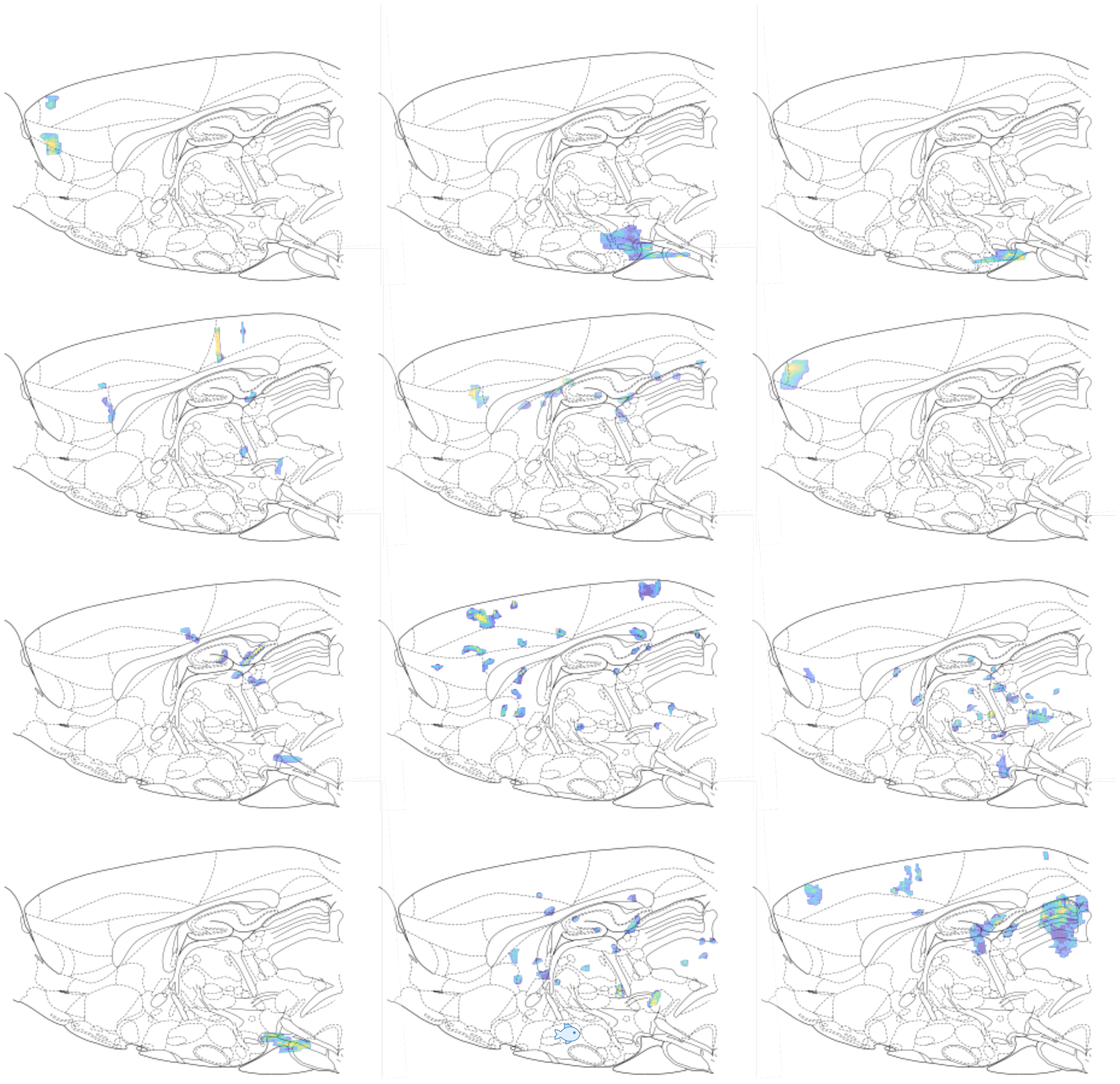

Supplementary Figure 16: (continued from Supp. Fig. 15). Spatial profiles (independently active areas) extracted using completely unsupervised learning on the motion-corrected and denoised fUS data.

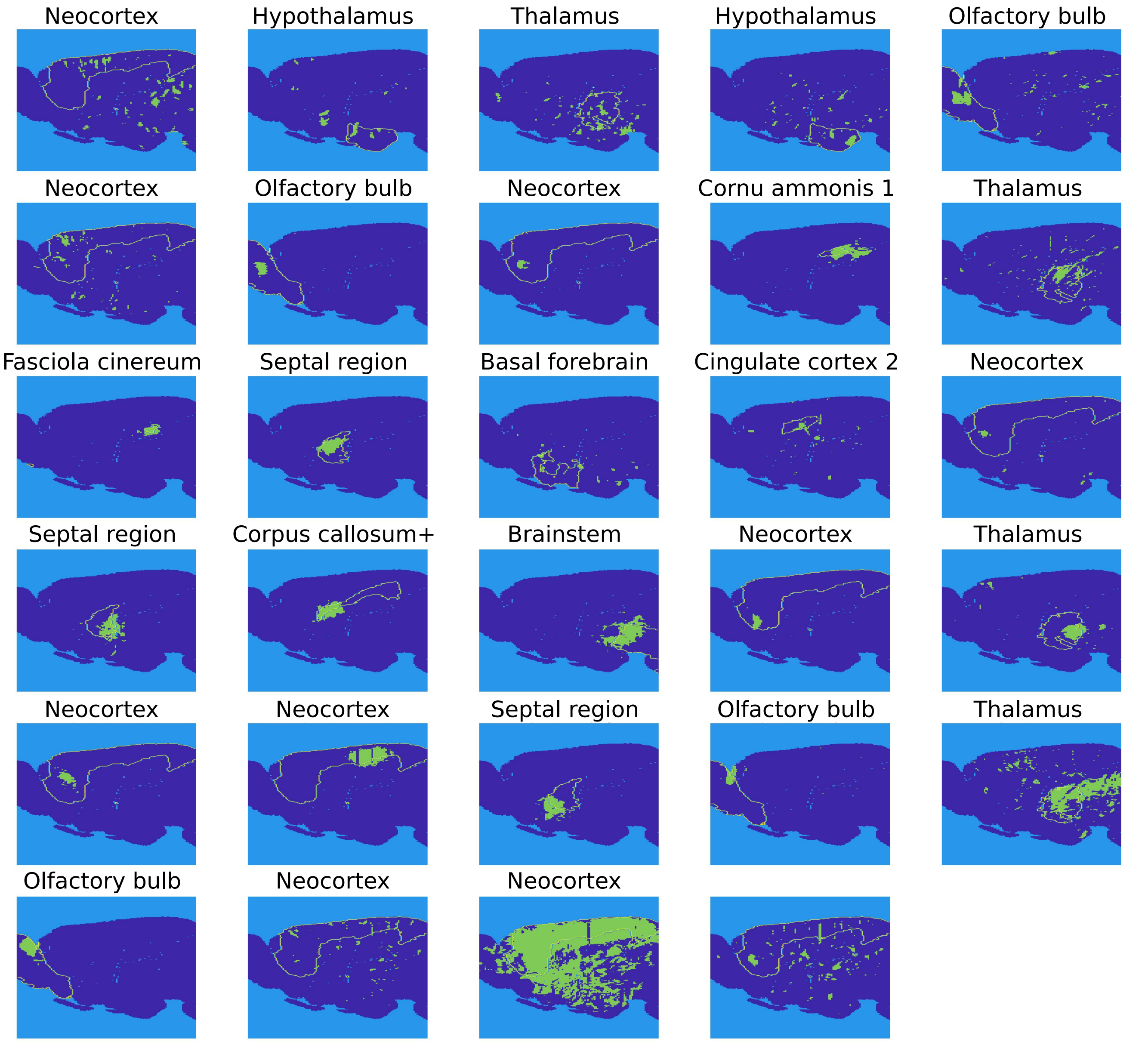


Supplementary Figure 17: Comparisons of spatial profile overlaps (independently active areas) extracted using completely unsupervised learning compared to the brain region with the highest pixel overlap. Titles designate the name of the most overlapping region, outlined in the figure.

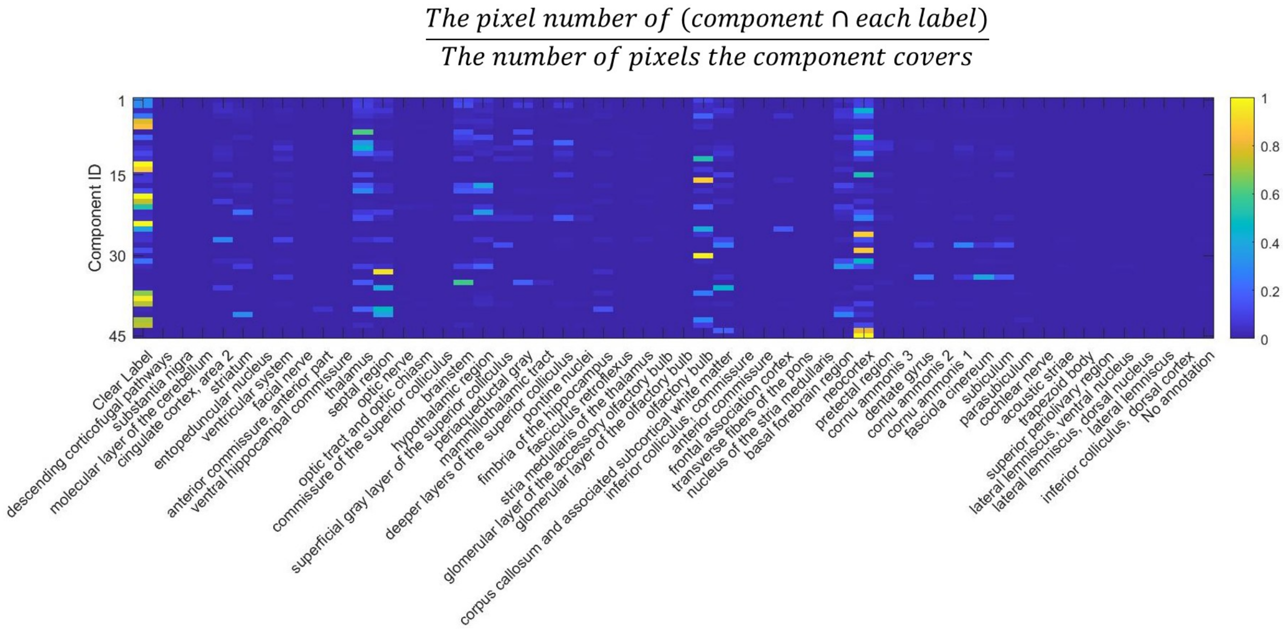


Supplementary Figure 18: Quantification of overlaps between spatial profiles identified via unsupervised learning and anatomically registered brain regions. Color designates percent of pixel overlap between spatial profiles (components in GraFT) and each brain region.

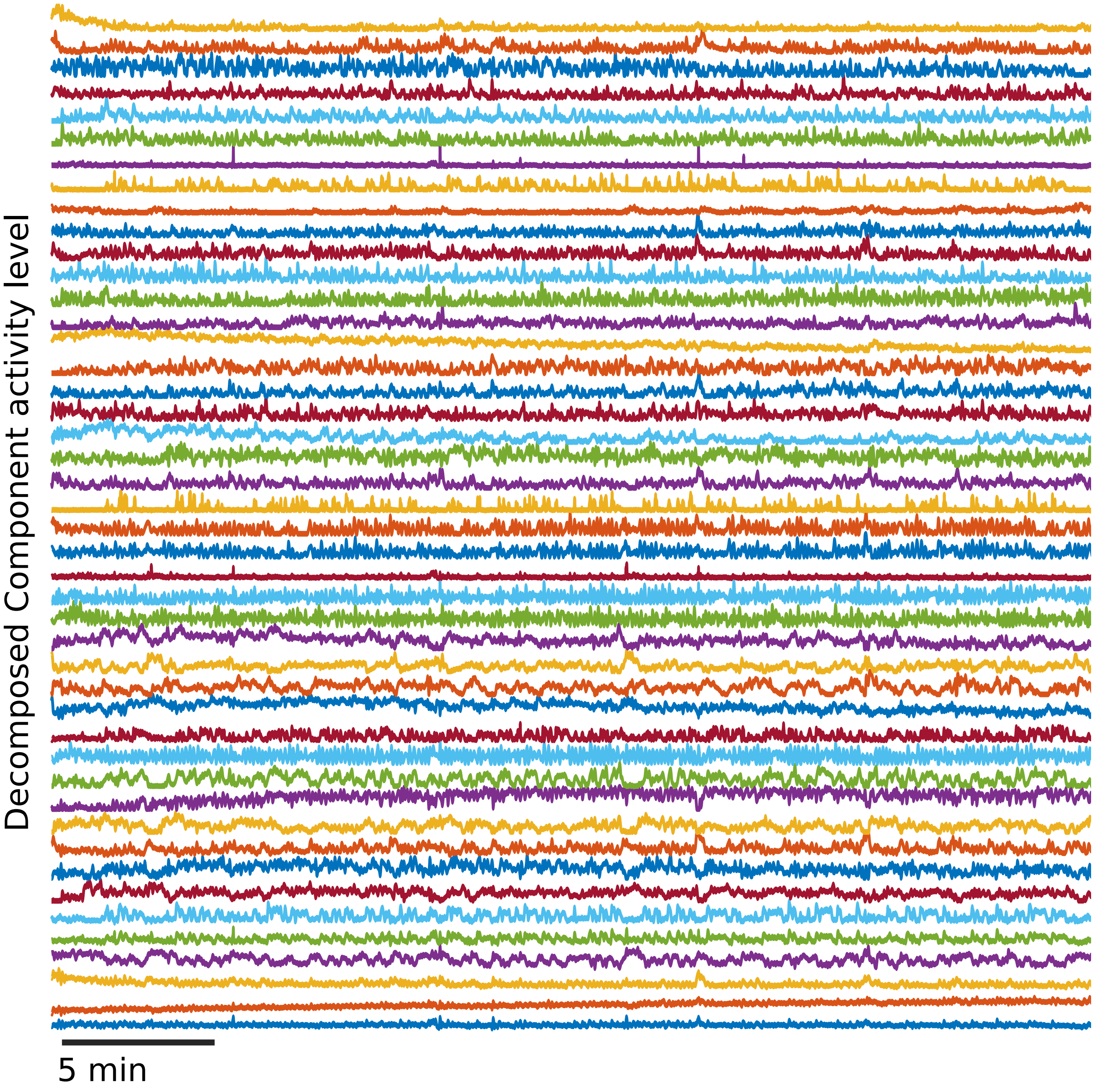


Supplementary Figure 19: Time traces identified in fUS data using GraFT.

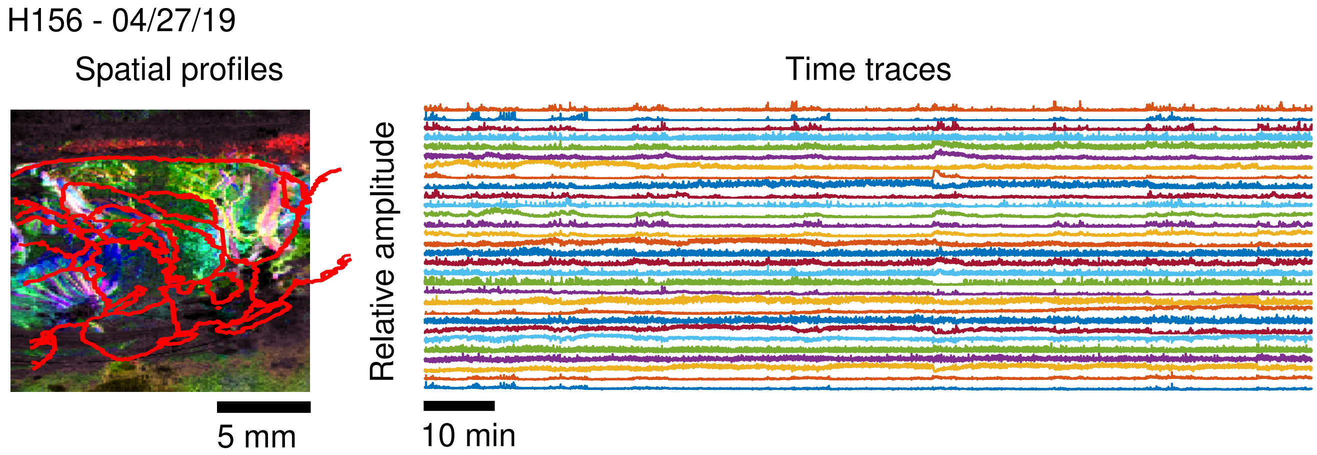


Supplementary Figure 20: Spatial maps and time traces identified in fUS data using GraFT.

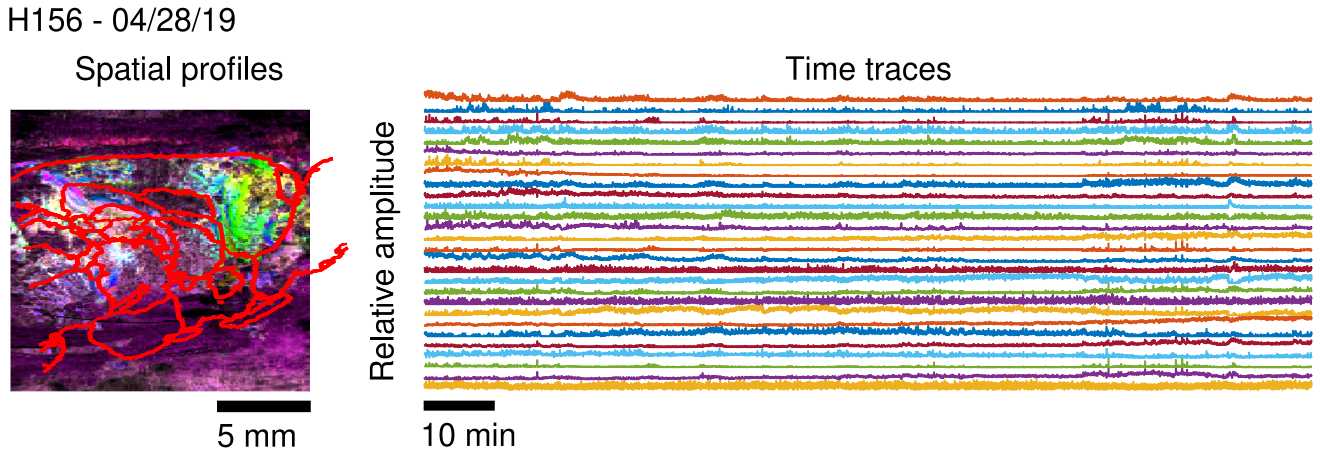


Supplementary Figure 21: Spatial maps and time traces identified in fUS data using GraFT.

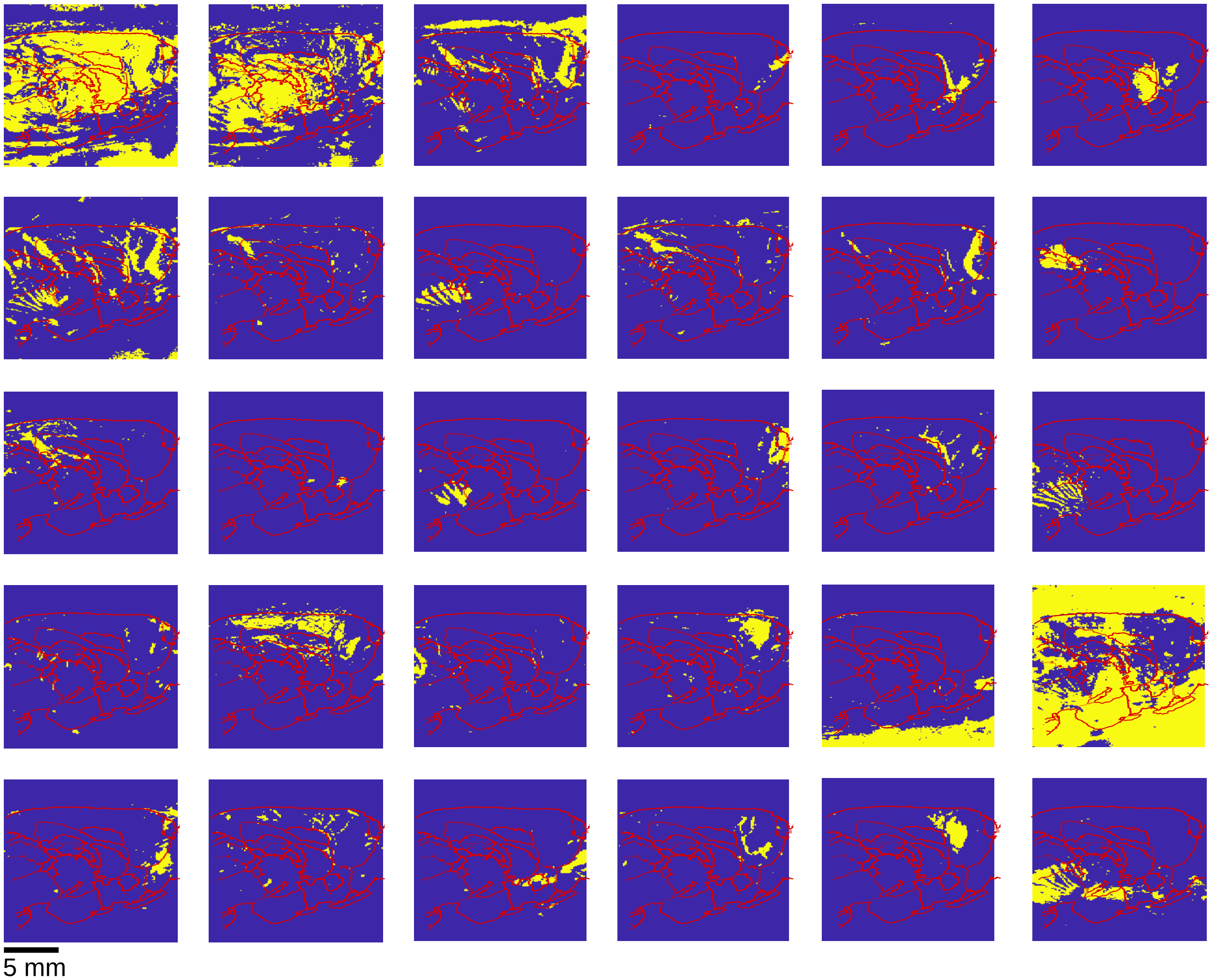


Supplementary Figure 22: Individual spatial maps identified in fUS data using GraFT.

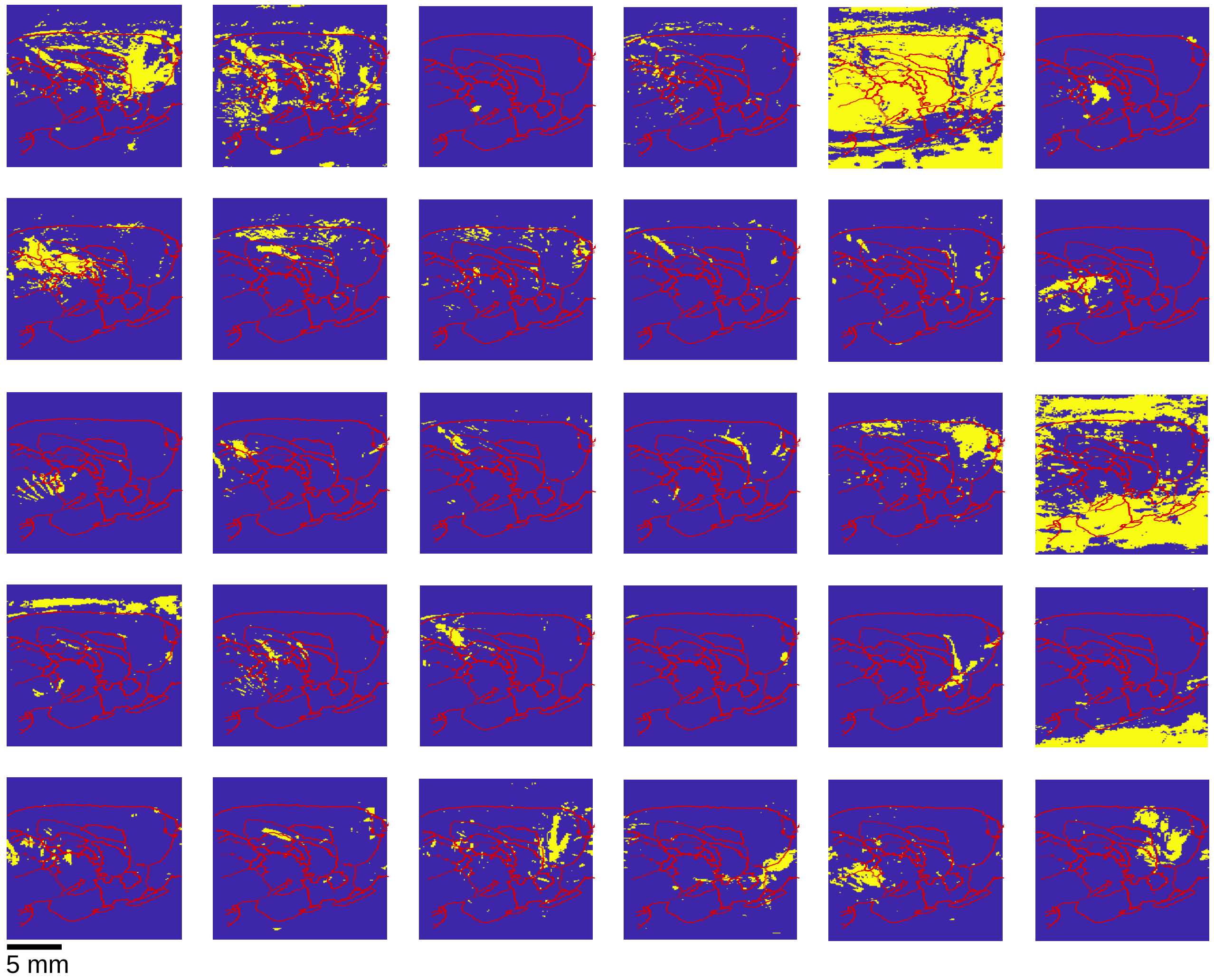


Supplementary Figure 23: Individual spatial maps identified in fUS data using GraFT.

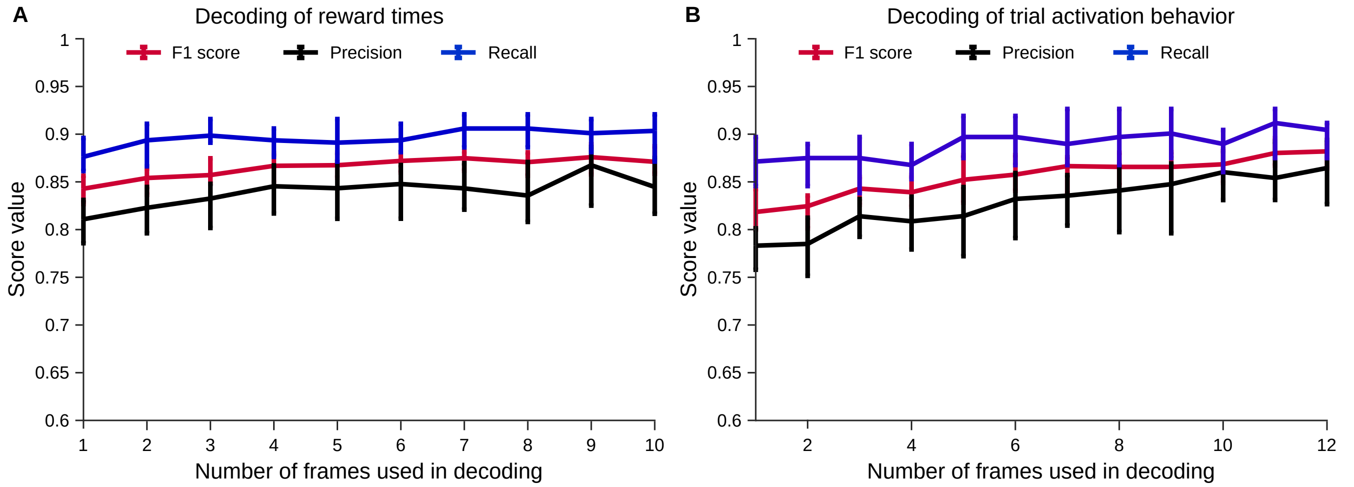


Supplementary Figure 24: Decoding of additional behavioral variables from decomposed fUS data.

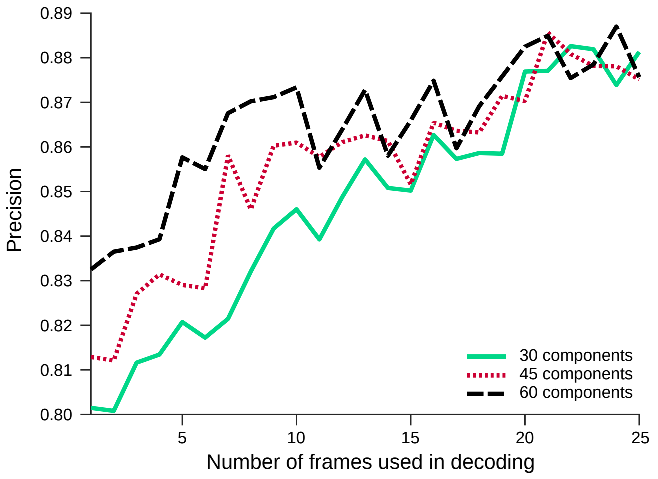


Supplementary Figure 25: Decoding of in/out of trial frames using different numbers of components in the GraFT decomposition. Frame rate is 2 Hz.

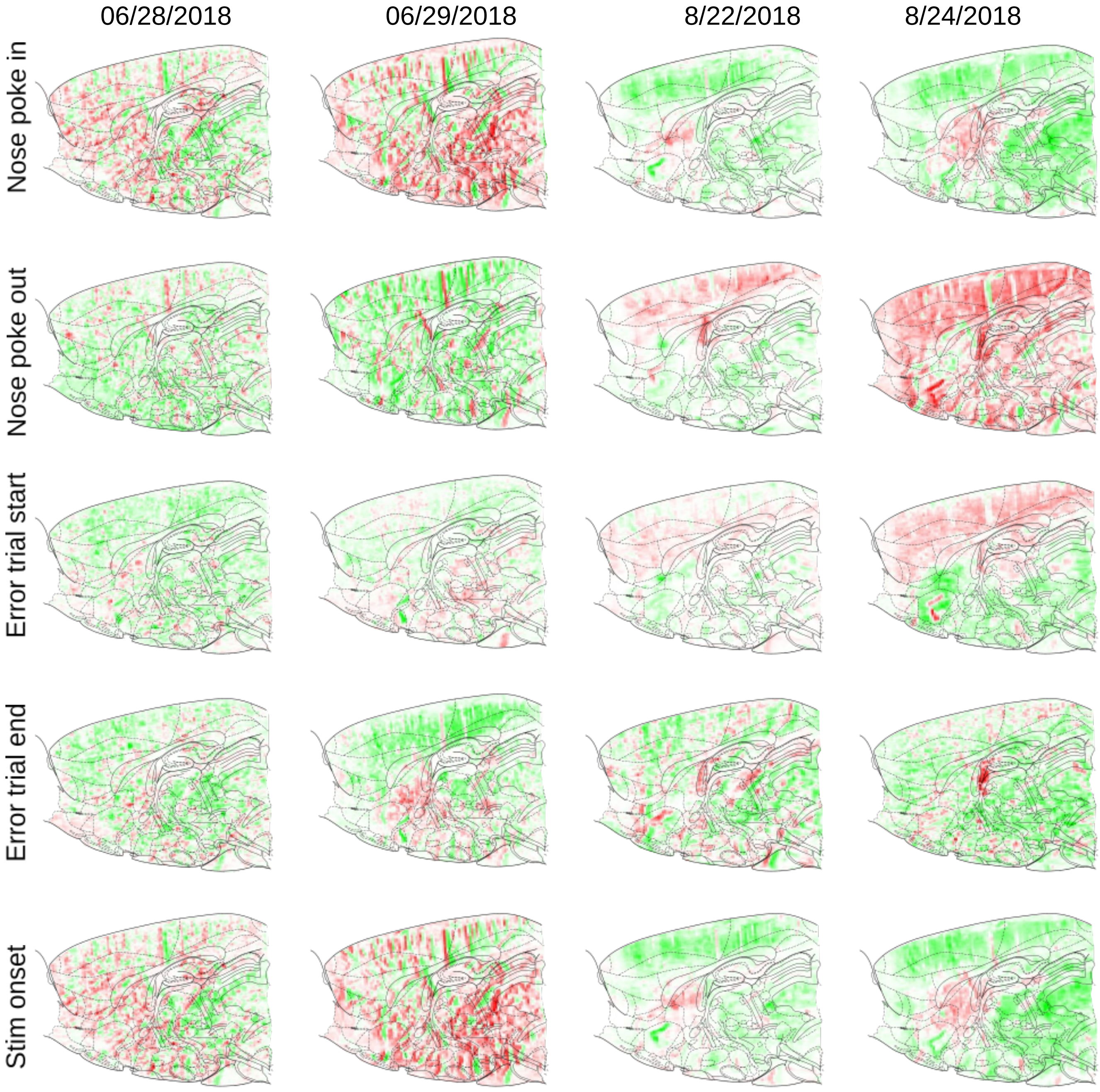


Supplementary Figure 26: Correlation maps for various behavioral events over four sessions spanning two months. Correlations are computed as in Supplementary Figure 9. Rats were fully trained and had, overall, stable behavior by the time of the first session shown here.

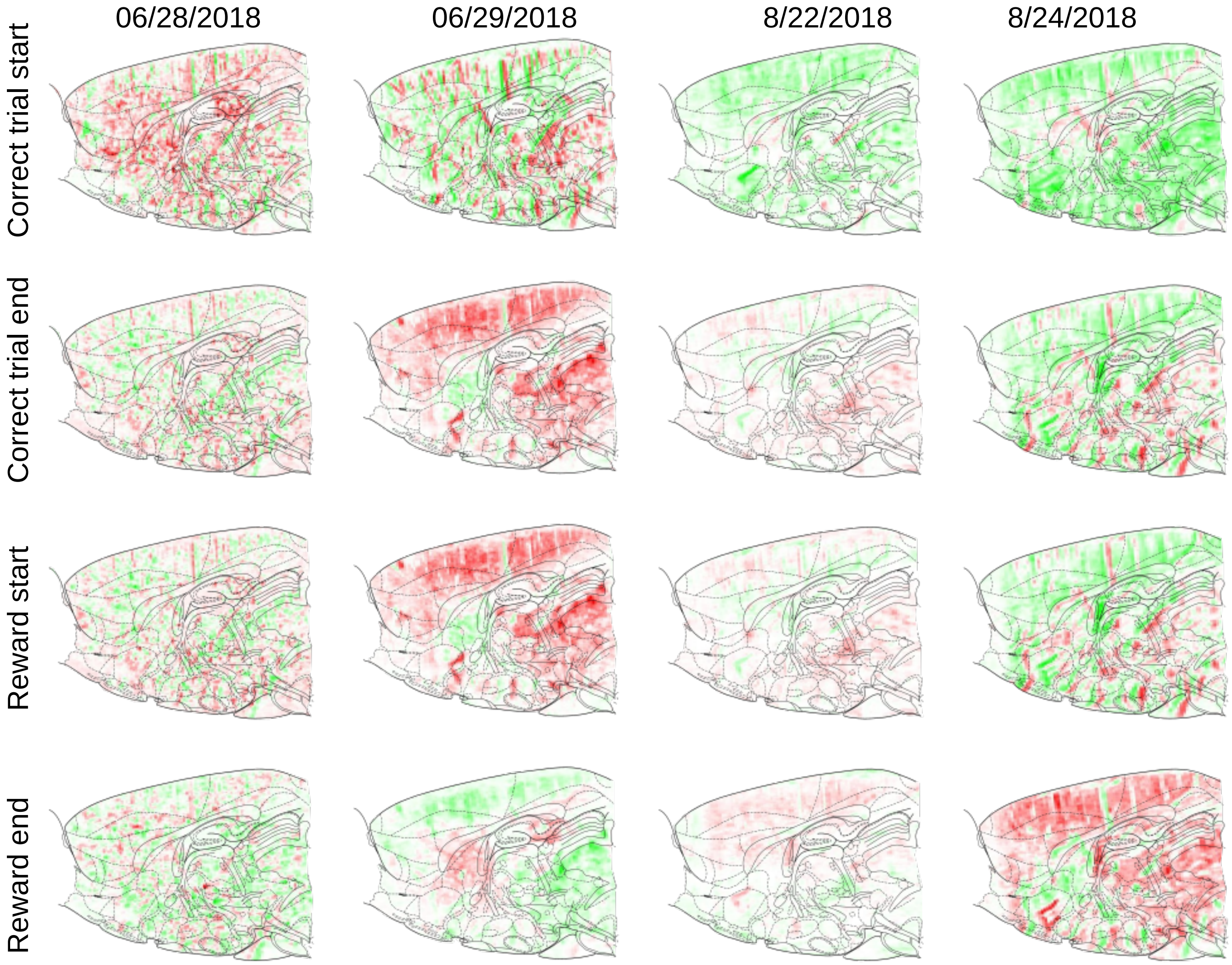


Supplementary Figure 27: Correlation maps for various behavioral events over four sessions spanning two months (continued from Supp. Fig. 26).

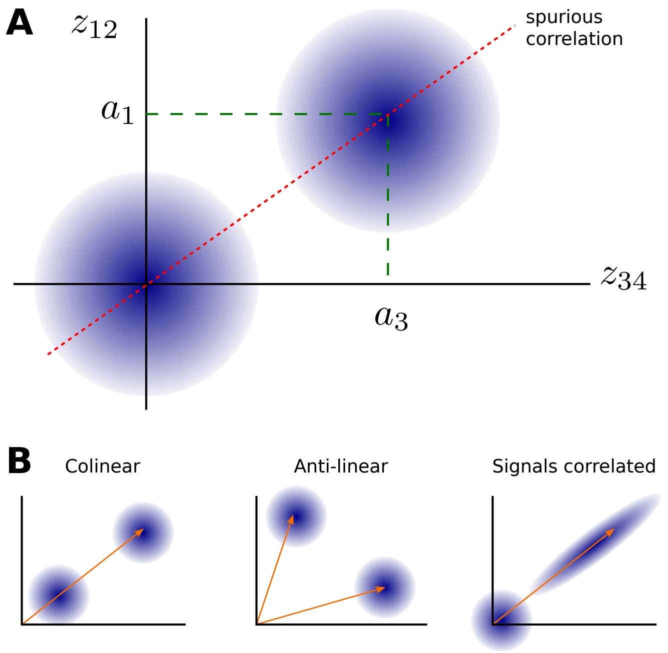


Supplementary Figure 28: Spurious correlations. A: when two vectors $z_{12}$ and $z_{34}$ contains elements from different vectors, interleaved due to motion (or over-correction of motion), differences in amplitudes can create spurious correlation. B: Spurious correlations can be positive (left) due to colinear alignment, they can be negative (middle) due to anti-linear alignment, and they can even influence otherwise correlated vectors (right).
