## Supplemental surgery guide and code documentation for "Chronic brain functional ultrasound imaging in freely moving rodents performing cognitive tasks"

#### Abstract

Functional ultrasound imaging (fUS) is an emerging imaging technique that indirectly measures neural activity via changes in blood volume. To date it has not been used to image chronically during cognitive tasks in freely moving animals. Performing those experiments faces a number of exceptional challenges: performing large durable craniotomies with chronic implants, designing behavioural experiments matching the hemodynamic timescale, stabilizing the ultrasound probe during freely moving behavior, accurately assessing motion artifacts and validating that the animal can perform cognitive tasks at high performance while tethered. In this study, we provide validated solutions for those technical challenges. In addition, we present standardized step-by-step reproducible protocols, procedures and data processing pipelines that open up the opportunity to perform fUS in freely moving rodents performing complex cognitive tasks. Moreover, we present proof-of-concept analysis of brain dynamics during a decision making task. We obtain stable recordings from which we can robustly decode task variables from fUS data over multiple months. Moreover, we find that brain wide imaging through hemodynamic response is nonlinearly related to cognitive variables, such as task difficulty, as compared to sensory responses previously explored.

### A Surgical Procedure

This appendix encompasses all details related to the performance of the surgery aimed at creating a large craniotomy on which a headplate can be fixed and functional ultrasoung imaging recording performed from same imaging plane over many months. The appendix includes three sections: first the surgical procedure explained in full details in text, second a step by step pictorial representation of the procedure step by step and third a list of the needed supplies and materials.

Surgical procedure:

Surgery day - Preparing the animal for surgery:

1. The animal is brought to the surgery room, then placed in the induction chamber with 4% isoflurane.

2. The animal is administered Dexamethasone 1 mg/kg IM to reduce brain swelling and Buprenorphine 0.01–0.05 mg/kg IP for pain management

3. The animal is put back to the induction chamber (with 4% isoflurane) until the animal achieves the appropriate depth of anesthesia. The depth of anesthesia can be tested by pinching the hindpaw. If the hindpaw reflex does not happen, this indicates that the animal has reached the appropriate depth of anaesthesia.

4. The rat hair ,covering his head, is shaved using an electric trimmer. It is important to remove all the hair on the head skin that will be the site of incision afterwards. One should also slightly trim the hair of the surrounding areas. Particular attention should be paid to the areas around the ear where there is a lot of growing hair that can infect the incision site if it is left too long.

5. The animal head is put in the ear bar. It is performed by holding gently the skin under the ear and inserting the ear bar slowly into the ear till it stabilizes on the bones inside the ear . This is critical for the rest of surgery, so make sure that the animal is in the ear bar properly by testing the stability of the insertion to the ear bar. The stability can be tested by pushing slightly with your thumb over the side of the rat head (if the rat head falls from the ear bar then this means that the rats head was not properly inserted in the earbar)

6. The concentration of isoflurane is slowly reduced to 2%, ideally below or equal to 2%. The hindpaw is pinched in order to ensure the appropriate depth of anesthesia has been reached.

7. The shaved head is cleaned copiously with alternating betadine and alcohol. This procedure is repeated at least 3 times. One should make sure to clean a wide region around the area where one plans to do the incision.

Surgery day - Surgery

1. From now on , the procedure becomes sterile meaning that there are two people working (one is the surgeon and the other is the helper). From now on, we will call the Surgeon A and the Helper B. Surgeon A uses sterile gloves. B gives the sterile drape to A and opens up all the sterilized surgical instruments to Surgeon A. It is important to note that the instruments are autoclaved beforehand in sterile bags ( ideally one day before the surgery) . It is crucial

that the Surgeon A and helper B both use appropriate techniques to keep the sterility of the instruments. If it is your first time doing this procedure, you should watch some online videos about sterile procedures or watch a sterile surgery done in person. Anything that touches the animal's head has to be sterile (autoclaved or chemically sterilized), in particular the surgeon's hand gloves and tools. During the surgery, the surgeon A should not touch anything that was not sterilized, instead should ask the Helper B for those tasks.

2. Once an appropriate depth of anesthesia is confirmed, the surgical procedure can start.

3. Sterile surgical drapes are put on the surgical stage. 0.15 ml a mixture of lidocaine / norepinephrine (2% lidocaine with norepinephrine 20 $\mu$g/mL mixture under the head skin where the incision is planned. Wait for five minutes) is injected into the head skin while holding the center part of the skin using a serrated Adson forceps.

4. The skull overlying the brain areas of interest is exposed by making an incision ($\sim$10-20 mm, rostral/caudal orientation) along the top of the head, through skin and muscle using a surgical scalpel feather blade (that is put in a scalpel handle before usage). Tissue will be reflected back until the skull is exposed and held in place with curved crile forceps or tissue clips depending on the preference of the surgeon performing the procedure.

5. The surface skull is thoroughly cleaned and scrubbed using a double ended Volkman bone curette and sterile saline. The cleaning and scrubbing continue until the surface of the skull is totally clean and devoid of signs of bleeding, blood vessels or clotting.

6. A piece of the skin on the sides ($\sim$2 mm thickness), parallel to the surface of the skull, is cut using sharp/sharp operating curved scissors. This is important to avoid excess skin that will make more difficult to implant the headplate in place stably.

7. Bregma is marked with a sterile pen then a rectangle is drawn using a sterile waterproof ruler. The center of mass of the rectangle is Bregma and it is 19 mm AP x 8 mm ML. This step will depend on the imaging planes including which areas the experimenters are planning to image.

8. Using the surgical scalpel blades, diagonal lines are engraved on the surface of the skull.

9. Metabond (C&B Metabond Quick Adhesive Cement System) is then prepared and put on ice by Helper B. Then using a small sterile brush supplied with the C&B Metabond Quick Adhesive Cement System , metabond is put all over the surface of the skull (except the area onto which the rectangle is drawn) by the Surgeon A. It is highly advisable to put low toxicity silicone adhesive KWIK-SIL in the craniotomy region (the drawn rectangle delineating the skull area that will be removed in subsquent steps) using gas sterilized KWIK-SIL mixing tips to avoid that any metabond covers this area. In this way one avoid the extra work to drill through the metabond and also diminish the chance that one loose track from the desired craniotomy borders.

10. Using a positioning device (shown in Fig. 9), the sterile headplate is positioned over the drawn rectangle on the surface of the skull. The headplate is pushed down pressing on the skin but taking care to allow for enough slack for the skin muscles underneath.

11. Metabond is prepared and put on ice by Helper B. Then it is put by Surgeon A, using a sterile brush, on the area surrounding the headplate beginning from the outer part that is

covered with metabond. Moving to the inner part of the head plate, use the minimal amount of metabound possible down to one layer taking care not to spill over the drawn rectangle on the skull. It is important that the inner part of the head plate is properly sealed to avoid any leaking. If there is any leaking, it can be a source of infection or instability of the headplate.

12. The positioning system to which the headplate is attached and pushing on the skull is left in place for ∼15 minutes in order to allow for the metabond to dry and glue the headplate in place to the skull.

13. The headplate is then slowly unscrewed from the positioning system and the positioning system is then slowly raised leaving the headplate firmly adhered to the skull. It is very crucial at this point to test two things: whether the headplate is stable and whether the skin underneath the sides of the headplate is not totally immobilized. A very slight slack will allow the skin and muscles not to develop necrosis.

14. The skull is then copiously rinsed with sterile saline.

15. The piezosurgery apparatus is set up by Helper B using the Micro-saw insert and set the speed of the apparatus at slow speed.

16. Surgeon A then changes his sterile gloves to new sterile gloves to begin doing the craniotomy.

17. Surgeon A slowly use the piezosurgery drilling to remove the skull demarcated by the rectangle drawn by the sterile pen beforehand . Helper B will be continuously applying sterile saline to the area ensuring that it is always wet in order to prevent overheating from the piezosurgery drilling. It is very important to keep the surface of the skull wet while drilling to avoid overheating and skull / tissue damage. The drilling using piezosurgery is performed one side a time. It is recommended that the craniotomy is performed in the following order: first drilling into the skull in the upper and lower parts then continue by drilling through the sides. The whole craniotomy procedure should be done very slowly and the time it takes depends on the size of it. The advantage of using the piezosurgery drilling is that it cuts only through the bone and not through the vessels thus avoiding bleeding nevertheless one should remain vigilant during the craniotomy.

18. In case of older rats, bleeding from the skull can happen excessively so surgeon A should make sure to use Sugi Eyespear (pointed tip with handle ) to dry it to be able to perform the craniotomy smoothly.

19. Once Surgeon A feels that the skull delineated by the sterile pen is not attached the dura anymore, the skull is removed very slowly under continuous flushing using sterile saline with the support of helper B . The surgeon A uses a dissecting knife to clear the attachment of the dura to the skull, while helper B is holding the opposing side (to where the surgeon is clearing) using a serrated curved Graefe forceps . This step has to be done very slowly coordinating the movement of both the surgeon and the helper. If done properly, you should be able to have a large craniotomy with perfectly intact dura.

20. Once the piece of skull is removed, sterile absorbable gelatin sponge is cut into a piece that fits into the craniotomy then soaked with sterile saline and placed on the brain in order to drain any bleeding.

21. The autoclaved metal headcover is then screwed to the headplate then a layer of KWIK-SIL is added on the sides of the headplate.

22. After that, as the portion of incised skin is usually bigger than the headplate. The remaining areas around the headplate (between the headplate and the incised skin) are filled with metabond then the whole skin is closed with a very small drops of VETBOND.

23. The surgeon A should make sure that metabond is hardened then the rat is removed from the anesthesia.

24. The rat is left to wake up in the cage with a heat pad under the cage.

25. After 8-12 hours (calculated from the beginning of the surgery), 0.05 mg/kg buprenorphine and 1 mg / kg dexamethasone are administered.

26. After 24 hours , 0.05 mg/kg buprenorphine and 1 mg/kg dexamethasone are administered.

27. After 36 hours, 0.025 mg/kg buprenorphine and 0.5 mg /kg dexamethasone are administered.

28. After 48 hours, 0.025 mg/kg buprenorphine and 0.5 mg/kg dexamethasone is administered.

29. After 72 hours, 0.25 mg /kg dexamethasone is administered.

30. The aformentioned drug regimen insures that rats do not develop excessive inflammation or pain as a result of the surgical procedure.

31. The rat is then left on free water for 10-14 days which is considered the recovery period before any experiments are done on him.

    After recovery, the following steps are performed to insert the mock probe and streamline the functional ultrasound imaging so that we do not need to anaesthetize the rat every time we are performing an imaging session.

    (a) After the recovery from the surgery, the rat is ready for experiments.
    (b) In order to perform the functional ultrasound imaging, there should be a direct access to the open cranial window (that we opened via craniotomy during the surgery)
    (c) First the rat is anaesthetized . Throughout the whole coming procedure, sterile procedures are to be strictly followed in order to prevent infection of the exposed brain surface.
    (d) The rat is then put in ear bars centered
    (e) The headcover is unscrewed and removed.
    (f) The brain surface is flushed with sterile saline then a piece of Polymethelyene Pentene film (PMP) ,cut to a size that fits the craniotomy ,is glued onto the surface of the brain by carefully applying small drops of VETBOND around the sides. It is crucial to make sure that the PMP is tighly glued to the surface of the brain. This step is very crucial to ensure the total minimization of the brain motion during functional ultrasound imaging in a freely moving setting.
    (g) Afterwards the probeholder is screwed onto the headplate. Note that in this case the headcover is not screwed back. Ideally when you look through the probe holder , one can see the brain surface onto which the PMP is glued .

(h) The 3D printed mock probe is added for further protection while the rat is in his cage.

(i) To perform the ultrasound imaging, the rat is taken to the experimental chamber and put inside the operant behavior box.

(j) the mock probe is removed from inside the probe holder.

(k) De-bubbled sterile ultrasonic gel is added on the surface of fUSi probe.

(l) Gently hold the rat and insert the fUS probe ( covered with ultrasonic gel) inside the probe holder.

In order to ensure that the fUS can be performed chronically over extended periods of time, one should perform regular checks and cleaning of the craniotomy every two - three weeks. It is performed as follows:

(a) First the rat is anaesthesized . Throughout the whole coming procedure, sterile procedures are to be strictly followed in order to prevent infection of the exposed brain surface.

(b) The rat is then put in ear bars centered

(c) The headplate along with the probe holder are unscrewed and removed.

(d) The PMP (glued with VETBOND to the surface of the brain ) is slowly removed using a forceps by holding it from the side and slowly taking it off the brain surface.

(e) The brain surface is then flushed with alternating chlorhexidine 0.05% and sterile water. This alternation is repeated for three times.

(f) New PMP piece matching the size of the craniotomy is added and glued onto the surface of the brain by carefully applying small drops of vetbond around the sides.

.

| Surgical Material / Instrument | Supplier | Reference |
| --- | --- | --- |
| Surgical Scalpel Feather Blades for Handle #4 | World Surgical Instruments | https://www.wpiinc.com/var-504169-feather-blades |
| Scalpel Handle #3 | World Surgical Instruments | https://www.wpiinc.com/var-500236-scalpel-handles |
| Dumont Tweezers #5, 0.1x0.06 mm, Dumoxel | World Surgical Instruments | https://www.wpiinc.com/14098-dumont-tweezers-5-01-x-006mm-dumoxel |
| Adson Forceps, 12 cm, Serrated Tips | World Surgical Instruments | https://www.wpiinc.com/var-14226-adson-forceps-12cm |
| Curved Graefe Forceps, 7 cm, Serrated | World Surgical Instruments | https://www.wpiinc.com/var-14142-graefe-forceps-7cm-serrated |
| Curved Crile Forceps | World Surgical Instruments | https://www.wpiinc.com/var-501242-crile-forceps |
| Volkman Bone Curette, 17 cm, Double Ended | World Surgical Instruments | https://www.wpiinc.com/503752-volkman-bone-curette-17cm-double-ended |
| Sharp/Sharp Operating Scissors, Curved, 14 cm | World Surgical Instruments | https://www.wpiinc.com/var-501220-operating-scissors-curved-14cm |
| Dissecting Knife, 13 cm, 1 mm Blade, Right Angle | World Surgical Instruments | https://www.wpiinc.com/14135-dissecting-knife-13cm-1mm-blade-right-angle |
| Sterile - 10 ct - Ultra fine Value Tip Marker and fluid resistant waterproof ruler | Viscot | https://www.viscot.com/products/value-ultra-fine-marker/ |
| C&B Metabond® Quick Adhesive Cement System | Parkell | https://www.parkell.com/C-B-Metabond-Quick-Adhesive-Cement-System |
| Low Toxicity Silicone Adhesive (KWIK-SIL) | World Surgical Instruments | https://www.wpiinc.com/kwik-sil-low-toxicity-silicone-adhesive |
| Replacement Mixing Tips for KWIK-SIL | World Surgical Instruments | https://www.wpiinc.com/600022-replacement-mixing-tips-for-kwik-sil-and-kwik-cast-pkg-of-10 |
| Piezosurgery Apparatus | Piezosurgery Inc. | https://dental.mectron.us/en/products/piezosurgeryr/units/piezosurgeryr-white/ |
| Pointed insert | Piezosurgery Inc. | https://dental.mectron.us/en/products/piezosurgeryr/inserts-implant-prep/im1s/ |
| Micro-saw insert | Piezosurgery Inc. | https://dental.mectron.us/en/products/piezosurgeryr/inserts-osteotomy/ot7/ |

| | | |
|---|---|---|
| Sugi® Eyespear (pointed tip), rectangular white handle, sterile | Questalpha GmbH | https://www.questalpha.com/sugi-products/details/product/eyespear |
| Gelfoam Absorbable Gelatin Sponge - Size 12 | Mwdental | https://www.mwdental.com/supplies/surgical-products/medicaments/gelfoam-absorbable-gelatin-sponge-size-12.html |
| Polymethelyene Pentene film size 150x150 mm and thickness 0.25 mm | Goodfellow Japan | https://www.goodfellow-japan.jp/en/material/polymethylpentene-ME31.htm |
| Parker Laboratories Sterile Aquasonic 100 Ultrasound Gel | Parker Labs | https://www.parkerlabs.com/sterile-aquasonic.asp |
| Butyl Cyanoacrylate, Low Toxicity Adhesive (VETBOND) | World Surgical Instruments | https://www.wpiinc.com/vetbond-butyl-cyanoacrylate-low-toxic |

Table 1: List of materials for fUSi surgery.

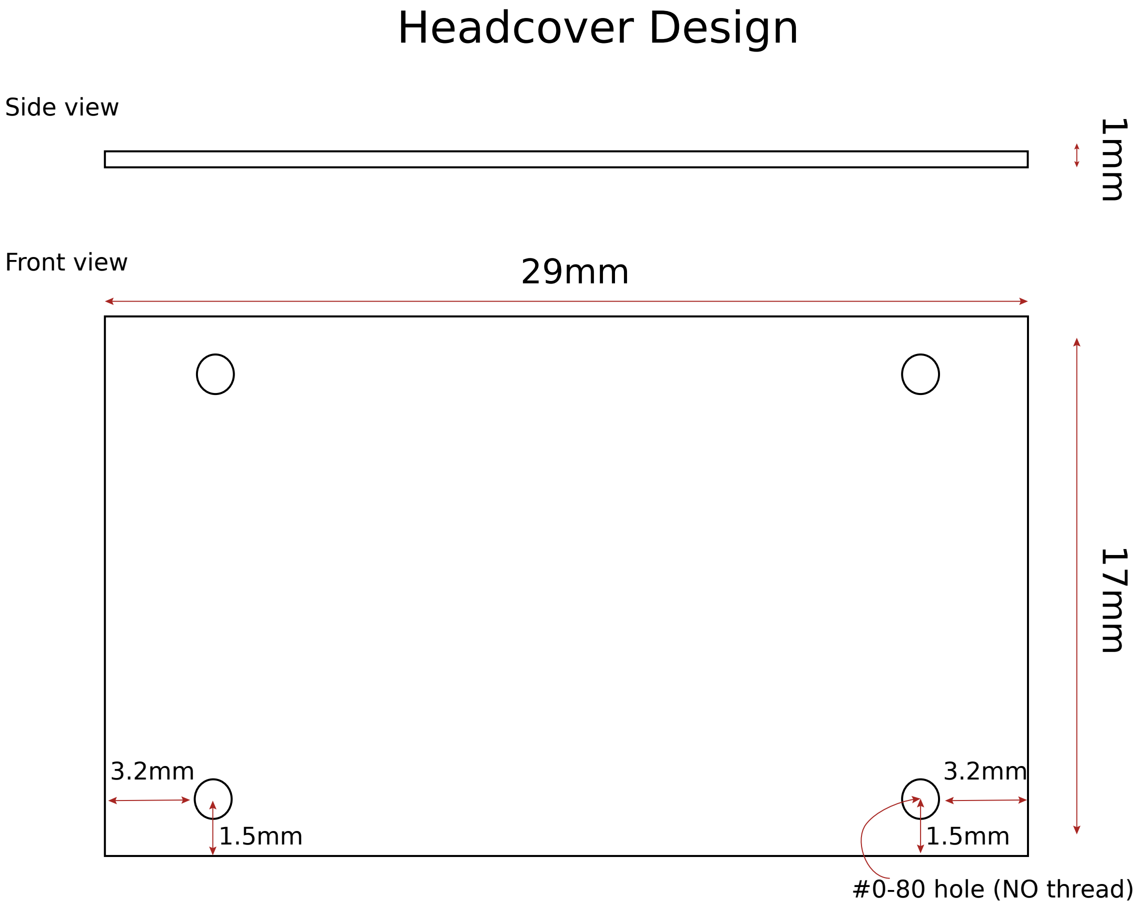

Figure 1: Headcover design

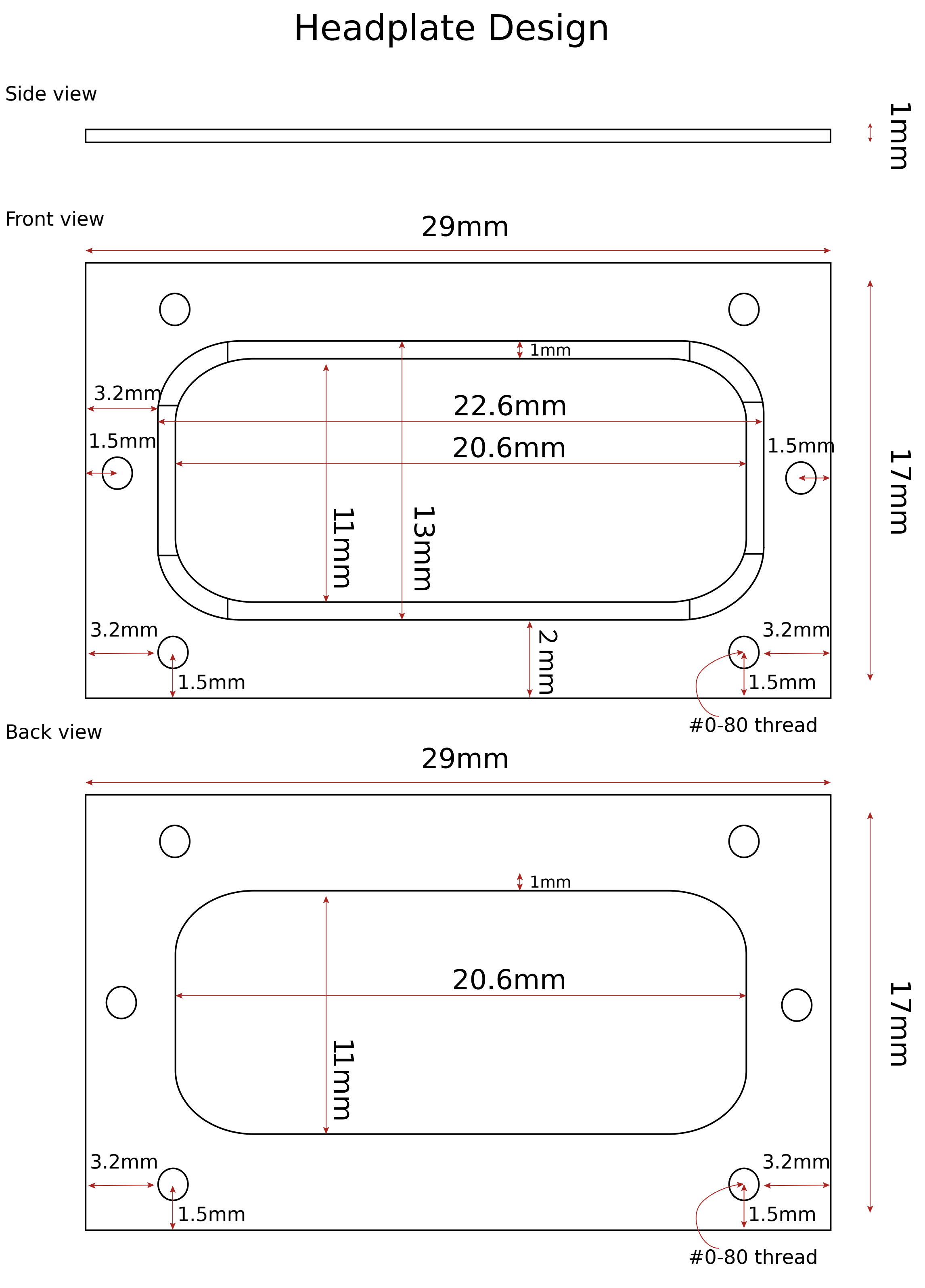

Figure 2: Headplate Design

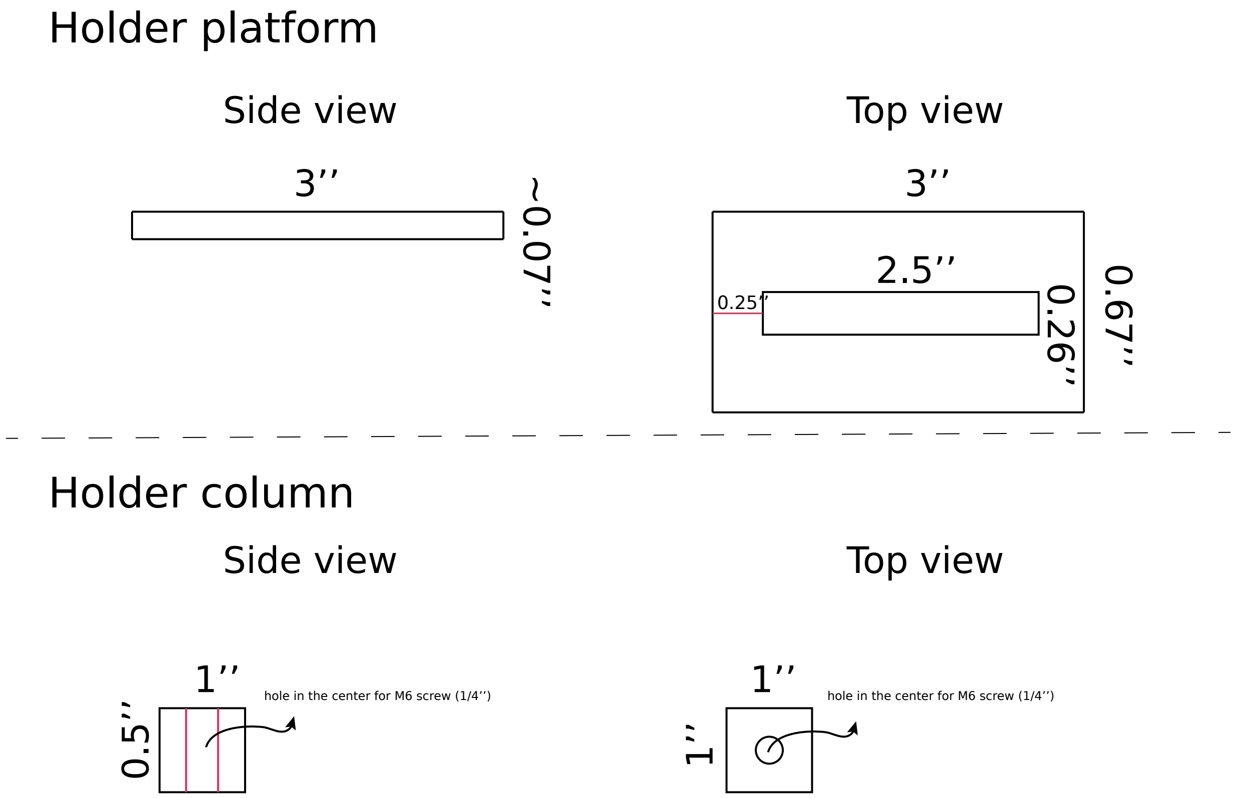

Figure 3: Headplate holder design

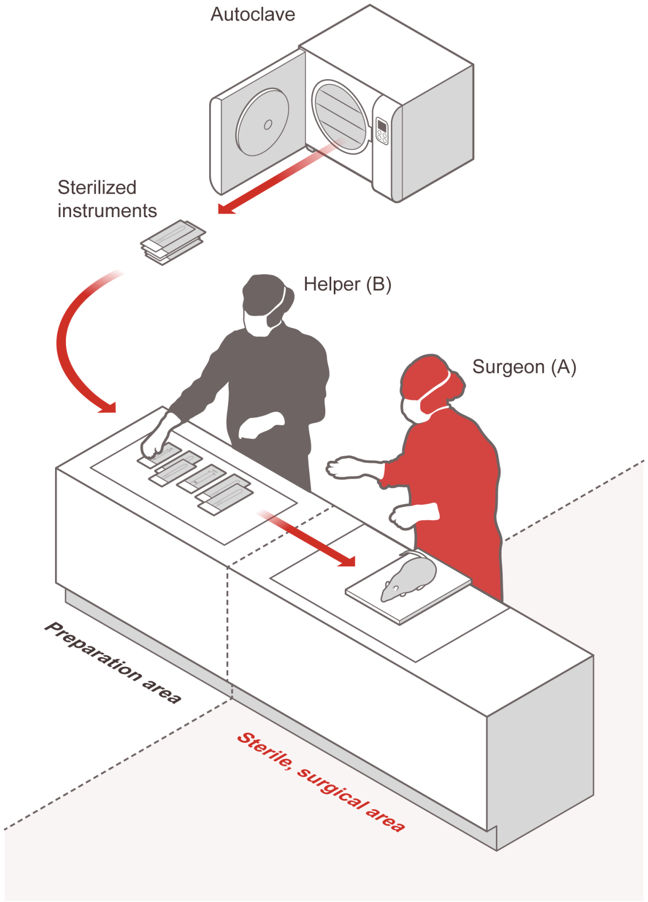

Figure 4: Surgery preparation. All tools used in the surgery should be sterilized and stored in sterile pouches. There are two areas: the preparation area where the helper B puts all the sterilized tools, the sterile surgical area where the surgeon A wearing sterile gloves take the sterile tools from the helper. This configuration ensures that the surgery is following a strict sterile procedure

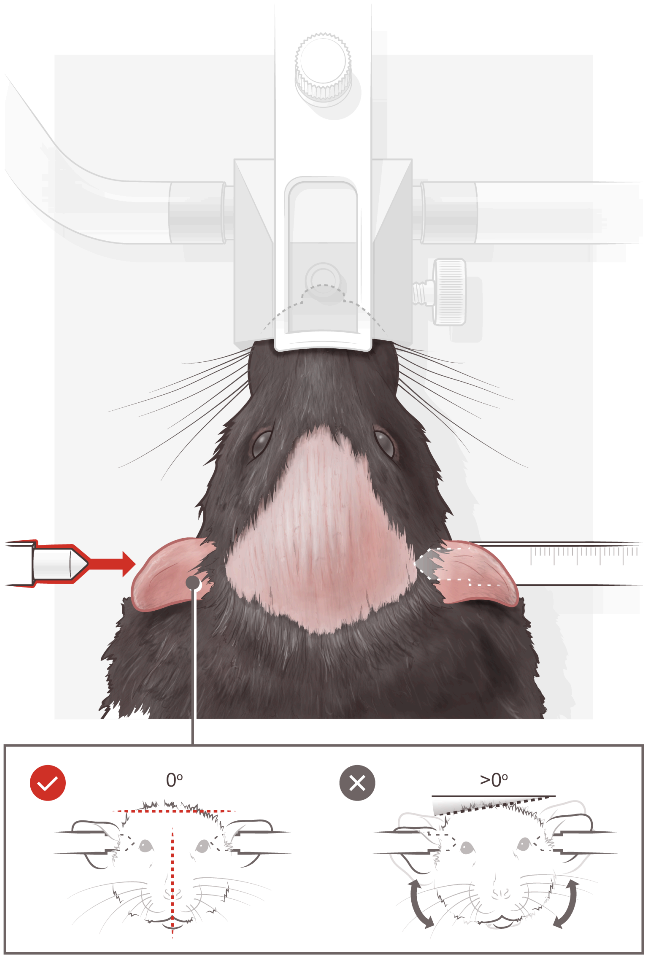

Figure 5: Ear bar insertion to stabilize the rat head. Note that the rat head is exposed after shaving ( the dark rose part in the middle). The shaving should be wide enough close to the ears and to the frontal side of the head. The inset shows the correct orientation of the head when inserted in the earbar

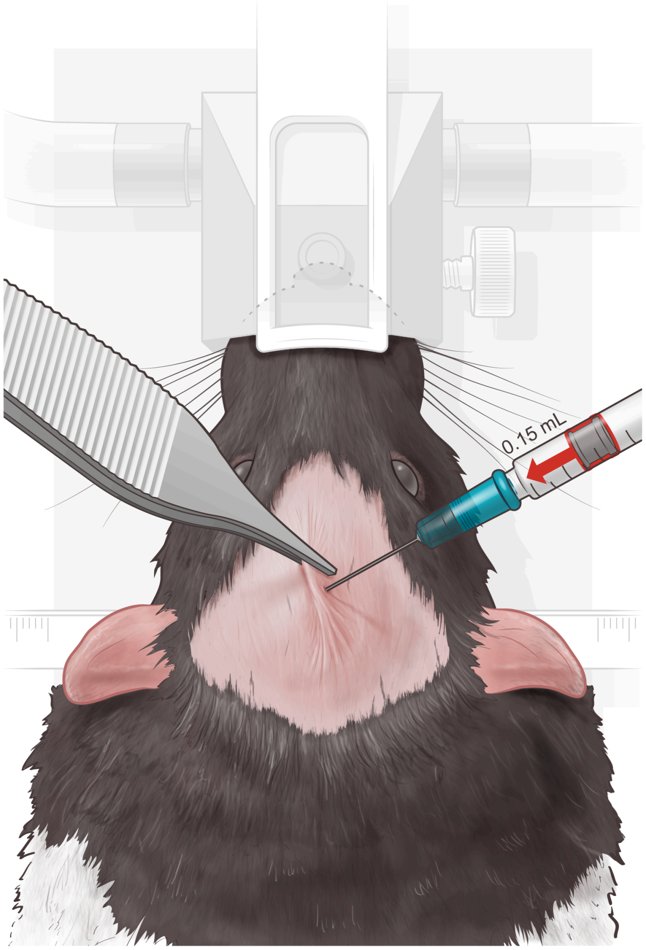

Figure 6: Lidocaine/Norepinephrine injection. Use a forceps to hold the skin to inject 0.15 ml lidocaine/norepinephrine mixture into the skin.

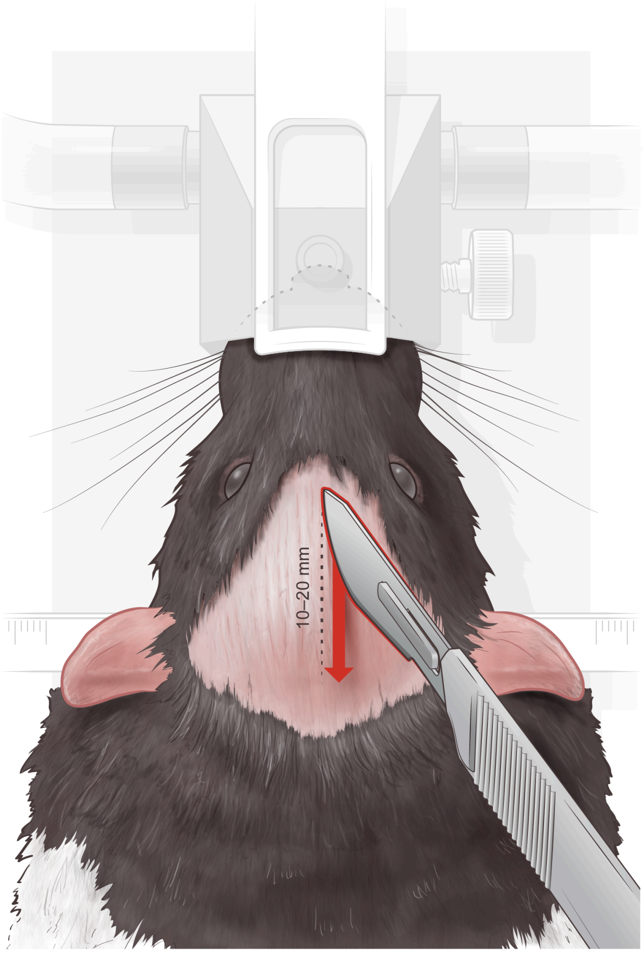

Figure 7: Use the scalpel to do a lateral incision in the skin varying between 10 - 20 mm depending on the intended size of the craniotomy to be performed afterwards

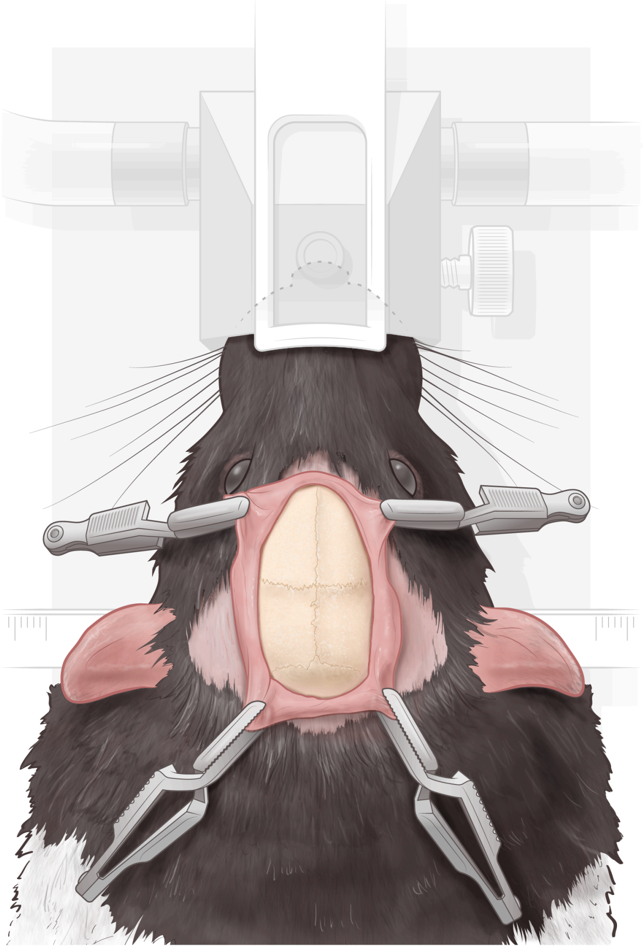

Figure 8: Using four surgical clips ,attached to the periosytum, to expose the skull and prevent the skin from folding over the skull. The skull appears clean after scrubbing away any remaining tissue debris or blood

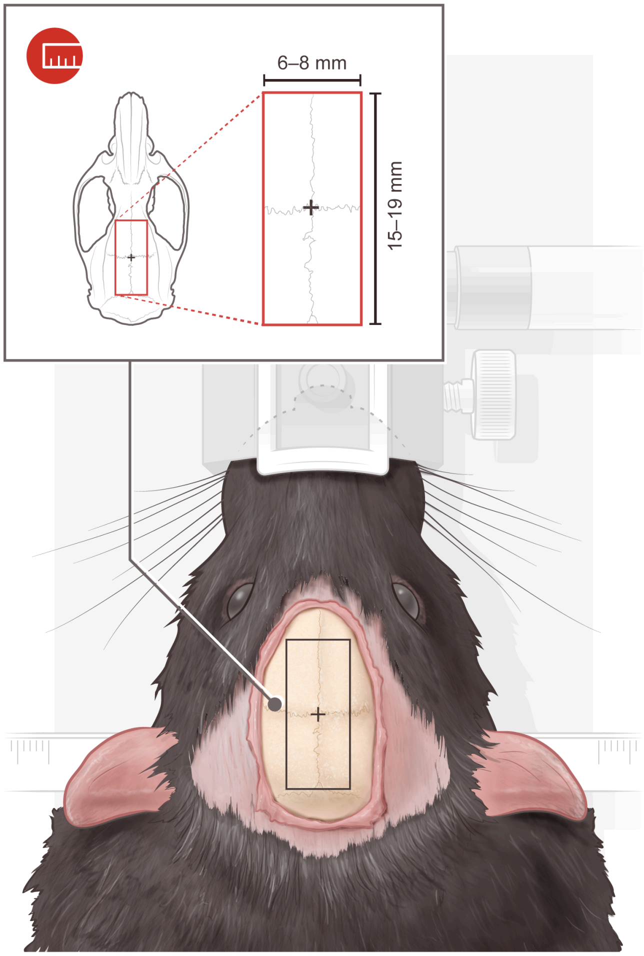

Figure 9: The surgical clips should remain there constantly but we remove them for clarity of presentation. The black rectangle represents the contours of the skull area in which the craniotomy will be performed. The black rectangle is marked and drawn using a sterile pen and a sterile water-proof ruler. The size of the rectangle can vary between 6 - 8 mm and 15 - 19 mm depending on the desired size of the craniotomy

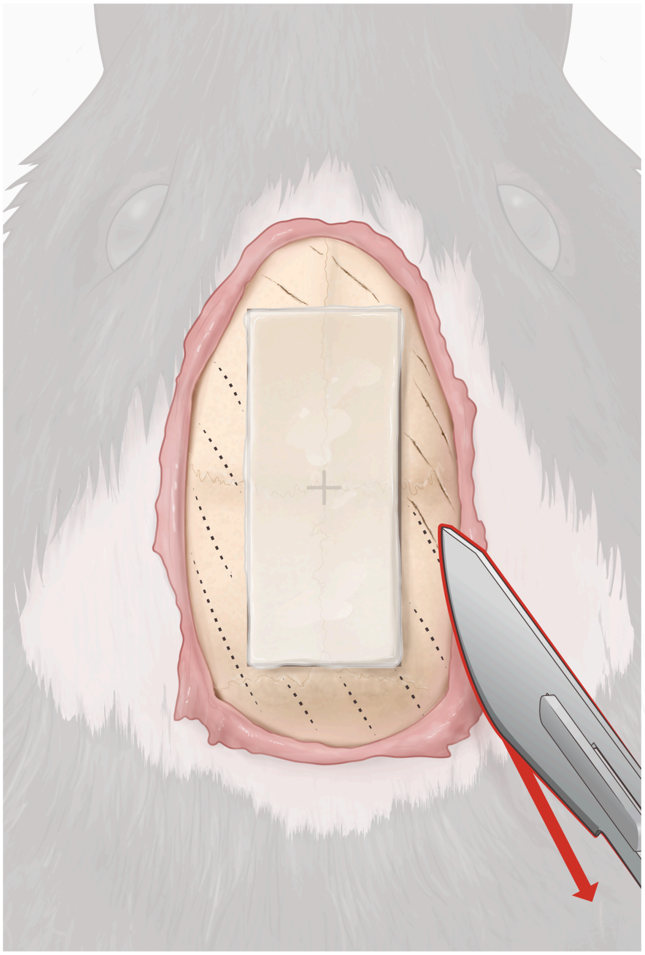

Figure 10: Kwik-sil is put on the rectangle carefully to fill in the black drawn rectangle then extras are cut using a scalpel to make sure that the rectangular gel piece is homogeneous and fits on the black rectangle marked on the skull. Using the scalpel one makes marks on the surface of the skull around the rectangular gel

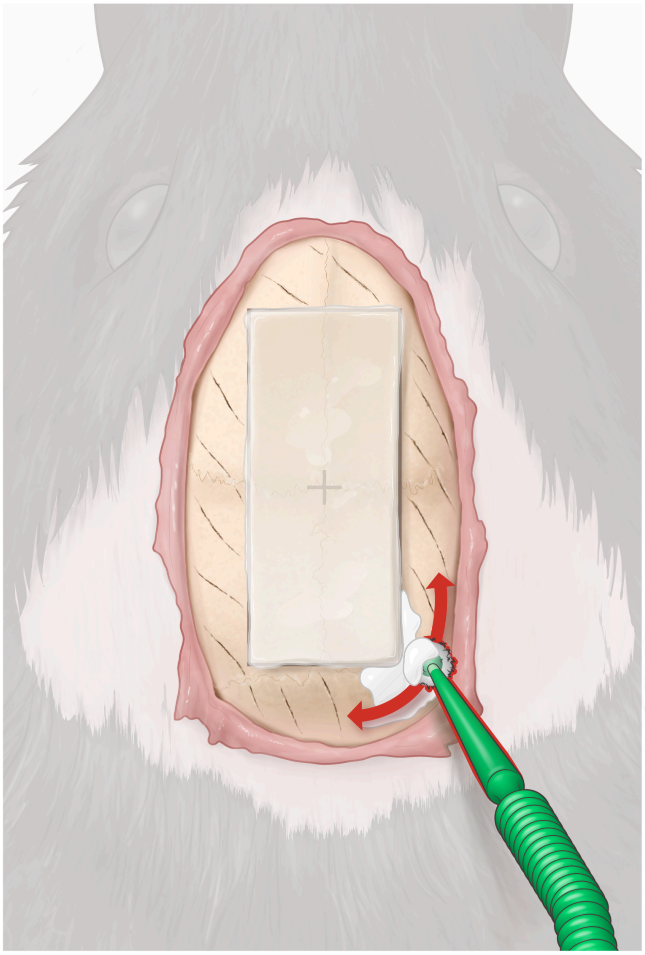

Figure 11: Using a sterile applicator, one puts copious amounts of metabond covering the skull surface around the rectangular gel piece

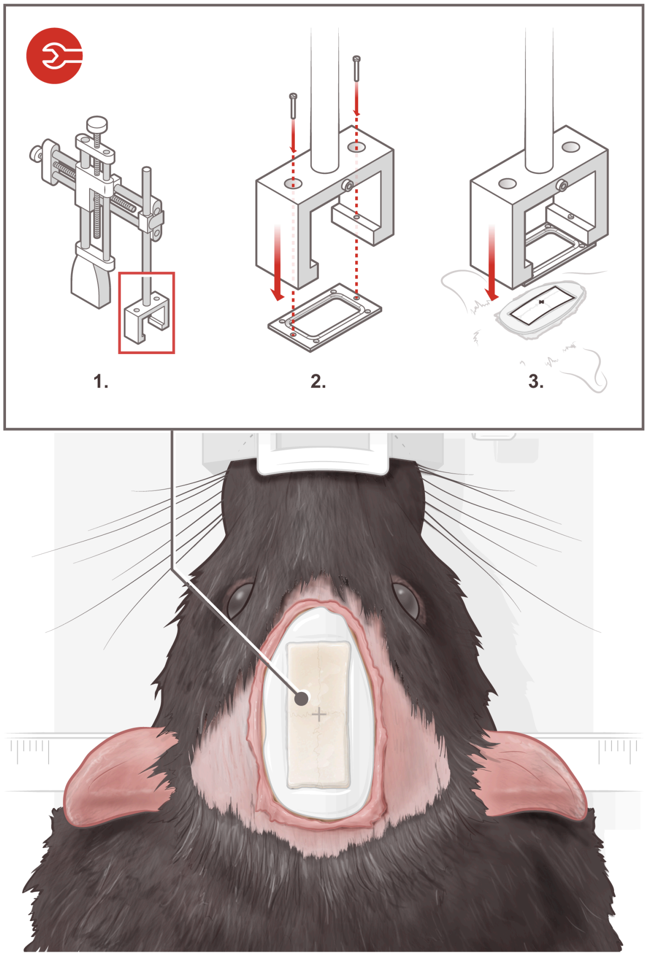

Figure 12: Inset: the headplate is screwed to the headplate positioning device attached to the stereotaxic positioning arm for precise manipulation. Subsequently it is lowered and centered carefully and precisely over the area where the rectangular gel is

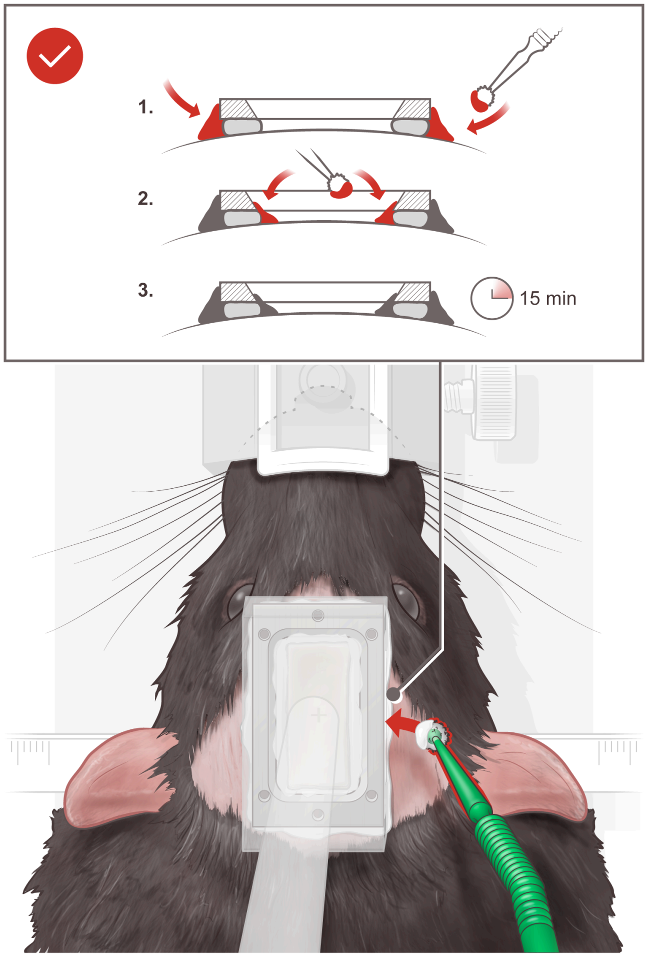

Figure 13: Using a sterile applicator metabond is applied. Metabond is applied on all sides and left for 15 minute. Then the rectangular gel over the skull is removed.

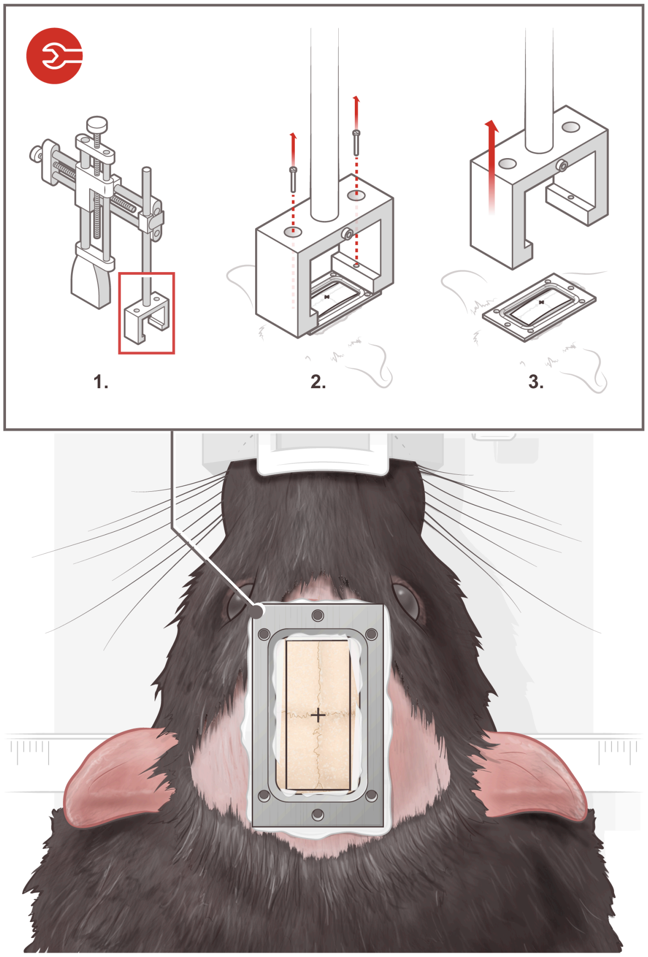

Figure 14: The headplate is then unscrewed from the headplate positioning system and lifted leaving the headplate firmly attached to the skull

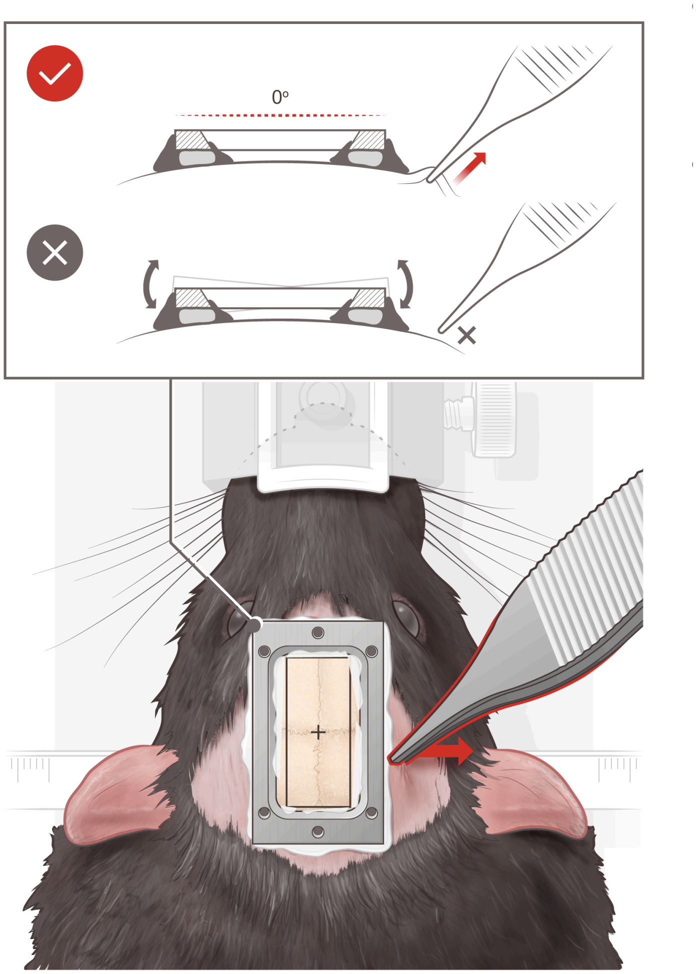

Figure 15: By pulling the skin using forceps, the headplate should be stable in a horizontal position and not move at all

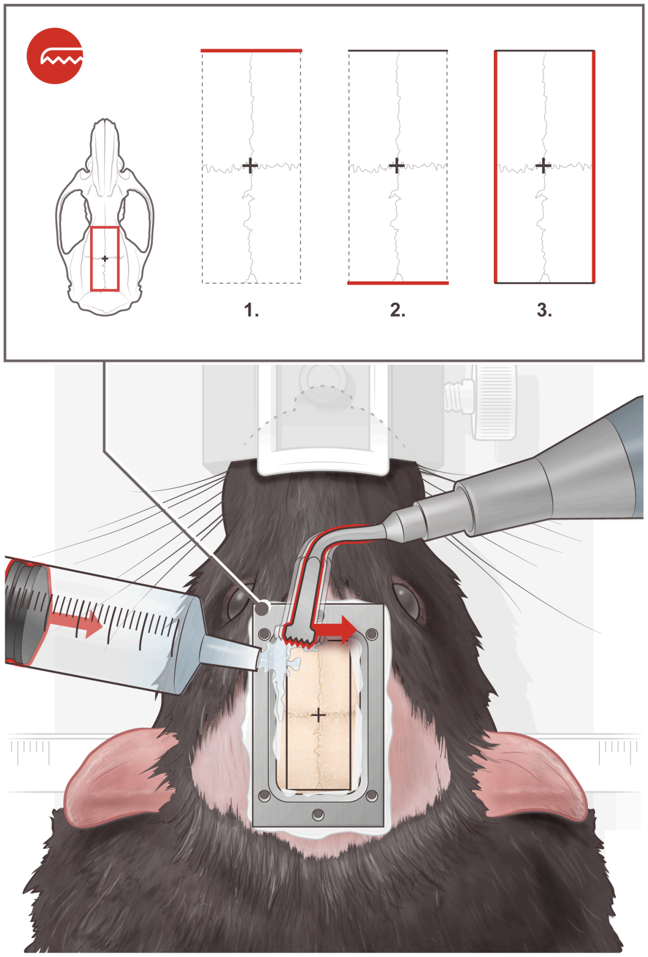

Figure 16: The craniotomy is performed using a piezo-electrical device called piezosurgery apparatus . The craniotomy is performed by beginning to cut , using a micro-saw insert, the upper part followed by lower part then the two sides of the cranial skull surface. To prevent overheating of the skull, sterile water is applied on the skull during the performance of the craniotomy

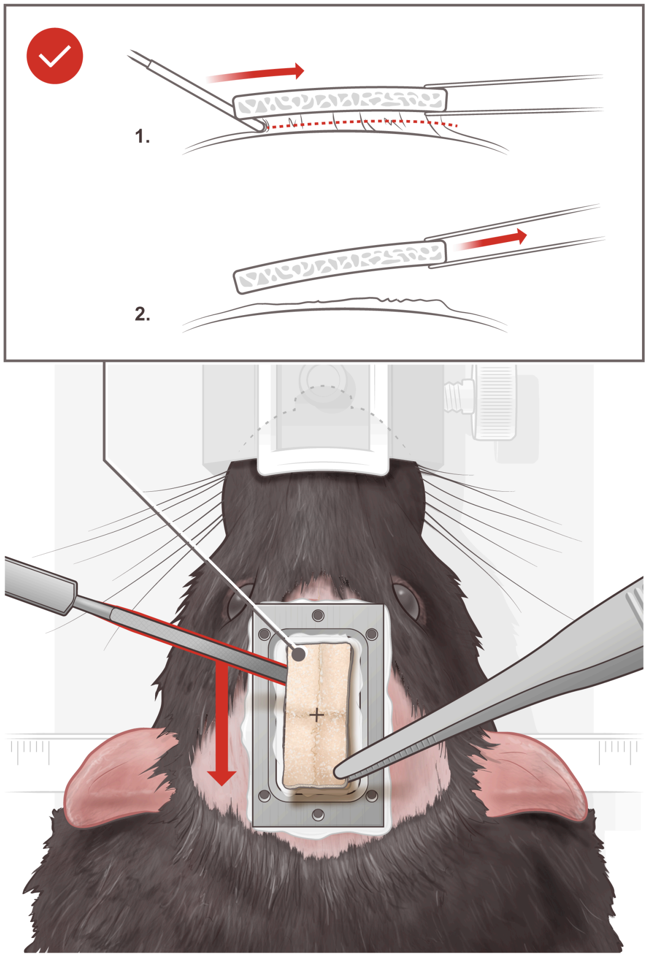

Figure 17: Using a forceps and a dissecting knife the cut skull after craniotomy is carefully removed very slowly.

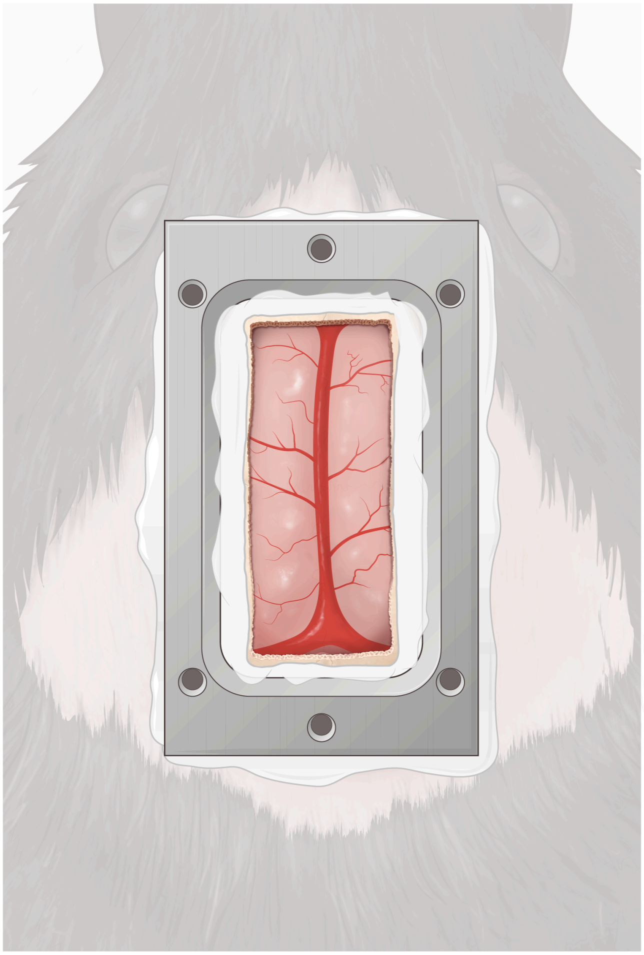

Figure 18: Exposed brain surface with the mid-saggital vein clear along the midline after the removal the skull piece

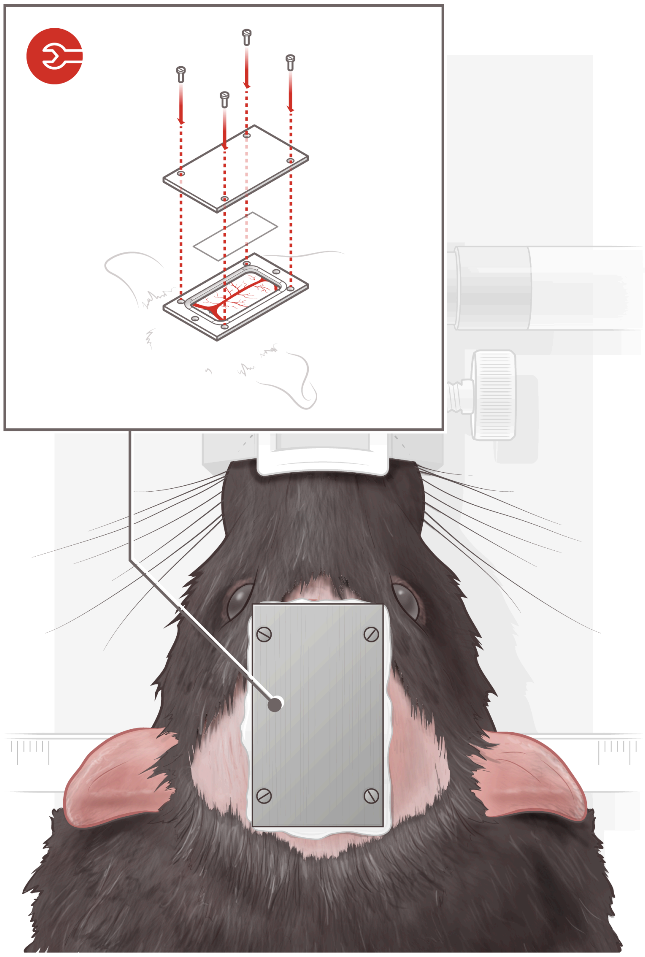

Figure 19: Polymer film (PMP) is put on the brain surface then headcover is screwed to the headplate .

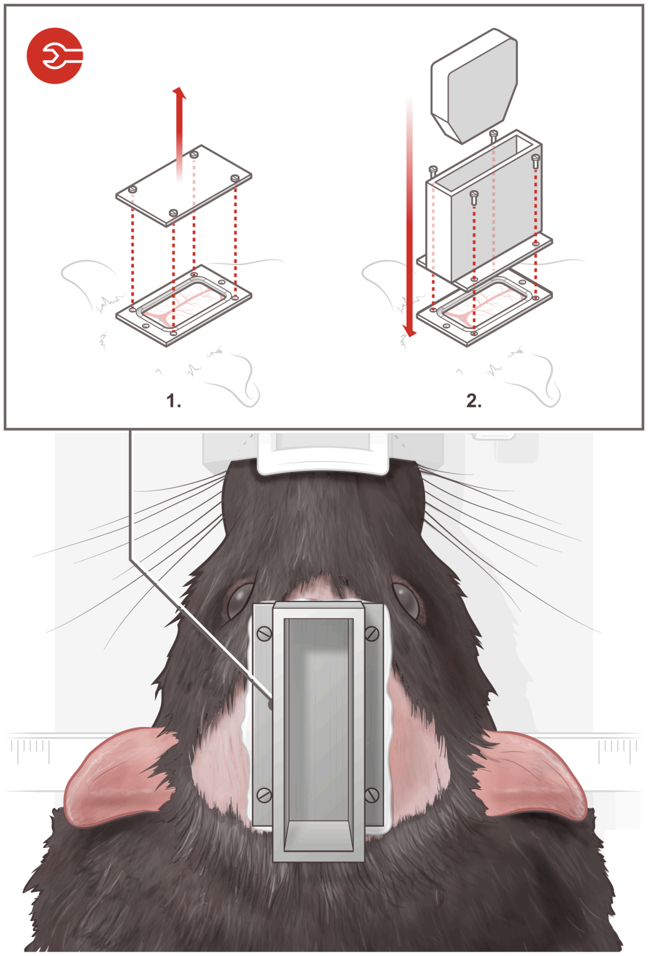

Figure 20: Headcover removed, probeholder attached to the headplate and the mock probe inserted

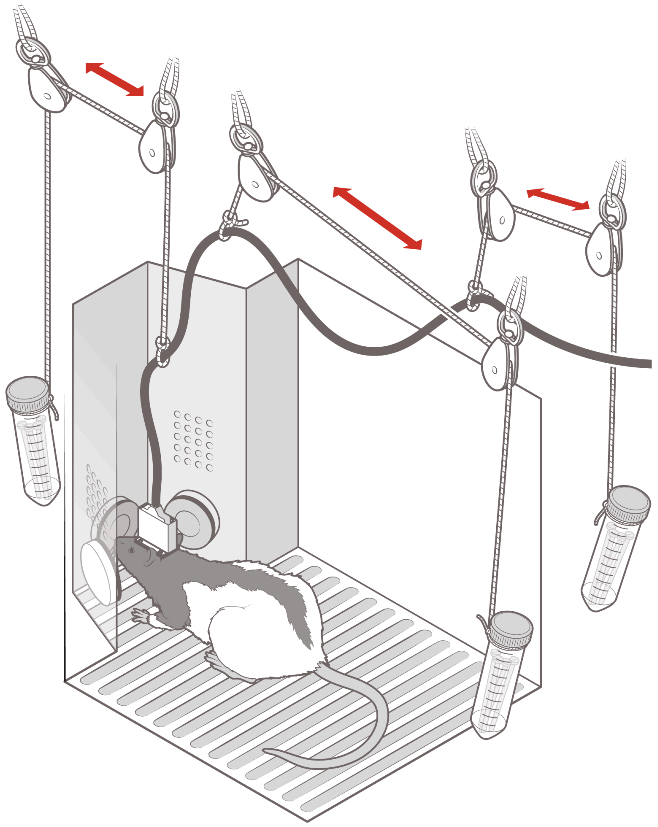

Figure 21: Rat tethered with the functional ultrasound probe inside the behavior operant chamber

### B Code documentation

This documentation outlines the basic functionality and methods implemented in the fUS analysis code-base. The code is written as a pair MATLAB object classes: one for single datasets, and one for analyzing groups of datasets. For a detailed example on the code's use, please see the `demo.m` script in the `code` folder.

#### B.1 fUS and fU-multi object classes

Two classes are defined in this code package: one for the exploration and processing of a single fUS movie, and one for multiple fUS movies taken over a sequence of experiments. The two classes (`uo` and `muo` for ultrasound object and multi-ultrasound object, respectively) provide a concise way of loading, viewing, and processing fUS data. The single data-set object `uo` contains most of the vital class methods and the multi-data set object class `muo` manages a set of `uo` object and handles the overhead of working with many datasets at once.

In initializing a single data-set `uo`, the data may either be loaded and then passed into the object,

```
>>fuo = uo(dataMatrix)
```

Alternatively a path to a `.mat` file can be provided, allowing the code to load data only as necessary:

```
>>fuo = uo([data/path/filename.mat])
```

This latter method of initializing an object is especially important when working with multiple data-sets, as it lowers the memory requirements tremendously. Loading the data directly or setting pointers to the data to prevent the full data being loaded can be toggled by the 'loadToRAM' option as

```
>>fuo = uo([data/path/filename.mat], 'loadToRAM', true)
```

For more details on functions for data loading, see Table 2. To load multiple functional ultrasound datasets into one object, the uom object can be used as

```
>>fuom = uom([data/path/])
```

where the path containing multiple `.mat` files are located and can be identified and accessed. In the creation of the `fuom` object, each identified movie file in the path has a single `uo` object created in `fuom.uo`, along with reasonable meta-data. Functions run on fuom often use the `uo` class functions to efficiently run on all files more conviniently.

#### B.2 Data visualization

Built in to the uo and uom objects are methods to visualize parts or all of the data (see Table 3). For example, one can randomly select a number of traces to view (e.g., Figure 1) by selecting a subset of pixels (in this example 10).

```
>>fuo.displayExampleTraces('numTraces', 10)
```

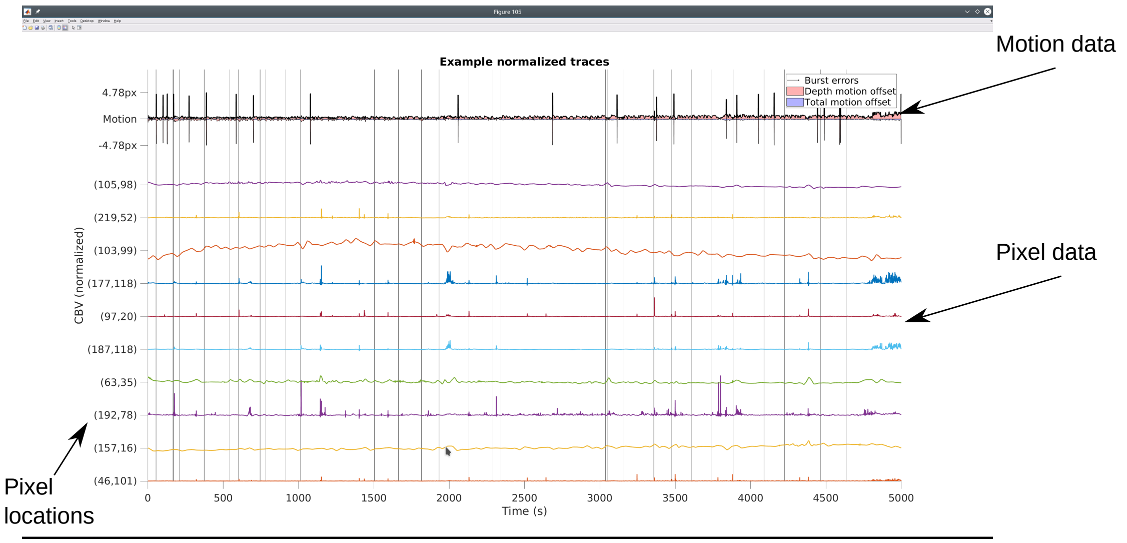

Figure 1: Example of displaying motion and time-trace data from an fUS movie.

To display the entire movie (e.g., Figure 2), the diplayMovie method can be used (NOTE: this method uses the MovieSlider package available at `https://github.com/sakoay/MovieSlider`)

```
>>fuo.displayMovie();
```

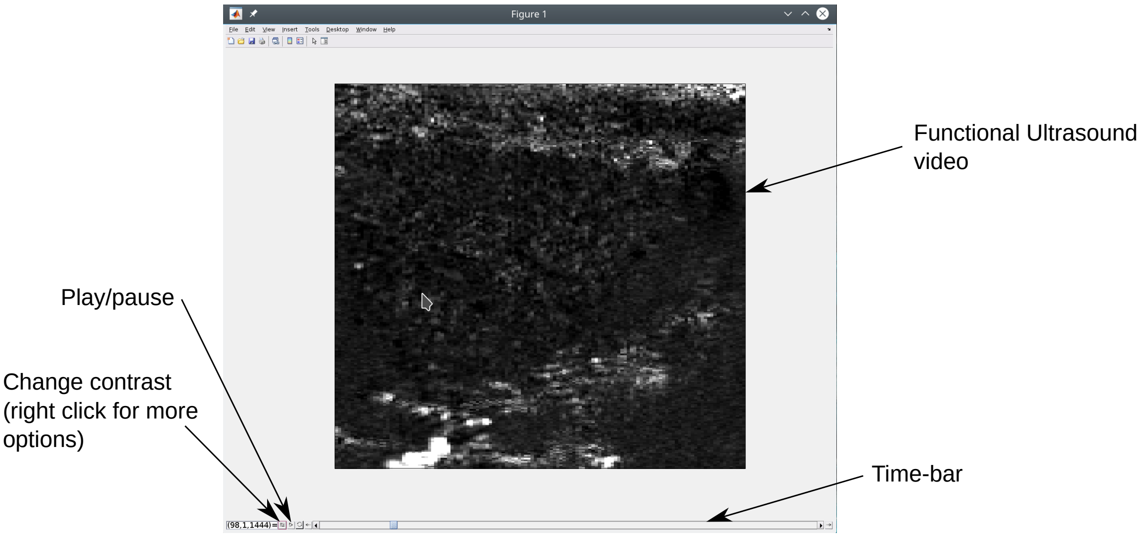

Figure 2: Example of single fUS movie display.

Single frames can also be displayed (e.g., Figure 3) via

```
>>fu.displayExampleFrame('frameNo','rand');
```

#### B.3 Motion and Error Correction

The first step in data analysis of fUS data is to determine where artifacts, in fUS primarily motion artifacts, exist in the data. Knowing which frames should be considered with lower confidence is

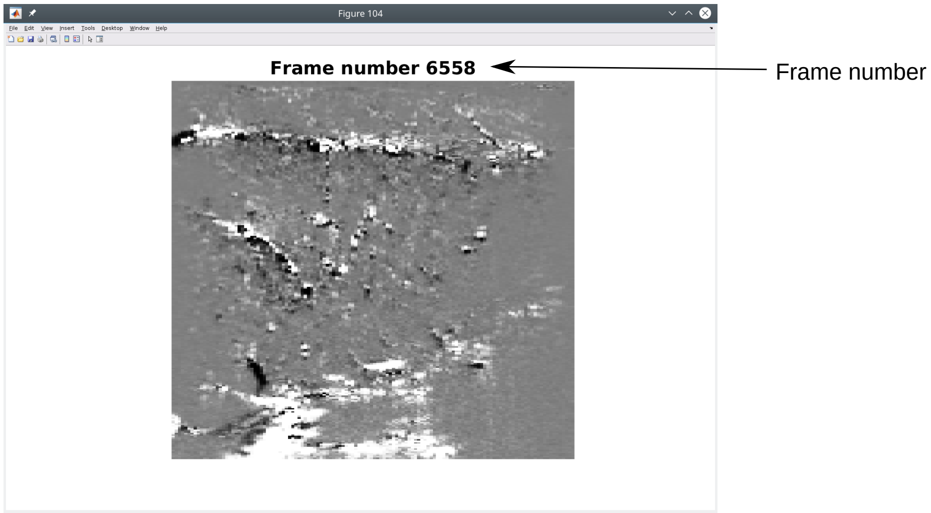

Figure 3: Example of single fUS frame display.

vital to extracting information from the fUS recordings. The single fUS object class `uo` contains methods that isolate both burst errors and lateral rigid motion errors in the data.

Burst errors are defined as sudden, large increases in movie intensity over the entire field-of-view. Such events can be caused, for example, by sudden movements of the animal. Consequently, these events are simple to detect by analyzing the total intensity of the frames, as measured by the $\ell_2$ norm $\sum_{ij} X_{ij}$ for each frame $X$. We dynamically select a cut-off for determining a burst frame be estimating the inflection point in the histogram of frame-norms.

To ascertain the accuracy of the motion correction algorithm, we estimated the residual motion shift using a sub-pixel motion estimation algorithm based on fitting a Laplace function to the autocorrelation $C_{ij} = \langle X_{ref}, X * \delta(x - \tau_i, y - \tau_j) \rangle$, where $X_{ref}$ is the reference image $X$ is the image with the offset we wish to compute. The Laplace function is parameterized as

$$L(i, j; \rho, \mu_x, \mu_y \rho, \sigma_x, \sigma_y) = e^{-|\tau_i - \mu_x|/\sigma_x - |\tau_j - \mu_y|/\sigma_y}, \quad (1)$$

where and $\{\rho, \mu_x, \mu_y, \sigma_x, \sigma_y\}$ is the parameter set for the 2D Laplace function, including the scale, x-shift, y-shift, x-spread and y-spread respectively. We optimize over this parameter space more robustly by operating in the log-data domain, i.e.,

$$\arg \min_{\rho, \mu_x, \mu_y, \sigma_x, \sigma_y} \sum_{ij} \left[\log(C_{ij}) - \log(L(i, j; \rho, \mu_x, \mu_y \rho, \sigma_x, \sigma_y))\right]^2 \quad (2)$$

$$= \arg \min_{\rho, \mu_x, \mu_y, \sigma_x, \sigma_y} \left[\log(C_{ij}) - \log(\rho) - \frac{|\tau_i - \mu_x|}{\sigma_x} - \frac{|\tau_j - \mu_y|}{\sigma_y}\right]^2 \quad (3)$$

which makes the optimization more numerically stable, gradients are easier to compute and solving for $\log(\rho)$ directly removes one of the positivity constraints from the optimization. We further reduce computational time by restricting the correlation range to only shifts of $|\tau_i|, |\tau_j| < 25$ pixels and by initializing the optimization to the max value of the cross-correlation function $\{i, j\} = \arg\max(C_{ij})$.

Computing motion offsets relies on a global reference. If few frames are shifted, than the median image of the movie serves as a reasonable estimate. In some cases this is not possible, and instead other references can be used, e.g., the median image of a batch of frames at start or end of the video. We also provide the option to estimate the motion residual over blocks of frames, sacrificing temporal resolution of the estimate for reduced sensitivity to activity. In fUS, the temporal resolution sacrifice is minimal due to the initial round of motion compensation built into the image rendering.

Burst errors and rigid motion can be corrected via the code

```
>>fuo.findBurstFrames();
>>fuo.findMotionCorrectionError();
```

The first line identifies the burst frames and the second corrects rigid motion. The results of the rigid motion identification and location of burst error frames can be visualized next to the time traces using `fu.displayExampleTraces()`, e.g., see Figure 1. For more details on this functionality, see Table 4. You can also extract the inpainted frames using the command

```
>>[frameInPt,frameIDX] = fuo.computeBurstErrorInpainting();
```

#### B.4 Denoising

The `uo` and `uom` classes enable the application of time-domain denoising via wavelets.

```
>>fuo.denoiseTracesWavelet();
>>fuo.displayMovie('denoised', 'true');
```

`fuo.displayMovie` uses MATLAB's internal wavelet denoising either `wdenoise()` or `wdenoise()` and `cmddenoise()`, which can be selected using the `'wDenoiseFun'` parameter. We suggest using the default `wdenoise()` for speed.

#### B.5 Running GraFT on fUSi data

GraFT can be run using the following command:

```
>>fuo.fuGraFT('n_dict', 45, 'lamForb', 0.3, 'lamCont', 0.4, 'lamCorr', 0.3)
```

Which creates a substruct `fuo.GraFTout` that contains the spatial components `fuo.GraFTout.spatial` and the temporal components `fuo.GraFTout.TimeTrace`.

#### B.6 Class methods

When initializing a multi data-set `uom`, either a list of `.mat` file-names, a cell array of datasets, or folder path can be provided. In the former two, a series of `uo` objects will be created to match the datasets or filenames input. In the last option, the class methods will determine all `.mat` files in the folder path and its sub-directories, and create a set of `uo` objects to access those files. The methods for these classes allow for loading, viewing, and performing basic processing steps. In particular, pre-processing steps such as de-noising, error detection and error correction.

**Data loading:** Basic tools for loading data are outlined in Table 2.

Table 2: Class methods for data access

| Function Name | Description |
|---|---|
| `uo()` | Main function to initialize and set up a fUS object |
| `makeGetBlockFunction()` | Creates a function that extracts a movie block from the full data |
| `makeGetMovieFunction()` | Creates a function that extracts the full movie data |
| `makeGetTraceFunction()` | Creates a function that extracts a single pixel time-trace from the full data |
| `isMatFileBased()` | Checks if a uo is mat-file based of if data is loaded to RAM |
| `ensureMask()` | Ensures that a mask has been provided to separate the in-brain pixels |
| `writeToAVI()` | Writes a dataset (pre- or post processing) to an AVI movie |
| `drawROI()` | Enables the user to draw a mask around the brain |

**Visualization:** Being able to visually sort through data is paramount to understanding the neural recordings. A number of class methods focus on this aspect, permitting the viewing of specific frames, movie snippets, example time-traces, lateral motion estimation through time, etc. These functions are detailed in Table 3.

Table 3: Class methods for visualization

| Function Name | Description |
|---|---|
| `displayMovie()` | Function to display a fUS movie using MovieSlider |
| `displayExampleFrame()` | Function to display an example frame from the fUS movie |
| `plotBurstErrors()` | Plot the burst errors to validate correct identification |
| `displayExampleTraces()` | function to display example time-traces from the fUS movie |
| `displayMotionError()` | Displays example motion errors visually |

**Motion and Error correction:** Code to remove artifacts from imaging are detailed in Table 4.

Table 4: Class methods for data cleaning

| Function Name | Description |
|---|---|
| `findBurstFrames()` | Function to identify frames with burst errors |
| `findMotionCorrectionError()` | Identifies rigid shift errors in fUS data |
| `ensureMotionErrsComputed()` | Ensures that the motion errors are computed for a given `uo` |
| `ensureMotionCorrection()` | Ensures that the movie for a given `uo` is motion corrected |
| `doesDenoiseExist()` | Checks if the denoised movie was already computed |
| `doesMotionCorrectedExist()` | Checks if the motion |
| `ensureDenoisedData()` | Checks if denoised data is available and if not denoises the data |
| `denoiseTracesWavelet` | Denoises a fUS movie one pixel at a time using wavelet denoising |
| `motionHypothesisTest()` | Tests the similarity of a time trace during large and small motion |
| `shuffleMotionHypothesisTest()` | Same as motionHypothesisTest but shuffling data |
| `getMotionVectors()` | Get the per-frame motion offsets |
| `getMotionCorrectedSize()` | Returns the correct size of the post-motion corrected movie |
| `compareMotionHypothesisTest()` | Compares the motion metric before and after motion correction |
| `computeBurstErrorInpainting()` | Fill in missing frames with bicubic interpolation |
| `correctResidualMotion()` | Shift frames to correct computed translational motion |

**Analysis:** Basic analysis tools, including PCA, GraFT and correlation computations can be computed using the functions in Table 5.

| Function Name | Description |
|---|---|
| `getBasicStats()` | Compute basic statistics per pixel and per movie |
| `getBaselineImage()` | Compute a baseline image to compare individual frames to |
| `calcBasicStats()` | Compute basic per movie statistics |
| `eventSTH()` | Compute an event-triggered average of the movie |
| `event2timeseries()` | Converts event times to a time-series spike train |
| `event2frame()` | Translate an event time to a movie frame |
| `ensureMedian()` | Ensure that the median image has been computed and is available |
| `fuGraFT()` | Apply GraFT to a fUS dataset |
| `fuPCA()` | Run PCA on a fUS dataset |
| `correlateMotionWithData()` | Correlate the motion estimates with individual pixel timetraces |
| `correlateToEvent()` | Compute each pixel's correlation with a cecurring event |
| `correlationMap()` | Compute a correlation map across the movie's spatial extent |
| `computeCorrsWithBaseline()` | Compute correlations of all pixels with a baseline time-trace |
| `clusterTraces()` | Cluster the pixels of the fUS data based on their time-traces |
| `alignTrials()` | Align all the trials based on event time-stamps |

Table 5: Class methods for basic analysis

**Extra functions:** The ultrasound class is supported by a number of additional functions in Table 6, and a number of external functions from other sources, detailed in Table 7.

Table 6: Support functions

| Function Name | Description |
|---|---|
| `applyMask()` | Applies a user-defined mask to a frame or video of the data. |
| `extractPixel()` | Extracts $N$ pixel time traces from the ultrasound movie |
| `findSigmoidPars()` | Fits a sigmoid to data |
| `fit2DLaplaceFun()` | Fits a 2D laplace function to data |
| `gaussfilt()` | Filters with a gaussian kernel |
| `greedyEND1D()` | Greedy solver for a 1D earth-mover's distance |
| `histogramDistance()` | Computes a distance between histograms |
| `lapFunc()` | Computes a laplace function in 2D given data and parameters |
| `lapLogFunc()` | Same as lapFunc but computation is in the log-domain |
| `motionFitSingleTest()` | Compare a single frame to a reference to identify translational motion |
| `plotAllAnats()` | Plots all the anatomical images for 4 different sessions |
| `plotWaveImages()` | Plots wavefronts of waves |
| `pmColorMap()` | Sets a color map with one color for pos. to a different color for neg. |
| `realignSingleFrame()` | Linearly translates a frame given a shift |
| `robustTwoSidedSTD()` | Computes a robust two-sided standard deviation |
| `selectFusiTraces()` | Selects a subset of time-traces (single pixels) from an ultrasound video |
| `softScale()` | Scale values of an array smoothly given a range to keep linear |

Table 7: External functions

| Function Name | Description |
|---|---|
| `AdvancedColormap()` | Function that creates additional color maps for plotting |
| `distinguishable_colors()` | Generates maximally distinguishable color sets for plotting |
| `halfSampleMode()` | Computes an approximate mode of a distribution |
| `robustSTD()` | Computes the robust standard deviation of a distribution |

**Multi-session data:** Functions for multi-session data are described in Table 8.

Table 8: Functions for the `uom` object class

| Function Name | Description |
|---|---|
| `uom()` | Main function for creating a uom object. |
| `ensureAllMasks()` | Makes sure all sub-objects have user-defined brain selections |
| `denoiseMulti()` | Denoise all uo datasets |
| `findMotionCorrectionErrorAll()` | Find translational motion for all datasets |
| `multiEventCorr()` | Correlate an event for all datasets |
| `multiMotionCorrect()` | Correct motion in all datasets |
| `multiMotionTest()` | Test motion correction in all datasets |
| `removeTrivialDatasets()` | Aux function to remove small datasets not worth analyzing |

### C Functional resolution

In this appendix, we summarize a method by which one can determine the functional resolution of ultrasonic imaging in rodents. We perform whisker stimulation experiments for this endeavour as the mapping of the somatosensory cortex to specific rodents whiskers is well established. We show that the whole whisker stimulation elicited consistent hemodynamic responses on somatosensory cortex barrel field (S1BF) and thalamic nuclei (VPL/VPM) marked by a significant increase in the cerebral blood volume (CBV) detected by functional ultrasound imaging. Moreover, we show that similar, but more localized fUSi responses were elicited by single whisker stimulation C2 and C3. The procedure we present in this appendix can be used a method for characterization and comparison of the functional resolution of custom made functional ultrasound imaging probes

#### C.1 Introduction

In order to characterize the functional resolution of ultrasound imaging, the paradigm of choice is whisker stimulation and the corresponding somatosensory activation. The somatotopic arrangement of the barrel fields upon whisker stimulation is stereotypical and has been previously characterized using calcium imaging, intrinsic optical imaging, multi-unit activity recording, two photon microscopy and functional magnetic resonance imaging. In this appendix, we complement this characterization with the functional ultrasound imaging technique that basically gives us advantages such as its ability to image at depth so one can image not only the activity in the somatosensory cortex, upon whisker stimulation, but also thalamic nuclei that are activated simulatenously and the functional relationships between the activated areas.

#### C.2 Methods

##### C.2.1 Surgery

In order to perform the functional ultrasound imaging, we need to perform a surgery to remove large parts of the skull. The craniotomy is necessary as the bone absorbs the ultrasound waves. In order to open an 6 x 12 mm craniotomy that goes as far lateral as possible to be able to image the somatosensory cortex, the following surgery was done:

The animal is anasthesized with 4% isoflurane. Dexamethasone 1 mg/kg IM is administered in order to reduce brain swelling and buprenorphine 0.05 mg/kg for pain management. The rat hair covering his head is then shaved using the trimmer. It is important to remove all the hair on the skin that will be the site of incision afterwards. The animal is then put in the ear bar. The shaved head is cleaned copiously with alternating betadine and alcohol. 0.15 ml Lidocaine / Norepinephrine mixture is injected into the head skin where the incision is planned. The skull overlying the brain region of interest is then exposed by making an incision (∼10-20 mm, rostral/caudal orientation) along the top of the head, through skin and muscle using a scalpel. Tissue is reflected back until the skull is exposed and held in place with forceps. The surface of the skull is thoroughly cleaned and scrubbed using a scraper and sterile saline. The cleaning and scrubbing continue until the surface of the skull is totally clean and devoid of signs of bleeding, blood vessels or clotting. A piece of the skin on the sides (∼2 mm thickness), parallel to the surface of the skull, is cut using sharp scissors. The muscles are also carefully dissected with a sharp scissor in order to be able to access the lateral sides of the skull. After dissecting the muscles, the skin is held open on both sides using sutures attached to hemostats (4 sutures and 4 hemostats are used). Using a marker, we mark the 6 mm craniotomy beginning from +1 AP till -4 AP. Just before beginning the craniotomy, we inject the animal with 2 g/kg mannitol in order to reduce brain swelling. Then using the piezosurgery apparatus with the micro-saw insert, the skull is slowly removed. The drilling has to be very slow and the skull has to be continuously irrigated with sterile saline. The size of the craniotomy done is 6 mm X 12 mm. Using piezosurgery instead of the conventional burr is crucial as it avoids touching the dura and injuring the mid-saggital vein while doing the craniotomy. When we feel that the skull is almost removed, we then use a small spatula in order to separate the dura from the skull very carefully. The brain is left to rest then the ultrasound gel is added and the imaging begins.

##### C.2.2 Stimulation setup

We used the verasonics ultrasound probe L22-14 which covers a width of 12,8mm (pitch 0,100 mm). The probe has 128 channels.

We used a fast sequence for functional imaging that we adapted from the $\mu$Doppler sequence described with the following parameters: five angles ($-6\,\text{deg}, -3\,\text{deg}, 0\,\text{deg}, 3\,\text{deg}, 6\,\text{deg}$) averaged three times, 7.5 kHz firing frequency, 500 Hz frame rate, 200 images, 0.4 s acquisition time, and impulse of two cycles at 15 MHz (Urban et al., 2015).

As a final result, we obtained an image I(x, z) proportional to the CBV signal that we termed 'CBV image' for simplification. For detection of hemodynamic variations, we captured CBV images every 0.5 s during the entire whisker stimulation period to obtain a time-dependent signal of CBV in each pixel I(x, z, t)(Mace et al., 2013)

##### C.2.3 Pre-processing

The CBV measured by fUSi signal includes stimulus specific and non-specific components. Low-frequency components ($< 0.02$Hz) are typically related to physiological system wide variation in the hemodynamics. We call this low-frequency component as baseline. The fUSi signals for each pixel was high-pass filtered at 0.02 Hz using a Butter filter of order 2. The high pass filtered signal was then compared to the baseline signal and the percentage change corresponds to CBV percentage change.

##### C.2.4 Region-of-interest (ROI)

To localize the brain areas with fUSi signals that were highly correlated with stimuli, we calculated the cross-correlation between the high pass filtered fUSi signal and the smoothed stimulus design. The stimulus design is a signal that is one when there is stimulus and zero otherwise. To take into account the low-pass filtering of the hemodynamic response, we convolved the stimulus design with a Gaussian kernel (gaussfilt) with standard deviation = 2.5 s. The maximum cross-correlation value between the high pass filtered fUSi and the smoothed stimulus design was used to localize the ROI in the brain map. We only considered the cross-correlation values in which the fUSi signal was delayed with respect to the stimuli. The ROI were the brain areas whose maximum cross-correlation values were above the 95 percentile of all the correlations in the brain section and it was located in the S1BF or ventral posterolateral nucleus (VPL) / ventral posteromedial nucleus (VPM) of the thalamus. Although, in most cases we only observed high correlation in S1BF and VPL/VPM, we also observed sometimes some correlation in auditory cortex (probably due to the sound of stimulation) and non-barrel S1 (probably because we stimulates the surrounding skin during our stimulation).

##### C.2.5 Registration

To anatomically localize the ROIs, we used the cortical surface and the contours of the hippocampus as landmarks to compare with the paxinos rat atlas(Paxinos and Watson, 1986). We also used the location with respect to the Bregma estimated using the stereotaxis. The comparison was done manually using Adobe Illustrator and overlapping the images. For more precise registration, one can use the same procedure we used in the main text of the manuscript that involves brain clearing and registration to a common rat atlas.

#### C.3 Results

To show that there are fUSi signals with dynamics that are correlated with stimuli, we manually stimulated multiple whiskers simultaneously using a single stroke lasting approximately 3 s while imaging simultaneously. We predicted that the barrel cortex and the corresponding barreloids in thalamus are the brain areas where the fUSi signal dynamics is most correlated with the whisker stimulation dynamics. We first calculated the maximum cross-correlation value between the fUSi signals for every pixel with the signal representing the stimuli (0 when there is no stimuli and 1 when there is). Then, we obtained the highest 5 percentile correlation values of the entire section. Figure 1A shows the regions of interest (ROI) - areas that have correlation values with stimuli in the top 5 percentile and that overlaps with S1BF and VPL/VPM at Bregma -3. Figure 1B shows the ROIs with its correlation values. Figure 1C shows the same ROIs with its delay values. The delay in S1BF is significantly larger than in VPL/VM (p¡0.05, t-test). We also calculated the ROI for coronal sections Bregma = -2 to -3 at 200 $\mu$m intervals, showing that the whisker stimulation successfully elicited highly correlated activities in S1Bf and/or VPL/VPM for all sections (Figure 2).

To further understand the fUSi signal response to whisker stimulation, we calculated the average fUSi activity in S1BF ROI and VPL/VPM ROI and aligned it to the the onset of the whisker stimulation. Figure 3A shows four exemplars of fUSi signals in S1BF ROI after slow frequency component (¡0.02 Hz) was subtracted. The slow frequency component is related to the systemic hemodynamic response independent of the stimuli. There is an increase in the fUSi signal approximately 6s from the onset of whisker simulation, consistent with delay map Figure 1C. Figure 3B shows that the mean fSUi signal for 24 trials (black lines) significantly increases after the onset of whisker simulation. To show that the fUSi signal in S1BF was specific to the ROI, we computed the average fUSi signal of the contralateral (ipsilateral to the stimuli) S1BF, symmetric to the S1BF ROI. Figure 3B, blue line, shows that there is no significant fUSi signal response after whisker stimulation. Similarly,the average fUSi signal of the ipsilateral M1 (contralateral to the stimuli) aligned to the stimuli onset was not significantly different from baseline (Figure 3B, yellow line). Figure 3C shows four exemplars of fUSi signals in VPL/VPM ROI after the slow frequency component ($< 0.02$ Hz) was subtracted. There is an increase in the fUSi signal approximately 3s from the onset of whisker simulation, consistent with delay map Figure 1C. Figure 3D shows that the mean fUSi signal for 24 trials (black lines) significantly increases after the onset of whisker simulation. Figure 3A,C suggest that the fUSi signal response profile for S1BF ROI was more variable than the profile for VPL/VPM ROI. To verify this observation, we plotted all the signals together. Figures 3E,F show that, indeed, the response profile in VPL/VPM is more stereotyped than the response profile in S1BF.

We next asked whether fUSi signals were able to detect hemodynamic response to single whisker stimulation. Each stimuli was a sequence of three backward-forward movement, lasting in total approximately 3s, of C3 and C2 whiskers. Figure 4A shows the S1BF ROI at Bregma -2 corresponding to C3 whisker stimulation. As expected the size of the ROI is smaller compared to the multiple whisker stimulation in Figure 1. The size of the ROI diameter parallel to the brain surface is $315\mu m$, which is compatible to the expected size of a single barrel in S1BF(Egger et al., 2012). The distance to the cortical surface is approximately 900 $\mu$m, which is also compatible with the strongest response expected in the cortical layer IV (between 500 to 1200 $\mu$m), although we cannot confirm the layer specificity. Figure 4B shows the registration using an approximate coordinate in the atlas confirming the S1BF response. Figure 4C shows three single trial exemplars of the fUSi signal responses. Each response is the average signal in S1BF ROI in which the slow frequency component ($< 0.02$ Hz) was subtracted. Figure 4D shows the average fUSi signal for 24 trials. Although the ROI clearly shows the response compatible to a response in a single barrel, the 95% threshold can be rather arbitrary. To better understand the localization property of fUSi signal, we show in Figure 4E-F the decay of correlation (maximum cross-correlation value between the fUSi signal and stimuli) as a function of distance from the location with peak correlation. The decay of correlation in all directions show a sharp decrease consistent with a well localized signal.

We repeated the analysis for C2 whisker stimulation. Figure 5A shows the S1BF ROI in coronal section Bregma -2.2 with the diameter parallel to cortical surface (393 $\mu$m) compatible with the size of a single C2 barrel in S1BF (Egger et al., 2012) and distance from surface (622 $\mu$m) compatible layer IV activation. Figure 5B confirms the anatomical location of S1BF ROI. We show in Figure 5C three single trial exemplars of fUSi signal responses in S1BF ROI. Figure 5D shows a clear response profile for the average fUSi response for 24 trials. To demonstrate that the C3 and C2 whisker stimulation ROI are distinct, we overlapped both maps in Figure 5D. Clearly, the location are different not only in depth and lateral position, it is also on different coronal sections. The more caudal position of C2 is also compatible with the known relationship between C3 and C2 barrels. Furthermore, the distance between the areas with the largest correlation in C3 and C2 ROIs is approximately 900 $\mu$m, which is compatible with the distance between C3 and C2 barrel centers(Egger et al., 2012).

We also wanted to verify if the fUSi signal is able to detect hemodynamic response in thalamus with a single whisker stimulation. Figure 6A shows the VPL ROI at Bregma - 2.4 for C2 whisker stimulation. We confirm the anatomical location by registering in the atlas (Figure 6B). Figure 6 shows three single trial exemplars for the fUSi response in VPL ROI. Figure 6 shows the clear fUSi response for the average of 24 trials.

#### C.4 Conclusion

Using the whisker system as a canonical model to determine the functional resolution of functional ultrasound imaging, our analysis has revealed that, in terms of spatial Resolution, one can go down to at least 300 $\mu$m. Moreover, the response to whisker stimulation can be observed in single trials. 24 trials were enough to see the effect very consistently. We can calculate anatomical distance between different functional brain areas involved upon whisker stimulation.The spatial correlation is very precise as we have seen from the decay of the spatial correlation in the single whisker experiments.We were able to differentiate the functional activation between two single whiskers C2 and C3. Such functional studies are crucial to determine and compare the functional resolution of different ultrasonic probes.

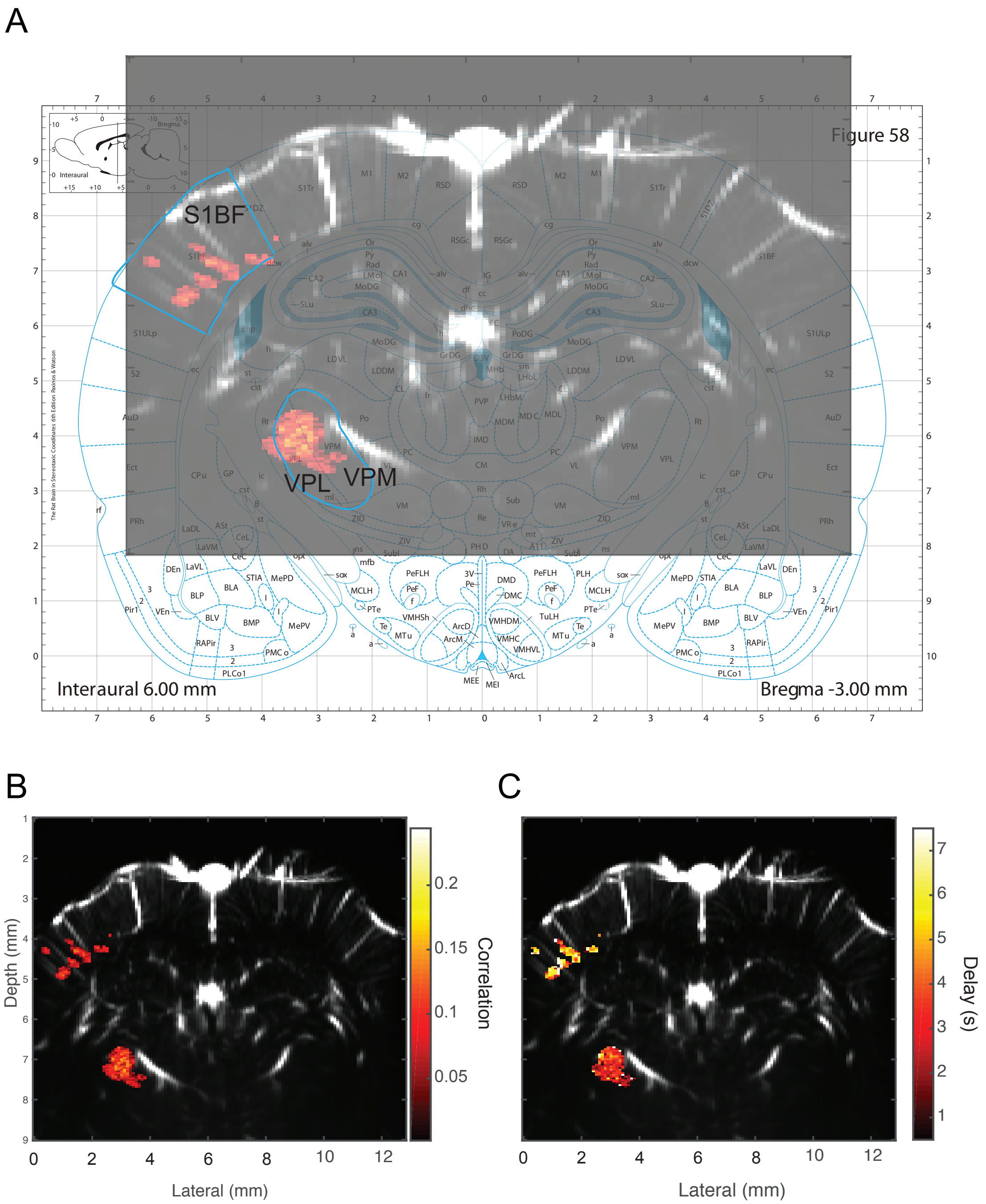

Figure 1: Activation of S1BF and VPL/VPM with full whisker stimulation.(A) ROI - highly correlated areas (red regions) - in the S1BF and VPL/VPM areas of the rat brain. Each pixel size is $48\mu m \times 100\mu m (depth \times width)$(B) Maximal cross-correlation values for the whisker related fUSi signals in the S1BF and VPL/VPM areas. The values are shown only for brain areas whose correlation values exceed the 95 percentile correlation values for the signals from entire section. (C) Delay values for the whisker related fUSi signals in the S1BF and VPL/VPM areas. The delays are obtained as the time point of the maximal cross-correlation value with respect to the stimuli. The delays are $\approx$ 3s for VPL/VPM and $\approx$ 6s for S1BF.

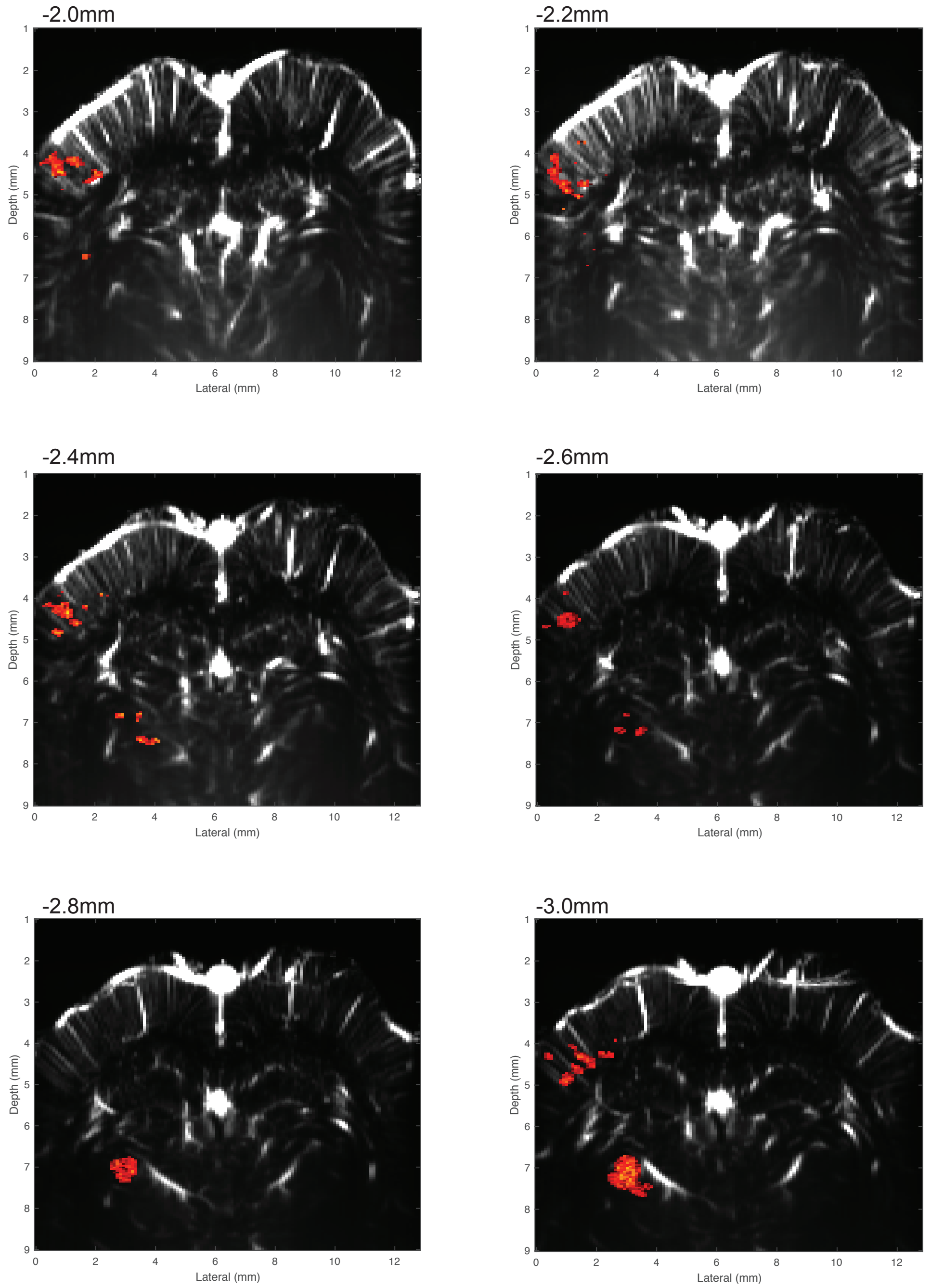

Figure 2: Activation of S1BF and VPL/VPM in different coronal sections. Regions highlighted in red indicate ROI - regions with fUSi signals that are highly correlated (above 95 percentile value of the correlation of the signals in the entire section) with the stimuli.

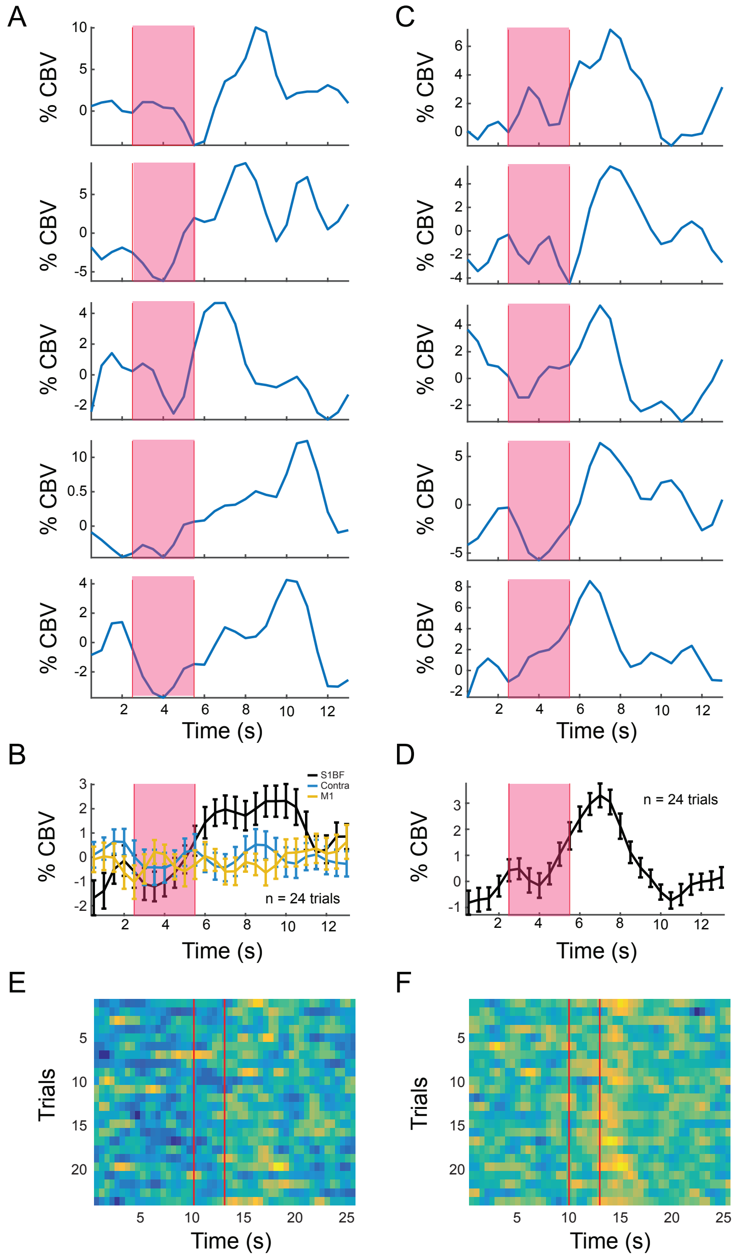

Figure 3: fUSi signals in S1BF and VPL/VPM. (A) Single trial exemplars for average fUSi signals in S1BF ROI. x-axis shows time in seconds; y-axis shows percentage change from baseline. The baseline is the low frequency component of the signal, below 0.02 Hz. The red regions indicate the duration of the full whisker stimulation. (B) Average fUSi signal for 24 trials. Error bars indicate one standard error of the mean. Black line shows the average for S1BF ROI. Blue and yellow lines show the average for symmetrically chosen contra lateral S1BF area and arbitrarily chosen M1 area, respectively. (C) Single trial exemplars for average fUSi signals in VPL/VPM ROI. Convention is the same as in (A). (D) Average fUSi signal in VPL/VPM ROI for 24 trials. Convention is the same as in (B). (E) Plot showing the aligned average fUSi signals in S1BF ROI for all 24 trials. Warmer color indicates larger values. (F) Plot showing the aligned average fUSi signals in VPL/VPM ROI for all 24 trials.

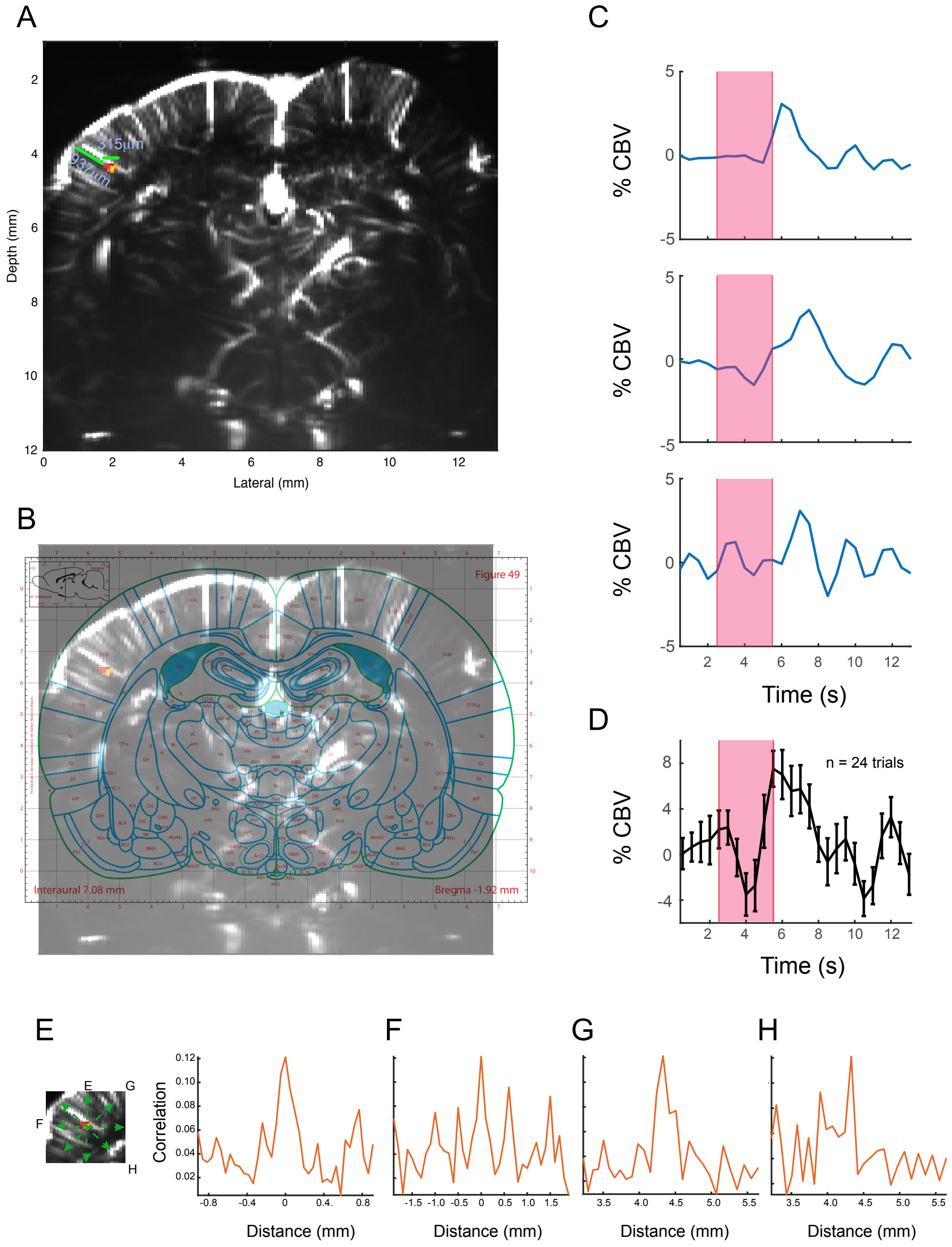

Figure 4: Activation of S1BF by C3 whisker stimulation. (A) Activation map showing the brain region with fUSi signal that is highly correlated with C3 whisker stimulation in S1BF (red/yellow). The diameter parallel to surface has $315\mu m$ and the distance from the surface to the most activated region is 935 $\mu$m. (B) Registration of the activation map into rat atlas showing the S1BF location of the activated region. (C) Single trial exemplars for average fUSi signals in S1BF ROI. Convention is the same as in Figure 3A. (D) Average fUSi signal for 24 trials. Convention is the same as in Figure 3B. (E-H) Spatial correlation plot for different directions showing the sharp decay of maximum cross-correlation values as a function of distance from the most correlated area.

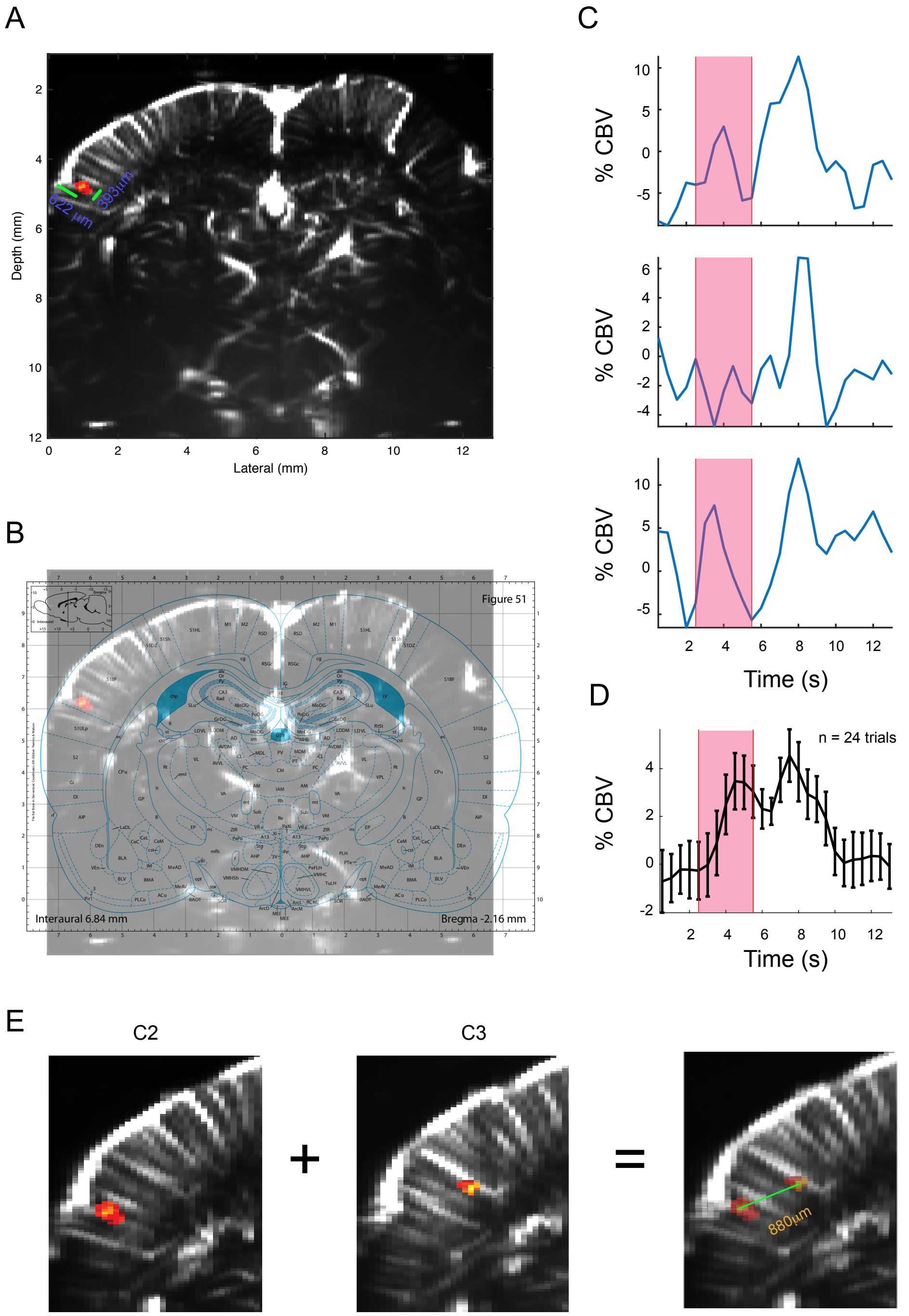

Figure 5: Activation of S1BF by C2 whisker stimulation. (A) Activation map showing the brain region with fUSi signal that is highly correlated with C2 whisker stimulation in S1BF (red/yellow). The diameter parallel to surface has 393 $\mu$m and the distance from the surface to the most activated region is 622 $\mu$m. (B) Registration of the activation map into rat atlas showing the S1BF location of the activated region. (C) Single trial exemplars for average fUSi signals in S1BF ROI. Convention is the same as in Figure 3A. (D) Average fUSi signal for 24 trials. Convention is the same as in Figure 3B. (E) Map comparing the location of S1BF areas activated by C3 and C2 whisker stimulation. The distance from the two most correlated regions in each ROI is 880 $\mu$m.

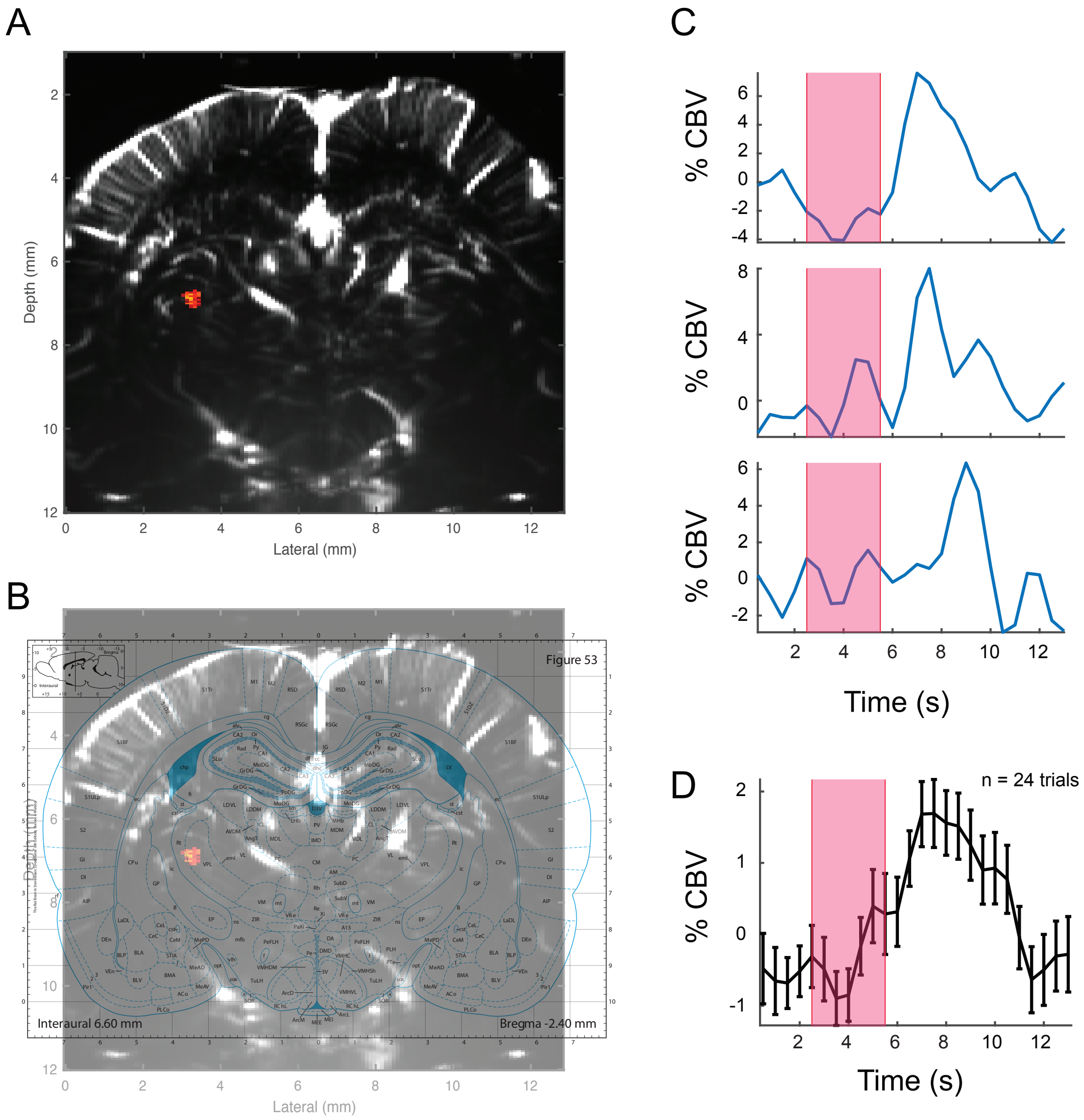

Figure 6: Activation of VPL by C2 whisker stimulation. (A) Activation map showing the brain region with fUSi signal that is highly correlated with C2 whisker stimulation in S1BF (red/yellow).(B) Registration of the activation map into rat atlas showing the VPL location of the activated region. (C) Single trial exemplars for average fUSi signals in S1BF ROI. Convention is the same as in Figure 3A. (D) Average fUSi signal for 24 trials. Convention is the same as in Figure 3B.
